# Improved genetic prediction of complex traits from individual-level data or summary statistics

**DOI:** 10.1101/2020.08.24.265280

**Authors:** Qianqian Zhang, Florian Privé, Bjarni Vilhjálmsson, Doug Speed

## Abstract

Most existing tools for constructing genetic prediction models begin with the assumption that all genetic variants contribute equally towards the phenotype. However, this represents a suboptimal model for how heritability is distributed across the genome. Therefore, we develop prediction tools that allow the user to specify the heritability model. We compare individual-level data prediction tools using 14 UK Biobank phenotypes; our new tool LDAK-Bolt-Predict outperforms the existing tools Lasso, BLUP, Bolt-LMM and BayesR for all 14 phenotypes. We compare summary statistic prediction tools using 225 UK Biobank phenotypes; our new tool LDAK-BayesR-SS outperforms the existing tools lassosum, sBLUP, LDpred and SBayesR for 223 of the 225 phenotypes. The increase in prediction accuracy from improving the heritability model tends to be substantial. For example, when using LDAK-Bolt-Predict, the proportion of phenotypic variance explained increased by on average 14% (range 5-29%), equivalent to increasing the sample size by a quarter.

## INTRODUCTION

There is a great demand for more accurate genetic prediction models of complex traits. Better models will, for example, improve our ability to investigate genetic architecture, detect genetic overlap between traits and search for gene-environment interactions.^1,2^ They will also enable more widespread use of precision medicine, for example, by enabling us to better identify subgroups of individuals with elevated risk of developing a particular disease, or those with lowest chance of responding to a particular medication.^3–7^

Many complex traits have high SNP heritability, which justifies the use of genome-wide, linear, SNP-based prediction models.^8,9^ The resulting predictions are called polygenic risk scores (PRS). They take the form **P** = **X**_1_ *β*_1_ + **X**_2_ *β*_2_ + …+ **X**_*m*_ *β*_*m*,_ where *m* is the total number of SNPs, while **X**_*j*_ and *β*_*j*_ denote, respectively, the genotypes and estimated effect size for SNP *j*. Tools for constructing PRS differ in how they estimate the SNP effect sizes. The simplest way to construct a PRS is using effect size estimates from single-predictor regression (classical PRS). However, it is generally better to use an advanced prediction tool that estimates effect sizes using a multi-SNP regression model.^10–14^

Advanced prediction tools start by making prior assumptions regarding how SNPs contribute towards the phenotype. These assumptions include specifying a heritability model, which describes how E[*h*^2^_*j*_], the expected heritability contributed by each SNP, varies across the genome.^15^ Almost all existing advanced prediction tools automatically assume that E[*h*^2^_*j*_] is constant. We refer to this as the GCTA Model, because it a core assumption of the software GCTA.^8^ In particular, the GCTA Model is assumed by any prediction tool that uses a multi-SNP regression model and assigns the same penalty or prior distribution to standardized SNP effect sizes.^9,16^ However, the GCTA Model is suboptimal. Recently, we provided a method for comparing different heritability models using summary statistics from genome-wide association studies.^17^ Across tens of complex traits, the model that fit real data best was the BLD-LDAK Model, in which E[*h*^2^_*j*_] depends on minor allele frequency (MAF), local levels of linkage disequilibrium and functional annotations.

In this paper, we construct PRS for a variety of complex traits using eight new prediction tools. The main difference between these and existing tools is that they allow the user to specify the heritability model. We show that for all eight tools, the accuracy of the PRS improves when we switch from the GCTA Model to the BLD-LDAK Model. When individual-level genotype and phenotype data are available, we recommend using our new tool LDAK-Bolt-Predict (a generalized version of the prediction tool contained within the existing software Bolt-LMM^18^). With access only to summary statistics and a reference panel, we recommend using our new tool LDAK-BayesR-SS (a generalized version of the existing prediction tool SBayesR^14^). Both tools are available in our software LDAK^15^ (www.ldak.org).

## RESULTS

### Overview of Methods

Figure 1a classifies our eight new prediction tools based on the form of the prior distribution they assign to SNP effect sizes. Our four individual-level tools, big_spLinReg, LDAK-Ridge-Predict, LDAK-Bolt-Predict and LDAK-BayesR-Predict, use the same prior distribution forms as the existing individual-level data tools Lasso,^16^ BLUP,^19^ Bolt-LMM^18^ and BayesR,^11^ respectively. Our four new summary statistic tools, LDAK-Lasso-SS, LDAK-Ridge-SS, LDAK-Bolt-SS and LDAK-BayesR-SS, use the same prior distribution forms as the existing summary statistic tools lassosum,^13^ sBLUP,^20^ LDpred^12^ and SBayesR,^14^ respectively. Figure 1b illustrates how our new tools incorporate alternative heritability models by allowing the parameters of the effect size prior distribution to vary across SNPs. We provide full details of our new tools in Methods, and scripts for repeating our analyses in Supplementary Note 1.

**Fig. 1.**
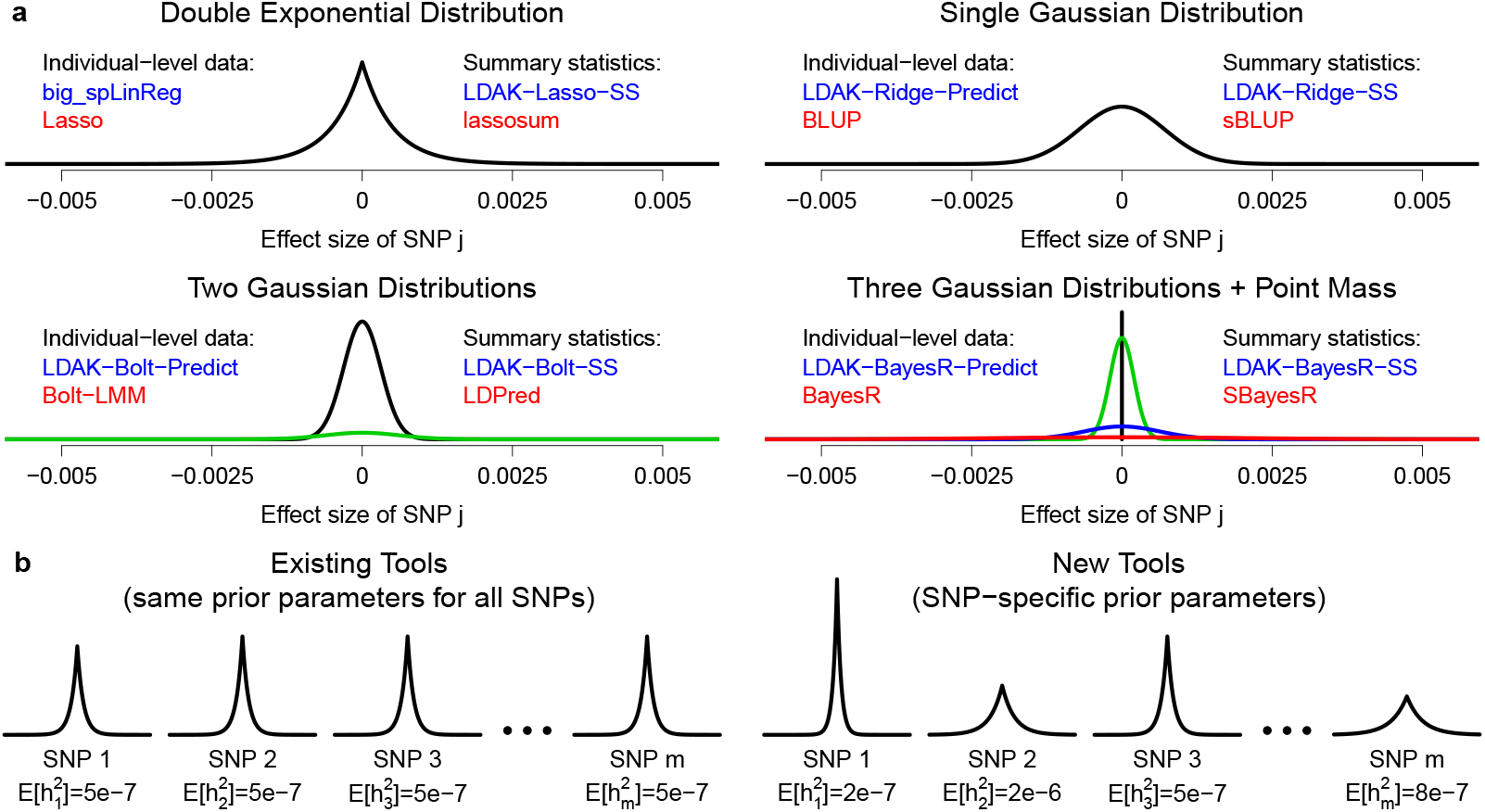
Prior distributions for SNP effect sizes. **a** We divide prediction tools based on the form of the prior distribution they assign to SNP effect sizes, and whether they use individual-level data or summary statistics. For each of our eight new tools (names in blue), there is an existing tool that uses the same prior distribution form (names in red). **b** Having selected the form of the effect size prior distribution, most existing prediction tools use the same parameters for each SNP. Our new tools, by contrast, use SNP-specific prior distribution parameters. To illustrate this difference, we consider lasso-based prediction tools that assign a double exponential prior distribution to standardized SNP effect sizes. While existing tools might, for example, set the variance of the prior distribution to 5e-7 (so that E[h^2^_j_]=5e-7 for all SNPs), our new tools instead let the variance vary across the genome (allowing E[h^2^_j_] to be set according to the chosen heritability model).

In total, we construct PRS for 225 phenotypes from the UK Biobank^21,22^ (Supplementary Data 1). When using individual-level prediction tools, we restrict to the 14 phenotypes for which we have access to individual-level data. Of these, eight are continuous (body mass index, forced vital capacity, height, impedance, neuroticism score, pulse rate, reaction time and systolic blood pressure), four are binary (college education, ever smoked, hypertension and snorer), and two are ordinal (difficulty falling asleep and preference for evenings). For each phenotype, we have 220,000 distantly-related (pairwise allelic correlations <0.03125), white British individuals, recorded for 628,694 high-quality (information score >0.9), common (MAF >0.01), autosomal, directly-genotyped SNPs. When constructing PRS, we use 200,000 individuals as training samples, and the remaining 20,000 individuals as test samples. When we require a reference panel, we use the genotypes of 20,000 individuals picked at random from the 200,000 training samples. We measure the accuracy of a PRS via *R*^2^, the squared correlation between observed and predicted phenotypes across the 20,000 test samples, and estimate the s.d. of *R*^2^ via jackknifing. For a given phenotype, *R*^2^ is upper-bounded by *h*^2^_SNP_, the SNP heritability, estimates of which range from 0.07 to 0.61 (Supplementary Table 1). When using summary statistic prediction tools, we construct PRS for all 225 phenotypes, using results released by the Neale Lab. These results come from association studies with average sample size 285k (range 35-361k), and the average *h*^2^_SNP_ is 0.22 (range 0.07-0.63).

We consider three different heritability models: the GCTA Model assumes E[*h*^2^_*j*_] is constant, the LDAK-Thin Model allows E[*h*^2^_*j*_] to vary based on the MAF of SNP j, while the BLD-LDAK Model allows E[*h*^2^_*j*_] to vary based on the MAF of SNP j, local levels of linkage disequilibrium and functional annotations.^17^ Our previous work compared heritability models based on how well they fit real data.^17^ Specifically, we measured their performance via the Akaike Information Criterion^23^ (AIC), equal to 2*K* - 2logl, where *K* is the number of parameters in the heritability model and logl is the approximate log likelihood (lower AIC is better). Across the 12 models we considered, AIC was lowest for the BLD-LDAK Model, highest for the the GCTA Model, and intermediate for the LDAK-Thin Model (we reproduce these results in Supplementary Table 2).

Supplementary Fig. 1 shows that when run assuming the GCTA Model, each of our new prediction tools performs at least as well as the corresponding existing tool. For some pairs of tools, the results are almost identical. For example, the PRS constructed using LDAK-Bolt-Predict and LDAK-BayesR-SS assuming the GCTA Model have similar accuracy to those constructed using Bolt-LMM and SBayesR, respectively.

However, for other pairs, our tools are superior. For example, the PRS constructed using LDAK-Lasso-SS and LDAK-Ridge-SS assuming the GCTA Model tend to be more accurate than those from lassosum and sBLUP, respectively. We explain the algorithmic innovations that lead to these improvements in Supplementary Note 2. As the aim of this paper is to demonstrate the impact on prediction accuracy of improving the heritability model (not due to algorithmic innovations), for the analyses below, we always use our new tools.

### Performance of individual-level data prediction tools

First we use our four new individual-level data tools to construct PRS for the first 14 UK Biobank phenotypes. When using all 200,000 training samples, the tools take approximately 4 hours (LDAK-Ridge-Predict), 20 hours (LDAK-Bolt-Predict) or 50 hours (big_spLinReg and LDAK-BayesR-Predict), and require 35Gb memory (note that for big_spLinReg, LDAK-Bolt-Predict and LDAK-BayesR-Predict, the runtimes can be reduced substantially by using multiple CPUs).

Figure 2 and Supplementary Table 3 show that the accuracy of PRS always increases when we replace the GCTA Model with either the LDAK-Thin or BLD-LDAK Model (i.e., for all four tools and for all 14 phenotypes). For our recommended tool, LDAK-Bolt-Predict, replacing the GCTA Model with the LDAK-Thin Model increases *R*^2^ by on average 9% (s.d. 2%), while replacing the GCTA Model with the BLD-LDAK Model increases *R*^2^ by on average 14% (s.d. 2%). Moreover, when run assuming the BLD-LDAK Model, LDAK-Bolt-Predict outperforms our implementations of the existing tools Lasso, BLUP, Bolt-LMM and BayesR for all 14 phenotypes. We note that the performances of LDAK-Bolt-Predict and LDAK-Bayes-Predict are very similar. For example, when run assuming the BLD-LDAK Model, the tools have average *R*^2^ 0.080 and 0.081, respectively (s.d.s 0.001), and each tool produces the most accurate PRS for seven of the 14 phenotypes. Therefore, our decision to recommend LDAK-Bolt-Predict simply reflects its faster runtime.

**Fig. 2.**
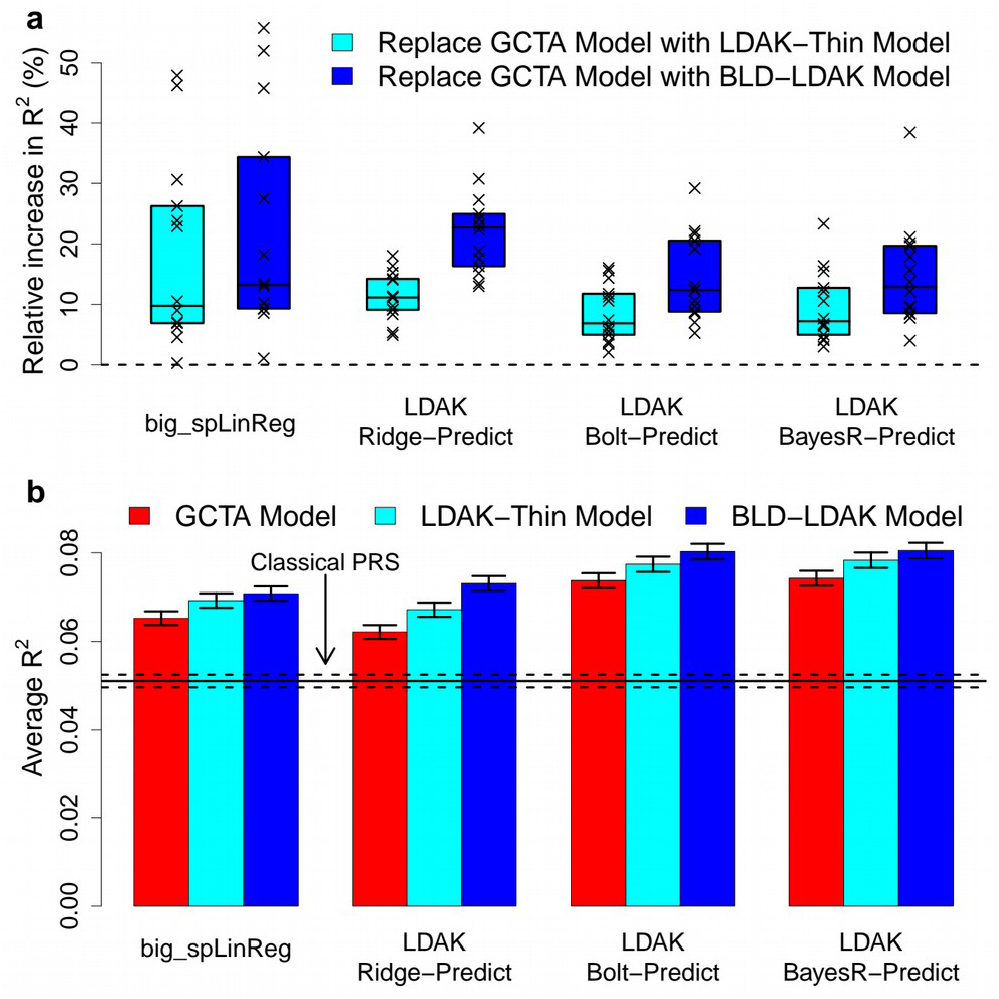
Impact of changing the heritability model when using individual-level data. **a** We use our four new individual-level data prediction tools to construct PRS for the first 14 UK Biobank phenotypes (using all 200,000 training samples). We measure the accuracy of each PRS via R^2^, the squared correlation between observed and predicted phenotypes across 20,000 test samples. Points report the percentage increase in R ^2^ for individual phenotypes when each tool is switched from assuming the GCTA Model to either the LDAK-Thin or BLD-LDAK Model (boxes mark the median and inter-quartile range across the 14 phenotypes). **b** For the same analysis as **a**, bars report R^2^ averaged across the 14 phenotypes (vertical segments mark 95% confidence intervals). Colors indicate the assumed heritability model, while blocks indicate the prediction tool. The horizontal lines mark average R^2^ for classical PRS and a 95% confidence interval

Figure 3 and Supplementary Table 1 show how the accuracy of PRS constructed using LDAK-Bolt-Predict varies with the number of training samples. We find that the increase we observed when we switched from the GCTA Model to the BLD-LDAK Model is equivalent to increasing the number of training samples by about 24%. The ratio *R*^2^/*h*^2^_SNP_ indicates the accuracy of a PRS relative to the maximum possible accuracy. When we use 200,000 training samples, the PRS achieve between 13% (difficulty falling asleep) and 62% (height) of their potential. The lines of best fit suggest that if we had individual-level data for 400,000 samples, the PRS would explain between 23% and 78% of SNP heritability.

**Fig. 3.**
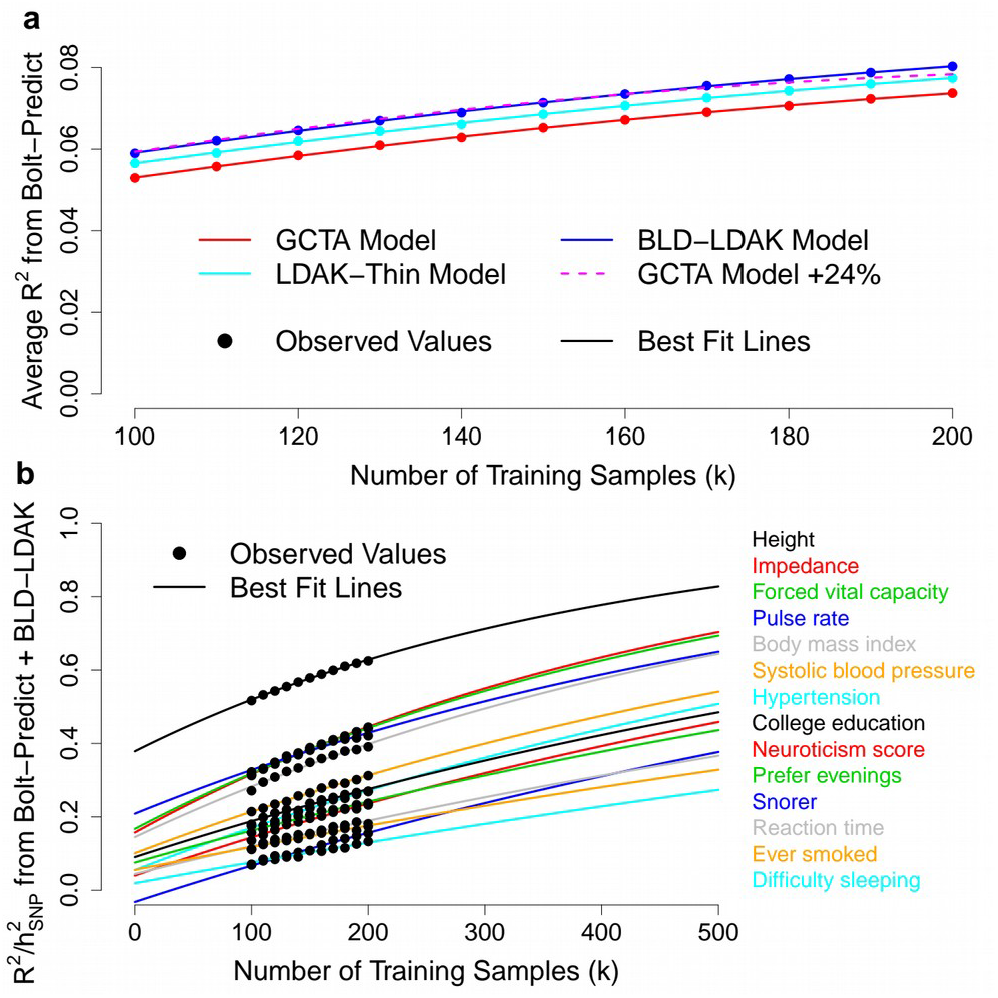
Dependency of prediction accuracy on sample size. **a** We use LDAK-Bolt-Predict to construct PRS for the first 14 phenotypes, varying n, the number of training samples, between 100,000 and 200,000. We measure the accuracy of each PRS via R^2^, the squared correlation between observed and predicted phenotypes across 20,000 test samples. Points report R^2^ averaged across the 14 phenotypes; colors indicate the assumed heritability model. The lines of best fit are obtained by regressing average R^2^ on a+bn+cn^2^; for the GCTA Model, we use the best fit line to predict average R^2^ if the sample size was 24% higher than specified (dashed line). **b** For the same analysis as **a**, points report R^2^/h^2^_SNP_ for PRS constructed assuming the BLD-LDAK Model, where h^2^_SNP_ is the estimated SNP heritability (the maximum possible R^2^). The lines of best fit are obtained by regressing R^2^/h^2^_SNP_ on 1-exp(a+bn).

### Performance of summary statistic prediction tools

Now we use our four new summary statistic tools to construct PRS for all 225 UK Biobank phenotypes. To construct each PRS takes under two hours (regardless of which tool we use) and requires less than 10Gb memory. Supplementary Fig. 2 and Supplementary Data 1 show that switching from the GCTA Model to the LDAK-Thin Model increases *R*^2^ for between 217 and 225 phenotypes (depending on tool), while switching from the GCTA Model to the BLD-LDAK Model increases *R*^2^ for between 223 and 225 phenotypes. LDAK-BayesR-SS has the highest average *R*^2^ of the four prediction tools, and produces the most accurate PRS for 137 of the 225 phenotypes.

Figure 4 shows that when run assuming the BLD-LDAK Model, LDAK-BayesR-SS outperforms our implementations of the existing tools lassosum, sBLUP, LDpred and SBayesR for 223 of the 225 phenotypes. Compared to the best existing tool, the average increase in *R*^2^ is 14% (s.d. 1%). Consistent with simulations (Supplementary Figs. 3 & 4), we find that the increase tends to be higher for phenotypes with lower *R*^2^. Nonetheless, the average increase remains substantial and significant (*P*<1e-16 from a one-sided Wald Test) if we consider only the 106 phenotypes with *R*^2^<0.05, only the 51 phenotypes with 0.05<*R*^2^<0.1, or only the 68 phenotypes with *R*^2^>0.1.

**Fig. 4.**
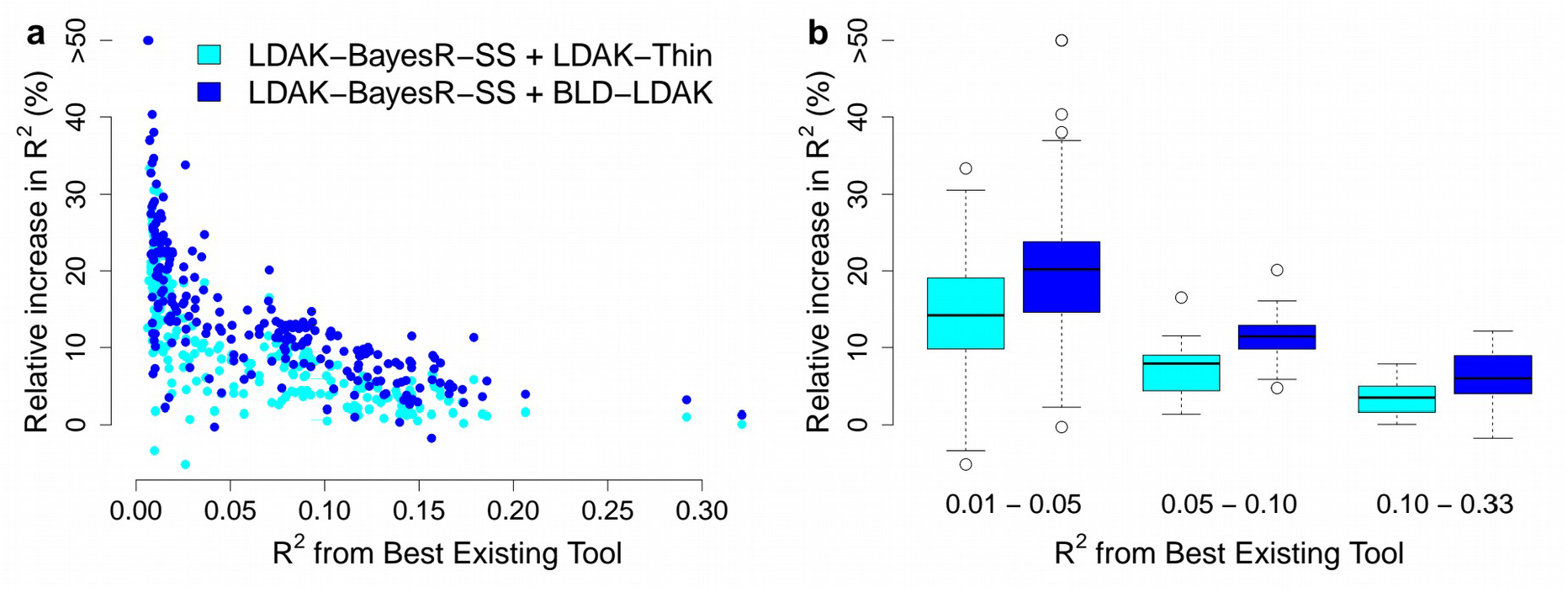
Impact of changing the heritability model when using summary statistics. **a** For each of the 225 phenotypes, we compare PRS constructed using LDAK-BayesR-SS assuming either the LDAK-Thin or BLD-LDAK Model, to PRS constructed using our implementations of the existing tools lassosum, sBLUP, LDpred and SBayesR. We measure the accuracy of PRS via R^2^, the squared correlation between observed and predicted phenotypes. The x-axis reports highest R^2^ across the four existing tools, while the y-axis reports the percentage increase in R^2^ if instead of using the existing tool with highest R^2^, we use LDAK-BayesR-SS assuming either the LDAK-Thin or BLD-LDAK Model (improvements above 50% are truncated). **b** The same as **a**, except phenotypes are grouped based on highest R^2^ across the four existing tools: 0.01-0.05 (106 phenotypes), 0.05-0.10 (51 phenotypes) or 0.10-0.33 (68 phenotypes). Boxes mark the median increase in R^2^ and the inter-quartile range.

### Additional Analyses

For our main analyses, we measured the accuracy of PRS using *R*^2^. Supplementary Fig. 5 shows that improving the heritability model improves accuracy if we instead measure mean absolute error or (for the binary phenotypes) area under the curve. For our main analyses, we used only directly-genotyped SNPs. Supplementary Fig. 6 shows that improving the heritability model also improves the accuracy of PRS when we increase the number of SNPs from 629,000 to 7.5M by including imputed genotypes.

For Supplementary Fig. 7 and Supplementary Table 4, we consider eight diseases: asthma, atrial fibrillation, breast cancer, inflammatory bowel disease, prostate cancer, rheumatoid arthritis, schizophrenia and type 2 diabetes. For each disease, we construct PRS using summary statistics from published studies (average sample size 117,000, range 35,000-215,000) that did not include UK Biobank data,^24–31^ then test them using UK Biobank data. Again, we find that for all phenotypes, the accuracy of PRS improves when we replace the GCTA Model with the LDAK-Thin or BLD-LDAK Model. This indicates that the improvements we observed in the main analyses are not an artifact of genotyping errors (as were this the case, we would expect the improvements to disappear when using training and test individuals that have been genotyped independently).

For our main analyses, we used white British individuals from the UK Biobank both to train and test the PRS. For Supplementary Fig. 8 and Supplementary Table 5, we instead test the PRS using UK Biobank individuals of South Asian, African and East Asian ancestry. While absolute accuracy is substantially lower, it remains that PRS constructed assuming the LDAK-Thin or BLD-LDAK Models are more accurate than those constructed assuming the GCTA Model. This indicates that the improvements we observed in the main analyses are not due to population structure (as were this the case, we would expect prediction models constructed assuming the LDAK-Thin or BLD-LDAK Models to perform worse across populations than those constructed assuming the GCTA Model).

## DISCUSSION

Most existing prediction tools start with the assumption that each SNP contributes equal heritability.^9^ We have instead developed tools that allow the user to specify more realistic heritability models, and shown how these enable the creation of substantially more accurate PRS. Of our eight new tools, we recommend using LDAK-Bolt-Predict when analyzing individual-level data, and LDAK-BayesR-SS when analyzing summary statistics (in both cases, we advise using the tools assuming the BLD-LDAK Model).

When using LDAK-Bolt-Predict, the average increase in *R*^2^ due to changing from the GCTA Model to the BLD-LDAK Model was 14% (s.d. 2%). We showed that this increase is equivalent to increasing the sample size by about a quarter. To provide further perspective, consider that the average increase when switching from using LDAK-Bolt-SS to LDAK-Bolt-Predict (i.e., changing from using summary statistics to individual-level data) was 2% (s.d. 2%), the average increase when switching from using directly-genotyped SNPs to imputed genotypes was 7% (s.d. 2%), the average increase when switching from using LDAK-Ridge-Predict to LDAK-Bolt-Predict (i.e., changing from a single prior distribution for effect sizes to a mixture prior) was 16% (s.d. 2%), while the average increase when switching from classical PRS to LDAK-Ridge-Predict (i.e., changing from classical PRS to the worst-performing advanced prediction tool) was 17% (s.d. 3%).

A strength of our study is that we have considered a variety of complex traits. These include continuous, binary and ordinal phenotypes, that have low, medium and high SNP heritability, and that are both closely and distantly related to diseases. Therefore, the fact that we increased prediction accuracy for almost all of the 225 phenotypes we analyzed, makes us confident that improvements will be observed for many more complex traits. Similarly, our new prediction tools have varying forms of prior distribution for SNP effect sizes. Therefore, the fact that prediction accuracy increased for all tools, indicates that if a new tool is developed with a superior prior distribution form, it is likely that this tool could also be made more accurate by improving the heritability model.

Except for ours, we are not aware of any individual-level data prediction tools that can both analyze biobank-sized datasets (say, over 50,000 samples) and allow the user to specify the heritability model. We are aware of two summary statistic prediction tools where the user can specify the heritability model, AnnoPred^32^ and LDpred-funct.^33^ AnnoPred is similar to LDAK-Bolt-SS. It assumes that SNP effect sizes have the prior distribution *p*_0_ N(0,*σ*^2^) + (1-*p*_0_) *δ*_0_, then incorporates the chosen heritability model by allowing either *σ*^2^ or *p*_0_ to vary across SNPs.^32^ Supplementary Fig. 1 shows that AnnoPred is outperformed by LDAK-Bolt-SS, regardless of whether we assume the BLD-LDAK Model (our recommended model) or the Baseline LD Model (recommended by the authors of AnnoPred). LDpred-funct is similar to LDAK-Ridge-SS. It first estimates effect sizes assuming the the prior distribution N(0,*σ*^2^), where *σ*^2^ varies across SNPs according to the chosen heritability model, then regularizes these estimates via cross-validation.^33^ Supplementary Fig. 1 shows that LDpred-funct is outperformed by LDAK-Ridge-SS, regardless of whether we assume the BLD-LDAK Model (our recommended model) or the Baseline LD Model (recommended by the authors of LDpred-funct).

When performing heritability analysis, we previously recommended choosing the heritability model with lowest AIC.^17^ We now recommend the same when constructing PRS. Based on average AIC, the BLD-LDAK, LDAK-Thin and GCTA models rank first, second and third, respectively, which matches their order when ranked based on the average accuracy of the corresponding PRS. We additionally construct PRS assuming the GCTA-LDMS-I^34^ and Baseline LD Models,^35^ those currently recommended by the authors of GCTA^8^ and LDSC,^36^ respectively. Based on average AIC, these two models rank between the LDAK-Thin and BLD-LDAK Models (Supplementary Table 2), which similarly matches their order when ranked based on the average accuracy of the corresponding PRS (Supplementary Fig. 9).

Although we observed improvement for almost all of the 225 UK Biobank phenotypes, we found that the relative advantage of our new prediction tools was largest for phenotypes with small and modest *R*^2^ (e.g., those with *R*^2^<0.1). This is relevant because, at present, most successful applications of genetic prediction models^37,38^ involve PRS with small or modest *R*^2^. For example in psychiatric research, a PRS with *R*^2^≈0.05 was used to show that impulsivity is an endophenotype for attention deficit hyperactivity disorder,^39^ a PRS with *R*^2^≈0.07 was used to show that individuals with chronic schizophrenia had higher-than-average genetic liability to schizophrenia,^40^ a PRS with *R*^2^≈0.02 was used to identify clinically-defined subtypes of autism that have significantly different genetic liabilities,^41^ a PRS with *R*^2^<0.05 was used to demonstrate that risk of developing emotional problems is moderated by an interaction between environmental sensitivity and type of parenting,^42^ and a PRS with *R*^2^≈0.01 was used to demonstrate that stressful life events and childhood trauma are risk factors for the development of major depressive disorder.^43^ Away from psychiatric research, Khera et al.^5^ demonstrated the utility of genetic risk prediction for atrial fibrillation, breast cancer, coronary artery disease, inflammatory bowel diseases and type 2 diabetes using PRS with *R*^2^ between 0.02 and 0.04.

We finish by noting that the performance of our new prediction tools will increase as more realistic heritability models are developed. To date, most of the improvement in PRS accuracy has come from increasing sample size, algorithmic innovations or developing more effective forms of prior distribution for SNP effect sizes. Our work indicates that in future, more focus should be placed on improving the heritability model.

## METHODS

We begin by explaining our new prediction tools. Note that before running each tool, it is necessary to estimate the expected heritability contributed by each SNP, given the heritability model. Our prediction tools then use these estimates to set the parameters of the effect size prior distribution for each SNP.

Suppose there are n individuals and m SNPs. Let **X** denote the matrix of genotypes (size n x m, where column **X**_*j*_ contains the genotypes for SNP *j*), and **Y** denote the vector of phenotypes (length n). For convenience, the **X**_*j*_ and **Y** are standardized, so that Mean(**X**_*j*_) = Mean(**Y**) = 0 and Var(**X**_*j*_) = Var(**Y**) = 1. We assume that the chi-squared (one degree of freedom) test statistic for SNP *j* from single-SNP analysis is *S*_*j*_ = *n r*_*j*_^2^ / (1 – *r*_*j*_^2^), where *r*_*j*_ = **X**_*j*_**Y**/*n* is the correlation between SNP *j* and the phenotype (this assumes the analysis performed linear regression, but remains a good approximation for *S*_*j*_ computed using logistic regression^44^). We consider prediction tools that use the linear model

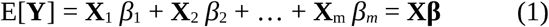

where *β*_*j*_ is the effect size for SNP *j*, and **β** = (*β*_1_, *β*_2_, …, *β*_*m*_)^T^. Because **X**_*j*_ and **Y** are standardized, the heritability contributed by SNP *j* is *h*^2^_*j*_ = *β* ^2^_*j*_.

### Heritability models

The heritability model takes the form^17^

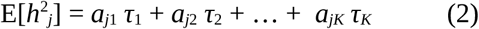

where the *a*_*jk*_ are pre-specified SNP annotations, while the parameters *τ*_*k*_ are estimated from the data.^44^ In total, we consider five heritability models (see Supplementary Tables 6 & 7 for formal definitions): the one-parameter GCTA Model assumes E[*h*^2^_*j*_] is constant;^8^ the one-parameter LDAK-Thin and 20-parameter GCTA-LDMS-I Model allow E[*h*^2^_*j*_] to vary based on MAF and local levels of linkage disequilibrium;^34,35^ the 66-parameter BLD-LDAK and 75-parameter Baseline LD Models allow E[*h*^2^_*j*_] to vary based on MAF, linkage disequilibrium and functional annotations.^17,35^ The GCTA Model is the most used model in statistical genetics.^9^ The GCTA-LDMS and Baseline LD Models are the recommended models of the authors of GCTA^8^ and LDSC,^36^ respectively. The BLD-LDAK Model is our preferred model, however, we recommend the LDAK-Thin Model for applications that demand a simple heritability model.^17^ We explain the biological intuition behind the GCTA, LDAK-Thin and BLD-LDAK Models in Supplementary Fig. 10.

For a given phenotype, we estimate the *τ*_*k*_ in Equation (2) using SumHer (an existing tool within the LDAK software), which requires summary statistics from single-SNP analysis and a reference panel.^44^ SumHer first calculates the expected value of *S*_*j*_ given the heritability model

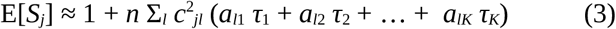

where *c*^2^_*jl*_ is the squared correlation between SNPs *j* and *l* in the reference panel, while the summation is across SNPs near SNP *j* (e.g., within 1cM). Then SumHer estimates the *τ*_*k*_ by regressing the *S*_*j*_ on the E[*S*_*j*_]. For further details see our earlier publications.^17,44^ Note that while SumHer can allow for confounding bias (by adding an extra parameter to Equation (3) designed to capture inflation of test statistics due to population structure and familial relatedness), we no longer recommend this feature, nor use it when constructing prediction models.^45^ The computational demands of SumHer depend on the complexity of the heritability model; for our analyses, it took approximately 20 minutes when assuming the GCTA or LDAK-Thin Model, and about one hour when assuming the BLD-LDAK Model (each time requiring less than 10Gb memory). As well as estimating *τ*_*k*_, SumHer also reports *e*_*j*_, the estimate of E[*h*^2^_*j*_] obtained by replacing the *τ*_*k*_ in Equation (2) with their estimated values.

### New prediction tools

Each of our new tools assumes that the error terms in Equation (1) are normally distributed, so that **Y** ~ N(**Xβ**, *σ*^2^_*e*_), where *σ*^2^_*e*_ is the residual variance. They differ in their forms of prior distributions for SNP effect sizes (Fig. 1a). big_spLinReg and LDAK-Lasso-SS use a double exponential distribution, *β*_*j*_ ~ DE(*λ* / E[*h*^2^_*j*_]^0.5^). LDAK-Ridge-Predict and LDAK-Ridge-SS use single Gaussian distributions, *β*_*j*_ ~ N(0, E[*h*^2^_*j*_]) and *β* _*j*_ ~ N(0, *v*E[*h*^2^_*j*_]), respectively. LDAK-Bolt-Predict and LDAK-Bolt-SS use a mixture of two Gaussian distributions, *β*_*j*_ ~ *p* N(0, (1-*f*_2_)/*p* E[*h*^2^_*j*_]) + (1-*p*) N(0, *f* /(1-*p*) E[*h*^2^_*j*_]). LDAK-BayesR-Predict and LDAK-BayesR-SS use a mixture of a point mass at zero and three Gaussian distributions, *β*_*j*_ ~ *π*_1_ δ_0_ + *π*_2_ N(0, *s*E[*h*^2^_*j*_]/100) + *π*_3_ N(0, *s*E[*h*^2^ _*j*_]/10) + *π*_4_ N(0, *s*E[*h*^2^ _*j*_]), where *π* _1_ + *π* _2_ + *π* _3_ + *π* _4_ = 1 and *s* = (*π* _2_ /100 + *π* _3_ /10 + *π* _4_)^−1^. The biological intuition behind the different prior distribution forms is explained in Supplementary Fig. 11. For each tool, we set E[*h*^2^_*j*_]=*e*_*j*_ (the estimate from SumHer), and *σ*^2^ _*e*_= 1-Σ*e*_*j*_. The remaining prior parameters (*λ, v, p, f*_2_, *π* _1_, *π* _2_ *π* _3_ and *π* _4_) are decided using cross-validation, as explained below.

### Model fitting using individual-level data

big_spLinReg is a function within our R package bigstatsr.^46^ The original version of the function is described in Prive et al.;^47^ the most recent version is the same, except that it allows the user to provide penalty factors that transform the prior from *β*_*j*_ ~ DE(*λ*) to *β*_*j*_ ~ DE(*λ* / E[*h*^2^_*j*_]^0.5^). In summary, big_spLinReg estimates the *β*_*j*_ using coordinate descent with warm starts.^48,49^ Given a value for *λ*, the *β*_*j*_ are updated iteratively (starting from zero) until they converge. Within each iteration, each *β*_*j*_ within the strong set (the subset of predictors determined most likely to have non-zero effects^49^) is updated once, by replacing its current value with its conditional posterior mode. *λ* starts at a value sufficiently high that *β*_*j*_ = 0 for all SNPs, then is gradually lowered to allow an increasing number of SNPs to have non-zero effects. big_spLinReg uses ten-fold cross-validation to decide when to stop reducing *λ*.

LDAK-Bolt-Predict uses the same algorithm for estimating the *β*_*j*_ and deciding values for *p* and *f*_2_ as the existing tool Bolt-LMM.^18^ In summary, LDAK-Bolt-Predict uses variational Bayes to estimate the *β*_*j*_. Given values for *p* and *f*_2_, LDAK-Bolt-Predict updates the *β*_*j*_ iteratively (starting from zero), until the approximate log likelihood converges. Within each iteration, each *β*_*j*_ is updated once, by replacing its current value with its conditional posterior mean. LDAK-Bolt-Predict considers 6 values for *p* (0.01, 0.02, 0.05, 0.1, 0.2 and 0.5) and three values for *f*_2_ (0.1, 0.3 and 0.5), resulting in 18 possible pairs for *p* and *f*_2_. First LDAK-Bolt-Predict estimates effect sizes for each of the 18 pairs using data from 90% of samples. Then it identifies which pair results in the best fitting model (based on the mean squared difference between observed and predicted phenotypes for the remaining 10% of samples). Finally, for the best-fitting pair, it re-estimates effect sizes using data from all samples. Note that whereas Bolt-LMM begins by using REML^50^ to estimate *h*^2^_SNP_, then sets E[*h*^2^_*j*_]=*h*^2^_SNP_ /*m*, LDAK-Bolt-Predict does not require this step because it instead sets E[*h*^2^_*j*_] based on estimates from SumHer (see above). Supplementary Fig. 1 shows that the results from LDAK-Bolt-Predict, when run assuming the GCTA Model, are very similar to those from Bolt-LMM.

The prior distribution used by LDAK-Ridge-Predict matches that used by LDAK-Bolt-Predict when *p* = *f*_2_ = 0.5. Therefore, LDAK-Ridge-Predict uses the same algorithm as LDAK-Bolt-Predict, except that it fixes *p* = *f*_2_ = 0.5 and it is no longer necessary to perform the cross-validation step. Supplementary Fig. 1 shows that the results from LDAK-Ridge-Predict, when run assuming the GCTA Model, are very similar to those from the existing tool BLUP^19^ (Best Linear Unbiased Prediction).

The existing tool BayesR estimates all parameters using Markov Chain Monte Carlo (MCMC).^11^ However, we do not have sufficient resources to apply BayesR to the full UK Biobank data (we estimate that this would require approximately 900Gb and weeks of CPU time). Therefore, LDAK-BayesR-Predict instead uses variational Bayes and cross-validation. The algorithm is the same as for LDAK-Bolt-Predict, except that it is now necessary to select suitable values for *π*_1_, *π*_2_, *π*_3_ and *π*_4_. In total, we consider 35 different combinations: the first is the ridge regression model (*π*_1_, *π*_2_, *π*_3_, *π*_4_) = (0, 0, 0, 1); the remaining 34 are obtained by allowing each of *π*_2_, *π*_3_ and *π*_4_ to take five values (0, 0.01, 0.05, 0.1, 0.2), with the restrictions *π*_4_ ≤ *π*_3_ ≤ *π*_2_ and *π*_2_ + *π*_3_ + *π*_4_ > 0. We investigated omitting the restriction *π*_4_ ≤ *π*_3_ ≤ *π*_2_, in which case there are 125 different triplets, however, we found that while this takes approximately four times longer to run, it did not significantly improve prediction accuracy. In Supplementary Fig. 1, we compare LDAK-BayesR-Predict to BayesR (for computational reasons, we analyze only 20,000 individuals and 99,852 SNPs); the accuracy of LDAK-BayesR-Predict is consistent with that of BayesR, yet our tool is approximately 60 times faster (takes under 20 minutes, compared to 20 hours) and requires 10 times less memory (2Gb instead of 20Gb).

The runtimes reported in the main text (approximately 50, 4, 20 and 50 hours for big_spLinReg, LDAK-Ridge-Predict, LDAK-Bolt-Predict and LDAK-BayesR-Predict, respectively) correspond to using a single CPU. However, for big_spLinReg, LDAK-Bolt-Predict and LDAK-BayesR-Predict, we also provide parallel versions. For LDAK-Bolt-Predict and LDAK-BayesR-Predict, the parallel versions utilize the fact that models corresponding to different parameter choices can be generated independently (i.e., on different CPUs). For big_spLinReg, this is not possible (because the final *β*_*j*_ for one value of *λ* are used as the starting *β*_*j*_ when *λ* is reduced), but instead, each of the ten cross-validation runs can be performed independently. Additionally, for the functions LDAK-Ridge-Predict, LDAK-Bolt-Predict and LDAK-BayesR-Predict, LDAK automatically creates a save-point every 10 iterations, so that the job can be restarted if it fails to complete within the allocated time.

### Model fitting using summary statistics

LDAK-Lasso-SS, LDAK-Ridge-SS, LDAK-Bolt-SS and LDAK-BayesR-SS are all contained within a new tool called MegaPRS. To run MegaPRS requires a reference panel and three sets of summary statistics: full summary statistics (computed using all samples), training summary statistics (computed using, say, 90% of samples) and test summary statistics (computed using the remaining samples). In some cases, you will already have (or be able to construct) training and test summary statistics. However, most likely, you will only have full summary statistics, in which case you should first generate pseudo training and test summary statistics (see below).

MegaPRS exploits that, in the absence of individual-level data, **X**_*j*_**Y** can be recovered from the results of single-SNP regression (as explained above, we assume *S*_*j*_ = *n r*_*j*_^2^ / (1 – *r*_*j*_^2^), where *n* is the sample size and *r*_*j*_ = **X**_*j*_**Y**/*n*), while **X**_*j*_**X**_*l*_ can be estimated from the reference panel (specifically, MegaPRS uses **X**_*j*_**X**_*l*_ = *n c*_*jl*_, where *c*_*jl*_ is the observed correlation between SNPs *j* and *l* in the reference panel). Note that in the equations below, **X**_*j*_, **Y** and *n* vary depending on context. When using full summary statistics, **X**_*j*_ and **Y** contain genotypes and phenotypes for all samples, and n is the total number of samples. When using training (test) summary statistics, **X**_*j*_ and **Y** contain genotypes and phenotypes for only training (test) samples, and *n* is the number of training (test) samples.

MegaPRS has three steps. In Step 1, it uses the reference panel to estimate SNP-SNP correlations. In Step 2, it constructs pairs of prediction models, first using the training summary statistics (we refer to these as the “training models”), then using the full summary statistics (the “full models”). In Step 3, it uses the test summary statistics to identify the most accurate of the training models, then reports effect sizes for the corresponding full model. For our analyses, each step took less than 30 minutes and required less than 10Gb memory.

In Step 1, MegaPRS searches the reference panel for local pairs of SNPs with significant *c*_*jl*_ (by default, we define local as within 3cM and significant as *P*<0.01 from a two-sided likelihood ratio test that *c*_*jl*_ *= 0*). MegaPRS saves the local, significant pairs in a binary file, which requires 8 bytes for each pair (one integer to save the index of the second SNP, one float to save the correlation). For the UK Biobank data, there were 260M local, significant pairs (on average, 413 per SNP), and so the corresponding binary file had size 1.9Gb.

In Step 2, MegaPRS uses the training and full summary statistics to estimates effect sizes for training and full prediction models, respectively. Pairs of training and test models correspond to different prior distribution parameters. MegaPRS constructs 11 pairs of models if running LDAK-Lasso-SS or LDAK-Ridge-SS, 132 pairs of models if running LDAK-Bolt-SS and 84 pairs of models if running LDAK-BayesR-SS (Supplementary Table 8 lists the prior parameters to which these correspond). MegaPRS estimates effect sizes iteratively using variational Bayes. As explained above (in the description of LDAK-Bolt-Predict), the variational Bayes algorithm replaces each current estimate of *β*_*j*_ with its conditional posterior mean. This is possible because for all four tools, the posterior distribution for *β*_*j*_ can be expressed in terms of **X**_*j*_**Y** and **X**_*j*_**X**_*l*_. For example, if we write the prior distribution for LDAK-Bolt-SS in the form *β*_*j*_ ~ *p* N(0, *σ*^2^_Big_) + (1-*p*) N(0, *σ*^2^_Small_), then the conditional posterior distribution of *β*_*j*_ has the form *p*’N(*μ*_Big_,*v*_Big_) + (1-*p*’)N(*μ*_Small_,*v*_Small_), where

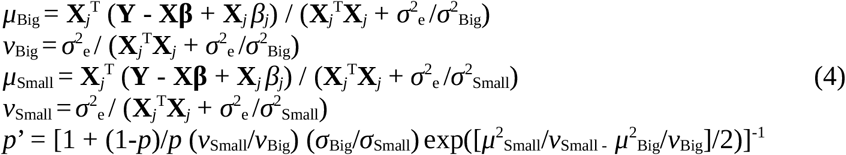

When performing variational Bayes using summary statistics, we found it was not feasible to iterate over all SNPs in the genome. This was due to differences between estimates of **X**_*j*_**X**_*l*_ from the reference panel and their true values (a consequence of the fact that individuals in the reference panel are different to those used in the original association analysis, and because we assume **X**_*j*_**X**_*l*_ = 0 for pairs of SNPs that are either distant or not significantly correlated). These differences accumulate over the genome, resulting in poor estimates of **X**_*j*_ ^T^**Xβ** = Σ_*l*_ **X**_*j*_ ^T^**X**_*l*_ *β* _*l*_, and therefore poor estimates of the conditional posterior distribution of *β*. To avoid these problems, MegaPRS uses sliding windows (see Supplementary Fig. 12 for an illustration). By default, MegaPRS iteratively estimates effect sizes for all SNPs in a 1cM window, stopping when the estimated proportion of variance explained by these SNPs converges (changes by less than 1e-5 between iterations). At this point, MegaPRS moves 1/8 cM along the genome, and repeats for the next 1cM window. Within each window, MegaPRS assumes *σ*^2^ = 1 (this approximation is reasonable because the expected heritability contributed by a single window will be close to zero). If a window fails to converge within 50 iterations, MegaPRS resets the *β*_*j*_ to their values prior to that window. We found this happened rarely. For example, our main analyses constructed 160,650 full models (across 225 phenotypes, four tools and three heritability models), and for only 990 of these (0.6%) did any of the ~12,000 regions fail to converge.

In Step 3, MegaPRS uses the test summary statistics to measure the accuracy of the training models. If **β**^’^ denotes the vector of estimated effect sizes for a model, MegaPRS calculates *R* = **β**^’T^**X**^T^**Y** /(*n* **β**’^T^**X**^T^**Xβ**’)^1/2^, an estimate of the correlation between observed and predicted phenotypes for the individuals used to compute the test summary statistics. MegaPRS then constructs the final model by extracting effect sizes from the full model corresponding to the training model with highest *R*.

In Supplementary Fig. 1, we compare LDAK-Lasso-SS with lassosum,^13^ LDAK-Ridge-SS with sBLUP,^20^ LDpred-inf^12^ and LDpred-funct,^33^ LDAK-Bolt-SS with LDpred2^51^ and AnnoPred,^32^ and LDAK-BayesR-SS with SBayesR.^14^ When we run our new tools assuming the GCTA model, they perform at least as well as the corresponding existing tools.

### Pseudo summary statistics

Given results from single-SNP analysis using *n* samples, we wish to generate two sets of results, mimicking those we would obtain from first analyzing *n*_A_ < *n* samples, then analyzing the remaining *n*_B_ = *n* - *n*_A_ samples. We can reword the task as follows. Let **γ** = (*γ*_1_, *γ*_2_, …, *γ*_*m*_)^T^ denote the vector of true SNP effect sizes from single-SNP analysis (note that *γ*_*j*_ differs from *β*_*j*_, because *β*_*j*_ reflects how much SNP *j* contributes directly to the phenotype, whereas *γ*_*j*_ reflects how much contribution the SNP tags). Given **X**^T^**Y**/*n*, the estimate of **γ** from all *n* samples, our aim is to generate **X**_A_ ^T^**Y**_A_ /*n*_A_ and **X**_B_ ^T^**Y**_B_ /*n*_B_, estimates of **γ** from *n*_A_ and *n*_B_ samples, respectively.

The method we use to generate **X**_A_ ^T^**Y**_A_ /*n*_A_ and **X**_B_ ^T^**Y** _B_/*n*_B_ is a modified version of that proposed by Zhao et al.^52^ First we sample **X** _A_ ^T^**Y** _A_ /*n*_A_ from N(**X**^T^**Y**/*n, n*_B_ /*n*_A_ **V**/*n*), where **V** is the variance of **X**^T^**Y**, then we set **X** _B_^T^**Y**_B_ /*n* _B_= (**X**^T^**Y** – **X** _A_ ^T^**Y** _A_)/*n*_B_. In their method, Zhao et al. restrict to independent SNPs, leading them to derive **V** = **I** + **X**^T^**YY**^T^**X**/*n*^2^, where **I** is an identity matrix. However, as we wish to accommodate SNPs in linkage disequilibrium, we instead use **V** = **X**^T^**X**, as proposed by Zhu and Stephens.^53^ If **X**’ denotes the (standardized) genotypes of the reference panel (size *n*’ × *m*), then an estimate of **V** is **X**’^T^**X**’*n*/*n*’, and therefore we achieve the desired sampling by setting **X**_A_ ^T^**Y**_A_ /*n* _A_ = **X**^T^**Y**/*n* + (*n*_B_ /*n*_A_)^1/2^ **X**’^T^/*n*’^1/2^ **g**, where **g** is a vector of length *n*’ with elements drawn from a standard Gaussian distribution.

As explained above, our primary use of pseudo summary statistics is to construct and test training prediction models, in order to decide parameters of the effect size prior distribution. Supplementary Fig. 13 investigates this use of pseudo summary statistics for the first 14 UK Biobank phenotypes and the eight additional diseases (those we used in Supplementary Fig. 7 and Supplementary Table 4). We see that, in general, the estimates of *R* for the training models (measured using pseudo test summary statistics) mirror the estimates of *R* for the corresponding full models (measured using the independent test data), indicating that it is valid to use pseudo summary statistics to decide prior distribution parameters. However, we note two caveats. Firstly, we observe that estimates of *R* can be unreliable when calculated using a reference panel that was also used to create the pseudo summary statistics and/or to construct the prediction models. Therefore, when running MegaPRS using pseudo partial summary statistics, we ensure that the reference panel used in Step 3 is distinct to the reference panel used in Steps 1 & 2. Secondly, we found that estimates of *R* can be unreliable when there are strong effect loci within regions of long-range linkage disequilibrium (e.g, this was an issue for rheumatoid arthritis, where a single SNP within the major histocompatibility complex explains 2% of phenotypic variation). Therefore, when estimating *R* using pseudo summary statistics, we recommend excluding regions of long-range linkage disequilibrium (a list of these are provided at www.ldak.org/high-ld-regions).

### Data

When using our individual-level tools, we constructed prediction models for 14 phenotypes from UK Biobank,^21,22^ for which we have access to phenotype and genotype data via Application 21432. These phenotypes are: body mass index (data field 21001), forced vital capacity (3062), height (50), impedance (23106), neuroticism score (20127), pulse rate (102), reaction time (20023), systolic blood pressure (4080), college education (6138), ever smoked (20160), hypertension (20002), snorer (1210), difficulty falling asleep (1200) and preference for evenings (1180). Starting with all 487k UK Biobank individuals, we first filtered based on ancestry (we only kept individuals who were both recorded and inferred through principal component analysis to be white British),^17^ then filtered so that no pair remained with allelic correlation >0.0325 (that expected for second cousins). Depending on phenotype, there were between 220,399 and 253,314 individuals (in total, 392,214 unique). From these, we picked 200,000 and 20,000 individuals to use for training and testing prediction models, respectively. For all analyses, we used adjusted phenotypes, obtained by regressing the original phenotypic values on 13 covariates (across all 220,000 training and test individuals). These covariates are age (data field 21022), sex (31), Townsend Deprivation Index (189) and ten principal components (five from the UK Biobank data, five derived from the 1000 Genomes Project^54^ data). Supplementary Table 1 reports the estimated proportion of phenotypic variation explained by cryptic relatedness (population structure and familial relatedness); across the 14 phenotypes, it is at most 0.001, and never significant (all P>0.7 from a one-sided likelihood ratio test).

The UK Biobank provides imputed genotype data, but in general we restricted to the 628,694 autosomal SNPs with information score >0.9, MAF >0.01 and present on the UK Biobank Axiom Array (the exception is for Supplementary Fig. 6, where we did not require that SNPs were present on the Axiom Array). We converted dosages to genotypes using a hard-call-threshold of 0.1 (i.e., dosages were rounded to the nearest integer, unless they were between 0.1 and 0.9 or between 1.1 and 1.9, in which case the corresponding genotype was considered missing). After this conversion, on average, 0.1% of genotypes were missing. Note that big_spLinReg does not allow missing values, so when using this tool, we used a hard-call-threshold of 0.5.

When using our summary statistic tools, we constructed prediction models for 225 phenotypes from UK Biobank (Supplementary Data 1), using the August 2018 results from the Neale Lab. In total, the Neale Lab analyzed 4,203 UK Biobank phenotypes, using up to 361,194 British individuals. We downloaded results for the 283 phenotypes that were computed using both sexes and had estimated SNP heritability >0.05 (using details provided in the file ukb31063_h2_topline.02Oct2019.tsv.gz). For each phenotype, we begun by generating pseudo training and test summary statistics corresponding to 90% and 10% of samples, respectively. We subsequently used the pseudo training summary statistics to construct prediction models, and the pseudo test summary statistics to measure their accuracy. Supplementary Fig. 2 confirms that it is possible to both construct and (fairly) measure the accuracy of different prediction models using a single set of summary statistics. Specifically, it shows that for the first 14 phenotypes, estimates of *R*^2^ are similar whether we use data from our own UK Biobank application (for which we have independent training and test data) or use summary statistics from the Neale Lab. Although we downloaded results for 283 phenotypes, in the main text, we restrict to the 225 phenotypes for which it was possible to generate a PRS with *R*^2^>0.01 (using any tool and any heritability model). We made this choice because it is difficult to reliably compare the performance of tools using PRS with very low, and often non-significant, *R*^2^. However, we note that for the 58 phenotypes we rejected, it remained that improving the heritability model increased *R*^2^ on average 93% of time, and that the average improvement in *R*^2^ was 19% (s.d. 3%) when we switched from the best existing tool to LDAK-BayesR-SS assuming the BLD-LDAK Model.

When we required a reference panel, we used genotypes of 20,000 individuals from the UK Biobank (when requiring multiple reference panels, we ensured 20,000 different individuals were used for each). Note that when using individual-level data prediction tools, for which we use a reference panel to estimate E[*h*^2^_*j*_] given the heritability model, we always picked the 20,000 individuals from the 200,000 training samples, to ensure that there was no overlap between the data used to train and test prediction models.

For the analysis in Supplementary Fig. 7 and Supplementary Table 4, we used results from published studies to construct PRS for eight diseases: asthma,^29^ atrial fibrillation,^28^ breast cancer,^31^ inflammatory bowel disease,^26^ prostate cancer,^30^ rheumatoid arthritis,^24^ schizophrenia^25^ and type 2 diabetes.^27^ We chose these diseases as they were the ones for which we could find both cases in the UK Biobank and publicly-available summary statistics from a genome-wide association study that did not use UK Biobank data. We excluded SNPs with ambiguous alleles (A&T or C&G) or that were not present in our UK Biobank dataset, after which on average 470,000 SNPs remained (range 191,000 to 559,000).

For the analysis in Supplementary Fig. 8 and Supplementary Table 5, we measured how well prediction models for the first 14 UK Biobank phenotypes (constructed using data from white British individuals) performed for individuals of non-European ancestry. For this, we used principal component analysis to identify 7,057, 2,717 and 1,331 individuals from the UK Biobank whose ancestries were consistent with individuals reported to be South Asian (Indian or Pakistani), African and East Asian (Chinese), respectively.

### Sensitivity of MegaPRS to setting choices

In Supplementary Fig. 14, we test the impact on prediction accuracy of changing the definitions of local and significant when calculating SNP-SNP correlations, the window settings and convergence threshold used when estimating effect sizes, and the choice of reference panel. In general, the impact on accuracy is small. It is largest when we replace the UK Biobank reference panel (20,000 individuals) with genotypes of 489 European individuals from the 1000 Genome Project.^54^ In this case, average *R*^2^ reduces by approximately 3% (about two thirds of this is due to reducing the number of individuals, and one third due to replacing UK Biobank genotypes with 1000 Genome Project genotypes).

### Other prediction tools

Supplementary Fig. 1 compares the performance of our new tools with nine existing tools, using the first 14 UK Biobank phenotypes. Note that when using summary statistic tools, we use our own UK Biobank data, rather than summary statistics from the Neale Lab (so there was no need to use pseudo summary statistics). When running BLUP,^19^ Bolt-LMM,^18^ BayesR,^11^ lassosum,^13^ sBLUP,^20^ LDpred-funct,^33^ LDpred2,^51^ AnnoPred^32^ and SBayesR,^14^ we used the default settings of each software (see Supplementary Note 1 for scripts). When it was necessary to select prior parameters (this was the case for lassosum, LDpred2 and AnnoPred, as well as for our four summary statistic tools), we used cross-validation. Similar to above, we constructed models corresponding to different parameter choices using 90% of the training samples (180,000 individuals), then tested these using the remaining 10% of the training samples (20,000 individuals). Having identified the best-performing parameters, we then used these to make the final model (using all training samples). Note that for sBLUP, we found that average *R*^2^ improved if we repeated the analyses excluding regions of long-range linkage disequilibrium,^14,33^ while for AnnoPred, we found it was necessary to exclude SNPs from the major histocompatibility complex (otherwise, the software would fail to complete).

When constructing Classical PRS, we used estimates of *β*_*j*_ from single-SNP analysis. We considered six p-value thresholds (*P*≤5e-8, *P*≤.0001, *P*≤0.001, *P*≤0.01, *P*≤0.1, all SNPs) and four clumping thresholds (*c*^2^ _*jk*_≤0.2, *c*^2^_*jk*_ ≤0.5, *c*^2^_*jk*_ ≤0.8, and no clumping). We decided the most-suitable pair of thresholds via cross-validation. Similar to above, we first constructed 18 models using 90% of training samples, then tested these models using the remaining 10% of training samples. We then used the best-performing p-value and clumping thresholds to construct the final model (using summary statistics from all training samples).

## DATA AVAILABILITY

Individual-level UK Biobank data can be applied for from www.ukbiobank.ac.uk. Neale Lab summary statistics can be downloaded from www.nealelab.is/uk-biobank. Summary statistics for the eight additional diseases can be downloaded from the websites of the corresponding studies: asthma (www.ebi.ac.uk/gwas/studies/GCST006862), atrial fibrillation (www.ebi.ac.uk/gwas/studies/GCST004296), breast cancer (bcac.ccge.medschl.cam.ac.uk/bcacdata/oncoarray), inflammatory bowel disease (www.ibdgenetics.org/downloads.html), prostate cancer (http://practical.icr.ac.uk/blog/?page_id=8164), rheumatoid arthritis (plaza.umin.ac.jp/~yokada/datasource/software.htm), schizophrenia (www.med.unc.edu/pgc/download-results/) and type 2 diabetes (diagram-consortium.org/downloads.html).

## CODE AVAILABILITY

We provide step-by-step scripts for constructing prediction models in Supplementary Note 1. Our eight new prediction tools are provided within our software packages LDAK (available from www.ldak.org) and bigstatsr (privefl.github.io/bigstatsr). When comparing against existing prediction tools, we additionally used the software packages Bolt-LMM (data.broadinstitute.org/alkesgroup/BOLT-LMM), gctb (cnsgenomics.com/software/gctb), lassosum (github.com/tshmak/lassosum), bigsnpr (privefl.github.io/bigsnpr), LDpred-funct, (github.com/carlaml/Ldpred-funct) and AnnoPred, (github.com/yiminghu/AnnoPred).

## ACKNOWLEDGMENTS

The authors thank Dr Veera Rajagopal for testing the LDAK software. F.P. and B.J.V. are supported by the Danish National Research Foundation (Niels Bohr Professorship to Prof. John McGrath) and the Lundbeck Foundation Initiative for Integrative Psychiatric Research, iPSYCH (R102-A9118, R155-2014-1724 and R248-2017-2003). B.J.V. is also supported by a Lundbeck Foundation Fellowship (R335-2019-2339). D.S. is supported by the European Union’s Horizon 2020 Research and Innovation Programme under the Marie Skłodowska-Curie grant agreement no. 754513, by Aarhus University Research Foundation (AUFF), by the Independent Research Fund Denmark under Project no. 7025-00094B, and by a Lundbeck Foundation Experiment Grant.

## AUTHOR CONTRIBUTIONS

D.S., Q.Z. and F.P. performed the analyses, D.S, F.P. and B.V. wrote the paper.

