## Supplementary Information for "Improved genetic prediction of complex traits from individual-level data or summary statistics"

**Supplementary Note 1: Scripts for repeating our analyses of UK Biobank phenotypes**

**Supplementary Note 2: Algorithmic improvements.**

**Supplementary Figure 1: Comparing our new tools with existing tools.**

**Supplementary Figure 2: Impact of changing the heritability model when using summary statistics.**

**Supplementary Figure 3: Impact of improving the heritability model for simulated data.**

**Supplementary Figure 4: Impact of increasing the training sample size.**

**Supplementary Figure 5: Alternative measures of prediction accuracy.**

**Supplementary Figure 6: Including imputed SNP genotypes.**

**Supplementary Figure 7: Eight additional diseases.**

**Supplementary Figure 8: Cross-ancestry prediction.**

**Supplementary Figure 9: Alternative heritability models.**

**Supplementary Figure 10: Comparison of heritability models.**

**Supplementary Figure 11: Comparison of prior distribution forms.**

**Supplementary Figure 12: A sliding window approach for estimating effect sizes.**

**Supplementary Figure 13: Validity of pseudo summary statistics.**

**Supplementary Figure 14: Sensitivity of MegaPRS to setting choices.**

**Supplementary Table 1: Heritabilities and best prediction models for the first 14 UK Biobank phenotypes.**

**Supplementary Table 2: Fit of heritability models for the first 14 UK Biobank phenotypes.**

**Supplementary Table 3: Accuracy of prediction models constructed from individual-level data.**

**Supplementary Table 4: Eight additional diseases.**

**Supplementary Table 5: Cross-ancestry prediction.**

**Supplementary Table 6: Definitions of heritability models.**

**Supplementary Table 7: Baseline LD Model SNP annotations.**

**Supplementary Table 8: Prior distribution parameter choices.**

**Supplementary Note 1: Scripts for repeating our analyses of UK Biobank phenotypes.** Here we provide the scripts we used to analyze the UK Biobank phenotypes using our new prediction tools. In general, the scripts use LDAK version 5.1 (www.ldak.org); the exception is when using big_spLineReg, for which the scripts also use the R package `bigstatsr` (rdrr.io/github/privefl/bigstatsr). At the end, we summarize the scripts we used to compare our new tools to existing tools.

Note that we provide these scripts mainly for completeness. If you wish to learn how to use our new tools LDAK-Ridge-Predict, LDAK-Bolt-Predict, LDAK-BayesR-Predict, LDAK-Lasso-SS, LDAK-Ridge-SS, LDAK-Bolt-SS and LDAK-BayesR-SS, you should visit the LDAK website, which provides instructions and test datasets, as well as tutorials on quality control, advice on choosing the heritability model and details of existing functions within LDAK. If you wish to use big_spLinReg, you should additionally visit the bigstatsr website. As a reminder, we recommend using LDAK-Bolt-Predict if using individual-level data, and LDAK-BayesR-SS if using summary statistics.

For the scripts below, we assume that the LDAK executable is called `ldak.out` (in practice, it will have a name like `ldak5.1.linux`, `ldak5.1.linux.fast` or `ldak5.1.mac`, depending on which version you download). We assume that the (main) reference panel is stored in PLINK format in the files `ref.bed`, `ref.bim` & `ref.fam`, and that summary statistics from single-SNP regression are stored in the file `linear.trait.summaries` (the required format of this file is described at www.ldak.org/summary-statistics). If you are performing the single-SNP analysis yourself, you can use LDAK with the command `-linear` (see www.ldak.com/single-predictor-analysis). To construct PRS assuming the BLD-LDAK Model (our recommended heritability model when analyzing human data), you must download the SNP annotation files `bld1`, `bld2`, ..., `bld64` from www.ldak.org/bldldak. To ensure consistency, we advise that all SNP names are in the format Chr:BP, where Chr and BP denote the chromosome and basepair of the SNP, respectively (in particular, this is the format of the BLD-LDAK Model SNP annotations).

When constructing PRS from individual-level data, we assume that the genotype data for training samples are stored in Binary PLINK format with the prefix `data`, and that the corresponding phenotypes are stored in the file `trait.pheno` (three columns, ID1, ID2, Phenotype). Note that `bigstatsr` does not allow missing values in the training genotype data (missing values are allowed when using LDAK). When testing PRS, we assume that the genotype data for test samples are stored in Binary PLINK format with the prefix `data.test`, and that the corresponding phenotypes are stored in the file `trait.test.pheno`.

When constructing PRS from summary statistics and using pseudo summary statistics, we use two extra reference panels. When using summary statistics from the Neale Lab, we generated pseudo summary statistics twice, and therefore used four extra reference panels. In the scripts below, we assume the extra reference panels are stored in PLINK format with the prefixes `ref2`, `ref3`, `ref4` and `ref5`. Additionally, it is necessary to identify SNPs within regions of high linkage disequilibrium, for which we downloaded the file `highld.txt` from www.ldak.org/high-ld-regions.

Step 1 explains how to estimate the heritability contributed by each predictor, given the heritability model. Steps 2 & 3 explain how to create PRS using our individual-level data prediction tools. Step 4 explains how to create PRS using our individual-level data prediction tools. Step 5 explains how to test PRS using an independent test dataset. Step 6 explains how, when using summary statistics from the Neale Lab (www.nealelab.is/uk-biobank), we were able to both construct and test PRS using a single set of summary statistics. Step 7 summarizes the scripts we used for constructing PRS using existing tools. Step 8 explains how we generated the simulated phenotypes used in Supplementary Figure 3.

When running LDAK, always watch the screen output, which will suggest options and explain how to fix any errors that occur.

```
#Step 1 - How to use SumHer to estimate total and per-SNP heritabilities given the heritability model
#Here we consider three heritability models, the GCTA, LDAK-Thin and BLD-LDAK Models
#We use a reference panel (ref.bed, ref.bim and ref.fam) and summary statistics (linear.trait.summaries)
#When assuming the BLD-LDAK Model, we also use the annotations bld1, bld2, ..., bld64

#GCTA Model
#All SNPs are given equal weight (effected by using --ignore-weights YES and --power -1)
ldak.out --calc-tagging gcta --bfile ref --ignore-weights YES --power -1 --window-cm 1 --save-matrix YES
ldak.out --sum-hers trait.gcta --tagfile gcta.tagging --summary linear.trait.summaries --matrix gcta.matrix

#LDAK-Thin Model
#SNPs are thinned for duplicates, then weighted based only on MAF (--ignore-weights YES and --power -0.25)
ldak.out --thin thin --bfile ref --window-prune .98 --window-kb 100
ldak.out --calc-tagging thin --bfile ref --extract thin.in --ignore-weights YES --power -.25 --window-cm 1
    --save-matrix YES
ldak.out --sum-hers trait.thin --tagfile thin.tagging --summary linear.trait.summaries --matrix thin.matrix

#BLD-LDAK Model
#Calculate the LDAK weightings, then add these to the 64 SNP annotations from www.ldak.org/bldldak
ldak.out --cut-weights sections --bfile ref
ldak.out --calc-weights-all sections --bfile ref
cp sections/weights.shorts bld65
ldak.out --calc-tagging bldldak --bfile ref --ignore-weights YES --power -.25 --annotation-number 65
    --annotation-prefix bld --window-cm 1 --save-matrix YES
ldak.out --sum-hers trait.bldldak --tagfile bldldak.tagging --summary linear.trait.summaries --matrix bldldak.matrix

#Notes
#The final lines of the ".hers" files provide the estimates of total SNP heritability
#The ".ind.hers.positive" files contain the estimates of per-SNP heritabilities

#################

#Step 2 - How to construct a prediction model using big_spLinReg
#Here we assume the BLD-LDAK Model (so use the file trait.bldldak.ind.hers.positive from Step 1)
#We use genotype data (data.bed, data.bim and data.fam) and phenotypes (trait.pheno)
#These scripts should be run from R; they assume you have first installed the R packages bigstatsr and bigsnpr

#Load the required package, then resave the genotypes as data.rds and data.bk
library("bigsnpr")
snp_readBed("data.bed")

#Load the data and phenotypes
obj.bigSNP=snp_attach("data.rds")
X=obj.bigSNP$genotypes
fam=obj.bigSNP$fam
map=obj.bigSNP$map
Y=read.table("trait.pheno")

#Match up individuals between the genotype and phenotype data
use=intersect(paste(fam[,1],"___",fam[,2],sep=""),paste(Y[,1],"___",Y[,2],sep=""))
X.index=match(use,paste(fam[,1],"___",fam[,2],sep=""))
Y.index=match(use,paste(Y[,1],"___",Y[,2],sep=""))

#Get the penalty factors
indhers=read.table("trait.bldldak.ind.hers.positive")
scales=rep(0,nrow(map))
scales[match(indhers[,1],map[,2])]=indhers[,2]
```

```
pred.index=which(scales>0)

#Fit the model
fit=big_spLinReg(X,Y[Y.index,3],X.index,pred.index,dfmax=Inf,pf.X=1/scales[pred.index]^.5)
print(summary(fit)$message)

#Save the effects in the format required by LDAK
effects=cbind(map[,c(2,5,6)],NA,0)
effects[pred.index,5]=summary(fit, best.only=TRUE)$beta[[1]]
colnames(effects)=c("Predictor","A1","A2","Centre","Effect")
write.table(effects[which(effects[,5]!=0),],"lasso.bldldak.effects",row=F,quote=F)

#Note
#The function big_spLinReg can be run across multiple CPUs by using the option "ncores"

#################

#Step 3 - How to construct a prediction model using LDAK-Ridge-Predict, LDAK-Bolt-Predict or LDAK-BayesR-Predict
#Here we assume the BLD-LDAK Model (so use the file trait.bldldak.ind.hers.positive from Step 1)
#We use genotype data (data.bed, data.bim and data.fam) and phenotypes (trait.pheno)

#LDAK-Ridge-Predict
ldak.out --ridge blup.trait.bldldak --bfile data --ind-hers trait.bldldak.ind.hers.positive --pheno trait.pheno

#LDAK-Bolt-Predict
ldak.out --bolt bolt.trait.bldldak --bfile data --ind-hers trait.bldldak.ind.hers.positive --pheno trait.pheno
    --cv-proportion .1

#LDAK-BayesR-Predict
ldak.out --bayesr bayesr.trait.bldldak --bfile data --ind-hers trait.bldldak.ind.hers.positive --pheno trait.pheno
    --cv-proportion .1

#Notes
#The effect sizes will be saved in the ".effects" files
#If jobs die before completion, you can resume them by rerunning adding "--restart YES"
#To run across multiple CPUs (Linux only), use the "fast" version of LDAK adding, say, "--max-threads 4"

#################

#Step 4 - How to construct a prediction model using LDAK-Lasso-SS, LDAK-Ridge-SS, LDAK-Bolt-SS or LDAK-BayesR-SS
#Here we assume the BLD-LDAK Model (so use the file trait.bldldak.ind.hers.positive from Step 1)
#We use three reference panel (prefixes ref, ref2 & ref3) and summary statistics (linear.trait.summaries)
#We also use the file highld.txt, that specifies the long-range linkage disequilibrium regions in the human genome

#A - generate pseudo partial summary statistics (not required if actual partial summary statistics are available)
ldak.out --pseudo-summaries pseudo.trait --bfile ref2 --summary linear.trait.summaries --training-proportion .9

#B - calculate SNP-SNP correlations
ldak.out --calc-cors cors --bfile ref --window-cm 3

#C - estimate effect sizes for training and full models

#If using LDAK-Lasso-SS:
ldak.out --mega-prs mega.trait.bldldak --bfile ref --cors cors --ind-hers trait.bldldak.ind.hers.positive --summary
    linear.trait.summaries --summary2 pseudo.train.summaries --window-cm 1 --model lasso

#If using LDAK-Ridge-SS:
ldak.out --mega-prs mega.trait.bldldak --bfile ref --cors cors --ind-hers trait.bldldak.ind.hers.positive --summary
    linear.trait.summaries --summary2 pseudo.train.summaries --window-cm 1 --model ridge
```

```
#If using LDAK-Bolt-SS:
ldak.out --mega-prs mega.trait.bldldak --bfile ref --cors cors --ind-hers trait.bldldak.ind.hers.positive --summary
    linear.trait.summaries --summary2 pseudo.train.summaries --window-cm 1 --model bolt

#If using LDAK-BayesR-SS:
ldak.out --mega-prs mega.trait.bldldak --bfile ref --cors cors --ind-hers trait.bldldak.ind.hers.positive --summary
    linear.trait.summaries --summary2 pseudo.train.summaries --window-cm 1 --model bayesr

#Note that when comparing with lassosum, we used LDAK-Lasso-Sparse-SS:
ldak.out --mega-prs mega.trait.bldldak --bfile ref --cors cors --ind-hers trait.bldldak.ind.hers.positive --summary
    linear.trait.summaries --summary2 pseudo.train.summaries --window-cm 1 --model lasso-sparse

#While when comparing with LDpred, we used LDAK-Bolt-Sparse-SS:
ldak.out --mega-prs mega.trait.bldldak --bfile ref --cors cors --ind-hers trait.bldldak.ind.hers.positive --summary
    linear.trait.summaries --summary2 pseudo.train.summaries --window-cm 1 --model bolt --LDpred YES

#D - identify SNPs within long-range linkage disequilibrium regions
ldak.out --cut-genes highld --bfile ref --genefile highld.txt

#E - measure the accuracy of the training models (excluding high-LD SNPs), and construct the final model
ldak.out --calc-scores mega.trait.bldldak --bfile ref3 --scorefile mega.trait.bldldak.effects.train --power 0
    --summary pseudo.test.summaries --final-effects mega.trait.bldldak.effects.final --exclude
    highld/genes.predictors.used

#Notes
#The effect sizes of the final model will be saved in mega.trait.bldldak.effects.best

#################

#Step 5 - How to create PRS (using the final prediction model) and test them using independent test data
#Here we use the model saved in bolt.trait.bldldak.effects
#We use genotype data (data.test.bed, data.test.bim and data.test.fam) and phenotypes (trait.test.pheno)

ldak.out --calc-scores bolt.trait.bldldak --bfile data.test --scorefile bolt.trait.bldldak.effects --power 0
    --pheno trait.test.pheno
ldak.out --jackknife bolt.trait.bldldak --profile bolt.trait.bldldak.profile --num-blocks 200

#Notes
#The polygenic risk scores for the test individuals will be saved in bolt.trait.bldldak.profile
#The accuracy of the model will be saved in bolt.trait.bldldak.jack; this file reports the correlation between
    observed and predicted phenotypes, mean squared error and mean absolute error
#To also compute AUC (for binary traits), add the option "--AUC YES" when jackknifing

#################

#Step 6 - How we constructed and tested PRS using summary statistics from the Neale Lab
#Suppose the summary statistics are saved in neale.summaries
#In total, we used five independent reference panels (prefixes ref, ref2, ref3, ref4 & ref5)
#We also used the file highld.txt and the SNP annotations bld1, bld2, ..., bld64

#A - generate pseudo training and test summary statistics (corresponding to 90% and 10% of samples), then rename
ldak.out --pseudo-summaries neale --bfile ref4 --summary neale.summaries --training-proportion .9 --allow-ambiguous
    YES
mv neale.train.summaries neale.90.summaries
mv neale.test.summaries neale.10.summaries

#We will use neale.90.summaries to construct the PRS, then test these using neale.10.summaries
```

```
#B - identify SNPs within long-range linkage disequilibrium regions
ldak.out --cut-genes highld --bfile ref --genefile highld.txt

#C - estimate per-SNP heritabilities assuming the BLD-LDAK Model (see Step 1 for other heritability models)
ldak.out --cut-weights sections --bfile ref
ldak.out --calc-weights-all sections --bfile ref
cp sections/weights.shorts bld65
ldak.out --calc-tagging bldldak --bfile ref --ignore-weights YES --power -.25 --annotation-number 65
    --annotation-prefix bld --window-cm 1 --save-matrix YES
ldak.out --sum-hers neale.bldldak --tagfile bldldak.tagging --summary neale.90.summaries --matrix bldldak.matrix

#D - construct the PRS using LDAK-BayesR-SS (see Step 4 for other tools)
ldak.out --pseudo-summaries neale.90 --bfile ref2 --summary neale.90.summaries --training-proportion .9
    --allow-ambiguous YES
ldak.out --calc-cors cors --bfile ref --window-cm 3
ldak.out --mega-prs neale.bldldak --bfile ref --cors cors --ind-hers neale.bldldak.ind.hers.positive --summary
    neale.90.summaries --summary2 neale.90.train.summaries --window-cm 1 --model bayesr --allow-ambiguous YES
ldak.out --calc-scores neale.bldldak --bfile ref3 --scorefile neale.bldldak.effects.train --power 0 --summary
    neale.90.test.summaries --final-effects neale.bldldak.effects.final --allow-ambiguous YES --exclude
    highld/genes.predictors.used

#The final prediction model is saved in neale.bldldak.effects.best

#E - measure the accuracy of the final prediction model
ldak.out --calc-scores neale.bldldak --bfile ref5 --scorefile neale.bldldak.effects.best --power 0 --summary
    neale.10.summaries --allow-ambiguous YES --exclude highld/genes.predictors.used

#Notes
#The estimated accuracy of the final PRS (measured by correlation, R) is saved in neale.bldldak.cors; as we do not
    have individual-level test data, we can not jackknife to obtain precision, nor compute measures such as AUC
#We added "--allow-ambiguous YES" to many commands because we are sure that the orientations of SNPs in the summary
    statistics match those in the reference panel (this is because the summary statistics were created from UK
    Biobank data, which we use as our reference panel)

#################

#Step 7 - Key commands from our analyses using existing software

#BLUP (here we use the GCTA Model, but we also used the LDAK-Thin Model)
#For computational reasons, we used only 50,000 individuals

ldak.out --calc-kins-direct blup.gcta --bfile data --ignore-weights --power -1
ldak.out --decompose blup.gcta --grm blup.gcta
ldak.out --reml blup.trait.gcta --grm blup.gcta --eigen blup.gcta --pheno trait.pheno
ldak.out --calc-blup blup.trait.gcta --bfile data --grm blup.gcta --remlfile blup.trait.gcta.reml

###

#Original Bolt-LMM

#The phenotype file must be in Bolt-LMM format (e.g., have column headings ID1, ID2 and Phen)
BOLT-LMM_v2.3.4/bolt --bfile data --phenoFile trait.bolt.pheno --phenoCol Phen --lmm --maxMissingPerSnp 0.5
    --maxMissingPerIndiv 0.5 --predBetasFile bolt.trait --statsFile output --LDscoresUseChip

#To force the ridge regression model, we repeated adding --pEst .5 --varFrac2Est .5

###

#BayesR
```

```
#For computational reasons, we used only 20,000 individuals and Chromosomes 1 & 2
gctb_2.0_Linux/gctb --bayes R --bfile data --pheno trait.pheno --out bayesr.trait

###

#lassosum (run from within R; requires the package lassosum)
#We copied the scripts provided at https://github.com/tshmak/lassosum

#The main two commands were (which we run for each chromosome in turn)
cor <- p2cor(p = ss$P_val, n = XXX, sign=ss$beta)
out <- lassosum.pipeline(cor=cor, chr=ss$Chr, pos=ss$Position, A1=ss$A1, A2=ss$A2, ref.bfile=reffile,
    test.bfile=reffile, LDblocks = LDblocks, exclude.ambiguous = FALSE, trace=1, destandardize = T)
#We replaced XXX with 180,000 when creating the partial models and 200,000 when creating the full models

#To save the results in the format required by LDAK
betas=matrix(unlist(out$beta),nrow(ss),byrow=F)
final=cbind(ss[,c(1,4,5)],"NA",betas)
colnames(final)=c("Predictor","A1","A2","Centre",paste("Effect",1:80,sep=""))
write.table(final,"lassosum.trait.scores",row=F,quote=F)

#We merged effect sizes across chromosomes PRIOR to deciding the best full model by cross-validation

###

#sBLUP
#The summary statistics must be in ma format (see https://cnsgenomics.com/software/gcta/#COJO for details)

gcta_1.93.2beta/gcta64 --bfile ref --cojo-file linear.trait.ma --cojo-sblup XXX --cojo-wind 1000 --out sblup.trait
#We replaced XXX with m(1-her-1), where m is number of SNPs, her is the SumHer estimate of SNP heritability

#We then repeated the analysis, excluding SNPs within the regions detailed at www.ldak.org/high-ld-regions

###

#LDpred-inf and LDpred-funct (here we use GCTA Model, but we also used LDAK-Thin, BLD-LDAK and Baseline LD Models)
#We copied the scripts provided at https://github.com/carlaml/LDpred-funct

#The main command was
miniconda2/bin/python2 LDpred-funct/ldpredfunct.py --gf=ref.[1:22] --FUNCT_FILE=trait.gcta.ind.hers.positive
    --ssf=linear.trait.sum --pf=trait.ldpred.pheno --N=200000 --H2=XXX --coord=output
    --posterior_means=ldpredinf.trait.gcta --out=ldpredfunct.trait.gcta --maf="0.00001" --skip_ambiguous
#We replaced XXX with the SumHer estimate of SNP heritability

###

#LDpred2
#We copied the scripts provided at https://privefl.github.io/bigsnpr/articles/LDpred2.html

#The main commands were (which we run for each chromosome in turn)
h2_seq <- round(h2_est * c(0.7, 1, 1.4), 4)
p_seq <- signif(seq_log(1e-4, 1, length.out = 17), 2)
params <- expand.grid(p = p_seq, h2 = h2_seq, sparse = c(FALSE, TRUE))
corr0 <- snp_cor(G, ind.col = ind.chr2, infos.pos = POS2[ind.chr2], size = 3/1000)
corr <- bigsparser::as_SFBM(as(corr0, "dgCMatrix"))
beta_grid <- snp_ldpred2_grid(corr, df_beta, params, ncores = NCORES)
pred_grid <- big_prodMat(G, beta_grid, ind.col = ind.chr2)

#We merged effect sizes across chromosomes PRIOR to deciding the best full model by cross-validation
```

```
###

#Annopred (here we use the GCTA Model, but we also used the LDAK-Thin, BLD-LDAK and Baseline LD Models)
#We copied the scripts provided at https://github.com/yiminghu/AnnoPred
#Except we found it was necessary to exclude the MHC region (otherwise AnnoPred would fail to complete)

#The main command was
miniconda2/bin/python2 AnnoPred/AnnoPred.py --ref_gt=ref --val_gt=ref --user_h2=trait.gcta.ind.hers.positive
    --sumstats=linear.trait.annopred --N_sample=XXX --coord_out=apred/output --out=annopred.trait.gcta
    --annotation_flag=tier3 --P=YYY --local_ld_prefix=output2 --temp_dir=output3
#We replaced XXX with 180,000 when creating the partial models and 200,000 when creating the full models
#As recommended, we considered 11 values for YYY (1,0.3,0.1,0.03,0.01,0.003,0.001,0.0003,0.0001,3e-05,1e-05)

###

#SBayesR
#The summary statistics must be in ma format (see https://cnsgenomics.com/software/gcta/#COJO for details)

#The reference panel must be divided by chromosome (suppose genotypes for Chromosome j have prefix ref$j)
for j in {1..22}; do
gctb_2.0_Linux/gctb --bfile ref$j --make-full-ldm --out ref$j
gctb_2.0_Linux/gctb --sbayes R --ldm ref$j --pi 0.95,0.02,0.02,0.01 --gamma 0.0,0.01,0.1,1 --gwas-summary
    linear.trait.ma --chain-length 10000 --burn-in 2000 --out-freq 10 --out sbayesr.trait$j
done

#This produces one set of effect sizes per chromosome, which we then merged to obtain the final model

#################

#Step 8 - Generating simulated phenotypes

for h in {.1,.2,.3,.4,.5}; do
for N in {10000,50000}; do
ldak.out --make-phenos phen.$h.$N --weights trait.bldldak.ind.hers.positive --num-causals $N --num-phenos 20 --her
    $h --power -1 --bfile data
done
done

#In total, this creates 200 simulated phenotypes, each with heritability 0.1, 0.2, 0.3, 0.4 or 0.5, and either
    10000 or 50000 causal SNPs (whose effects sizes are consistent with the BLD-LDAK Model).
```

**Supplementary Note 2: Algorithmic improvements.** In Extended Data Fig. 1, we compared our new prediction tools with existing tools, using the first 14 UK Biobank phenotypes. In theory, results from our new tools, when run assuming the GCTA Model, should match those from the corresponding existing tools (which automatically assume the GCTA Model). However, in many cases, our new tools performed substantially better. Here we explain the differences. Note that to understand this section, it will help if you have already read the technical details of our new prediction tools in Methods.

**Individual-level data tools.** We found that the accuracies of PRS from LDAK-Bolt-Predict were almost identical to those from the existing tool Bolt-LMM.[1] This reflects that our tool uses the same algorithms to estimate effect sizes and performs cross-validation as Bolt-LMM. The main difference between the two tools is that whereas Bolt-LMM uses REML[2] to estimate $h^2_{\text{SNP}}$, then sets $\mathbb{E}[h^2_j] = h^2_{\text{SNP}}/m$, LDAK-Bolt-Predict instead uses the estimates from SumHer.[3] Further, having selected values for $p$ and $f_2$ via cross-validation, Bolt-LMM estimates effect sizes for the final model from scratch (i.e., each $\beta_j$ starts at zero), whereas LDAK-Bolt-Predict updates estimates (i.e., the $\beta_j$ start at their values estimated from 90% of training samples). However, our comparison of the two tools indicate that these two differences have limited impact.

We found that the accuracies of PRS from LDAK-BayesR-Predict tended to be higher than those from the existing tool BayesR.[4] In theory, BayesR should be more accurate than LDAK-BayesR-Predict, because it allows the fractions $\pi_1$, $\pi_2$, $\pi_3$ and $\pi_4$ to take any values (provided they are non-negative and sum to one), whereas LDAK-BayesR-Predict considers only 84 values for the quadruplet $(\pi_1, \pi_2, \pi_3, \pi_4)$. Therefore, we believe the lower performance of BayesR reflects that it can be difficult to achieve convergence when running MCMC. Further, LDAK-BayesR-Predict was substantially more computationally efficient than BayesR (when analyzing 20 000 individuals and 99 852 SNPs, it was 60 times faster and required 10 times less memory). The difference in runtime reflects that BayesR, by default, uses 50 000 iterations, whereas LDAK-BayesR-Predict generally uses less than 8 500 iterations (it typically takes less than 100 iterations for the variational Bayes algorithm to complete for each of the 84 training models and the final model). Further, LDAK-BayesR-Predict benefits because it estimates effect sizes for the training models concurrently, rather than consecutively (the speed-up reflects that it is faster to multiply a matrix by an 84-column matrix than to multiply the same matrix by a vector 84 times). We believe the difference in memory usage reflects that LDAK-BayesR-Predict stores genotypes more efficiently (using 1/4 bytes per value when analyzing hard genotypes, or 1 byte per value when analyzing dosage data).

**Summary statistic data tools.** We found that the accuracies of PRS from LDAK-Bolt-SS and LDAK-BayesR-SS were similar to those from the existing tools LDpred2[5] and SBayesR,[6] respectively. However, we found that the accuracies of PRS from LDAK-Lasso-SS tended to be higher than those from the existing tool lassosum,[7] while the accuracies of PRS from LDAK-Ridge-SS tended to be higher than those from the existing tools sBLUP[8] and LDpred-funct.[9] We believe the main reason why some of our summary statistic tools performed better than the corresponding existing tools, is because of our novel window-based strategy for estimating effect sizes (illustrated in Extended Data Fig. 8). We note that many existing summary statistic tools (including lassosum, LDpred and SBayesR) estimate effect sizes iteratively for whole chromosomes. For example, suppose Chromosome 1 contains 50 000 SNPs; existing tools will often update effect size estimates for SNP 1, then SNP 2, ..., then SNP 50 000, then repeat until convergence. We found that while this approach was often successful, it was hard to devise a strategy for the times it fails (i.e., when the estimated variance explained by all 50 000 SNPs does not converge). One approach is to keep the estimates from the final iteration. However, while many times these would be "sensible," leading to a good final prediction model, sometimes they would be nonsensical, leading to a very poor prediction model. An alternative is to reset estimates for all 50 000 SNPs to zero, but this would be equivalent to ignoring the whole chromosome.

This is why we chose to estimate effect sizes for small windows (by default, 1 cM). If a window fails to converge, we reset the effect sizes to their estimates prior to that window, then move to the next window. We recognize that this approach remains suboptimal; rather than skipping windows that fail to converge, it would be better to identify the problematic SNPs, exclude these and then retry the window. However, even if our strategy will, in effect, exclude all SNPs within a 1 cM window, this is better than instead excluding SNPs for a whole chromosome. Further, the problem is mitigated to some degree by the fact that we use overlapping windows (by default, windows start 1/8 cM apart). In most cases, a window will fail to converge due to SNPs in its final 1/8 cM (these are the SNPs not encountered in the previous window). Therefore, when a window fails to converge, we are, in effect, only excluding SNPs within a 1/8 cM window (because SNPs in the remainder of the window will retain their estimates from the previous window). Note that our sliding window approach is not equivalent to independently estimating effect sizes for 1 cM windows of the genome. This is because, when updating the effect size of a SNP, we consider all significant SNP-SNP correlations that include this SNP (those calculated by MegaPRS in Step 1, which by default considers pairs of SNPs within 3 cM). Therefore, by default, the estimated effect size of a SNP will be affected both by other SNPs in the 1 cM window, and SNPs outside the window but within 3 cM.

Additionally, we speculate that our summary statistic tools might benefit from using shrinkage, instead of sparsity. This is because the variational Bayes algorithm replaces current effect size estimates with the posterior mean (which is almost always non-zero). By contrast, coordinate descent (e.g., used by lassosum) update estimates with the posterior mode, which can result in many estimates being zero and might hinder convergence. However, we note that LDpred and SBayesR perform well compared to LDAK-Bolt-SS and LDAK-BayesR-SS, respectively, despite both using MCMC, which often produces many zero effect size estimates.

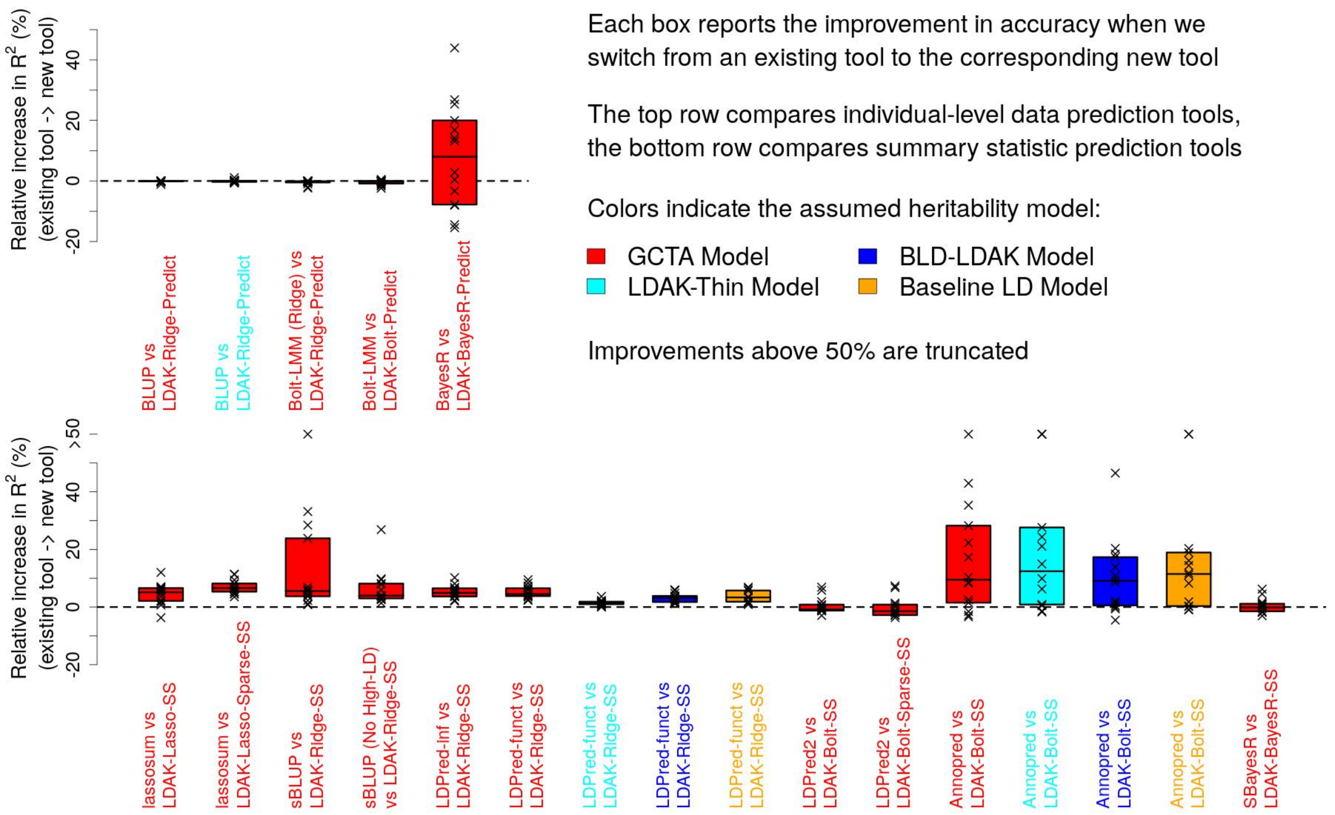


**Supplementary Figure 1: Comparing our new tools with existing tools.** We construct PRS for the first 14 UK Biobank phenotypes using our new tools and existing tools. Points report the percentage increase in $R^2$, the squared correlation between observed and predicted phenotypes across 20 000 test samples, when we switch from an existing tool to the corresponding new tool (improvements above 50% are truncated). Boxes mark the median and inter-quartile range across the 14 phenotypes; colors indicate the assumed heritability model. In all cases, our new tools perform at least as well as the corresponding existing tools.

Here we summarize the different analyses; for more details see Methods, while for scripts see Supplementary Note 1. In general, we trained prediction models using the full training data for each phenotype (200 000 individuals and 628 694 SNPs). However, this was not computationally feasible for BLUP (best linear unbiased prediction) and BayesR. Therefore, when comparing BLUP with LDAK-Ridge-Predict, we restricted to 50 000 individuals, while when comparing BayesR with LDAK-BayesR-Predict, we restricted to 20 000 individuals and 99 852 SNPs (Chromosomes 1 & 2). Further, when comparing AnnoPred with LDAK-Bolt-SS, it was necessary to exclude the major histocompatibility complex (Chr6:25-34Mb), as otherwise AnnoPred would often fail to complete. For Bolt-LMM (Ridge), we run Bolt-LMM with the options `-pEst .5 -varFrac2Est .5` (i.e., forcing the ridge regression model). For sBLUP (No High-LD), we run sBLUP excluding regions of long-range linkage disequilibrium. Note that there is no need to compare lasso-based tools that use individual-level data, because the best existing tool of this type is the original version of big_spLinReg (i.e., our new version run assuming the GCTA Model).

We compared LDpred-funct with LDAK-Ridge-SS. Strictly, this comparison is not fair (LDAK-Ridge-SS is disadvantaged), because LDpred-funct uses a regularized version of ridge regression (the effect size estimates from the ridge regression model are regularized via cross-validation).[9] However, we observed that this regularization made little difference to accuracy (we found that results from LDpred-funct were very similar to those from LDpred-inf, which omits the regularization). We first compared LDpred2 with LDAK-Bolt-SS. However, the two tools use slightly different effect size prior distribution forms; LDpred2 assumes $\beta_j \; p\mathbb{N}(0, \sigma^2) + (1-p)\delta_{\{0\}}$, where $\delta_{\{0\}}$ is a point mass at zero, while LDAK-Bolt-SS assumes $\beta_j \; p\mathbb{N}(0, (1-f_2)/p\sigma^2) + (1-p)N(0, f_2/(1-p)\sigma^2)$. Therefore, we also compared LDpred2 with LDAK-Bolt-Sparse-SS (see Supplementary Table 8), which sets $f_2 = 0$, so that the prior distribution form matches that of LDpred2. Although lassosum and LDAK-Lasso-SS use the same prior distribution form, their algorithms differ substantially. Therefore, we also compared lassosum with LDAK-Lasso-Sparse-SS (see Supplementary Table 8), whose algorithm is more similar to that of lassosum.

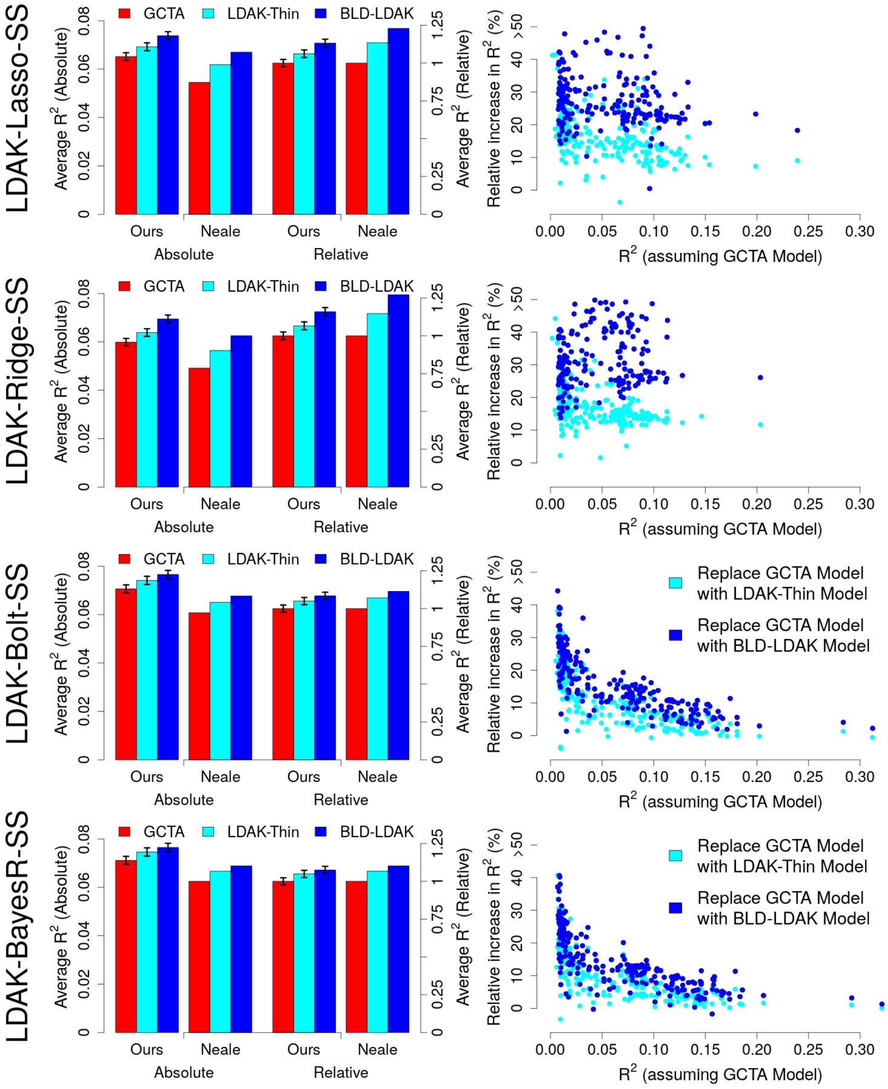


**Supplementary Figure 2: Impact of changing the heritability model when using summary statistics.** We construct PRS for the 225 UK Biobank phenotypes using LDAK-Lasso-SS (top row), LDAK-Ridge-SS (second row), LDAK-Bolt-SS (third row) and LDAK-BayesR-SS (bottom row), assuming the GCTA, LDAK-Thin or BLD-LDAK Model. We measure the accuracy of PRS via $R^2$, the squared correlation between observed and predicted phenotypes.

For the majority of phenotypes, we do not have separate training and test data, and therefore it is necessary to generate pseudo training and test summary statistics (see Methods); we then use the pseudo training summary statistics to train prediction models, and the pseudo test summary statistics to test them. The first column focuses on the first 14 phenotypes, for which we have access to individual-level data. Boxes report, for each heritability model, average $R^2$ (first six boxes) and average $R^2$ relative to the GCTA Model (last six boxes), calculated using either our individual-level data (for which we use separate training and test data) or summary statistics from the Neale Lab (for which we use pseudo training and test summary statistics). Vertical segments report 95% confidence intervals (to calculate these require individual-level data). We see that estimates of $R^2$ tend to be higher for PRS calculated using our data than for PRS calculated using Neale Lab summary statistics, likely reflecting that we performed more careful quality control. However, the relative estimates are similar whether we use our data or Neale Lab summary statistics, indicating that it is valid to compare heritability models using pseudo summary statistics. Note that we provide further support for the use of pseudo summary statistics in Supplementary Fig. 13.

The middle column considers all 225 phenotypes (using estimates of $R^2$ from pseudo summary statistics). The $x$-axis reports $R^2$ when assuming the GCTA Model, while the $y$-axis reports the percentage increase in $R^2$ if we instead assume the LDAK-Thin or BLD-LDAK Model (improvements above 50% are truncated). The last column provides the same results, except that phenotypes are grouped based on $R^2$ when assuming the GCTA Model (boxes mark the median and inter-quartile range).

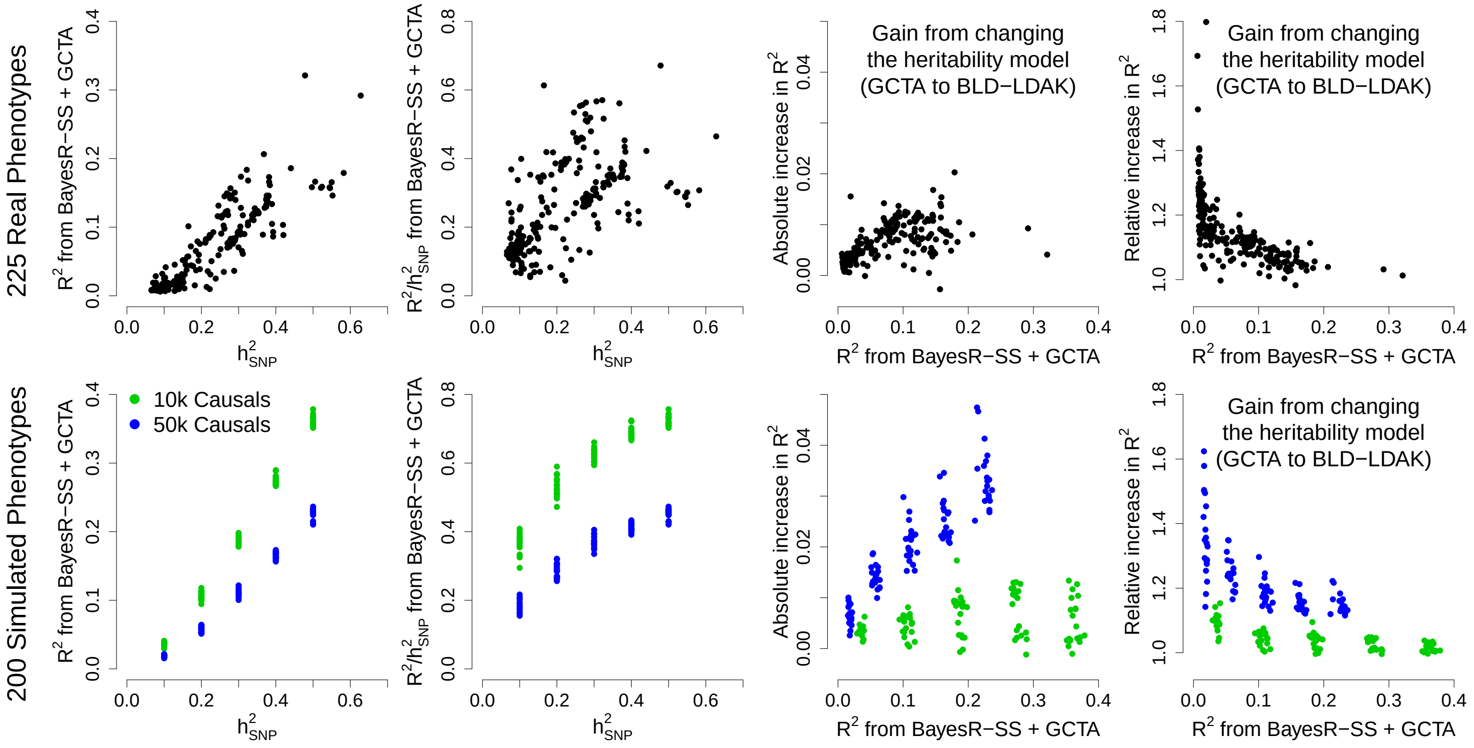


**Supplementary Figure 3: Impact of improving the heritability model for simulated data.** Our analyses of real data have shown that by improving the heritability model, we can substantially improve the accuracy of PRS for a wide variety of phenotypes. These include continuous, binary and ordinal phenotypes, that have low, medium and high SNP heritability, and that are both closely and distantly related to diseases. When we divided phenotypes based on $R^2$ (the squared correlation between observed and predicted phenotypes), we observed that the absolute advantage of our new tools was largest for traits with higher $R^2$, while the relative advantage of our new tools was largest for traits with lower $R^2$. This figure shows that these two trends are similar to those we observe in simulations.

Each panel shows how the accuracy of PRS constructed using LDAK-BayesR-SS changes when we switch from the GCTA Model to the BLD-LDAK Model. We measure accuracy as $R^2$, the squared correlation between observed and predicted phenotypes across 20 000 test individuals. The top row reports results when analyzing the 225 real phenotypes from the UK Biobank, while the bottom row reports results when analyzing 200 simulated phenotypes (constructed using UK Biobank genotypes). To generate each simulated phenotype, we randomly selected either 10 k (green points) or 50 k (blue points) of the 628 k directly-genotyped SNPs to be causal, then sampled their effect sizes consistent with the BLD-LDAK Model. Specifically, if SNP $j$ was picked to be causal, we sampled $\beta_j$ from $N(0, e_j)$, where $e_j$ is the estimate of $\mathbb{E}[h_j^2]$ from our analysis of height assuming the BLD-LDAK Model. Having generated the genetic effects of individuals (i.e., calculated $\sum_j X_j\beta_j$), we then added Gaussian-distributed noise so that the heritability of each phenotype was 0.1, 0.2, 0.3, 0.4 or 0.5. The scripts we used to produce these simulated phenotypes are provided in Supplementary Note 1.

The first two columns show that for both real and simulated phenotypes, $R^2$ and $R^2/h^2_{\text{SNP}}$ tend to be higher for phenotypes with higher $h^2_{\text{SNP}}$. These patterns reflect that a phenotype with higher $h^2_{\text{SNP}}$ will, all other things being equal, be easier to predict than one with lower $h^2_{\text{SNP}}$ (e.g., the more heritable phenotype will tend to have larger standardized effect sizes). The last two columns show that the absolute increase in $R^2$ due to improving the heritability model tends to be higher for phenotypes with higher $R^2$, while the relative increase in $R^2$ due to improving the heritability model tends to be higher for phenotypes with lower $R^2$. These patterns reflect that for phenotypes that are hard to predict (e.g., those with low heritabilities or very complex genetic architectures), there is a large benefit to improving the prior assumptions. By contrast, for phenotypes that are easy to predict, we can achieve relatively high $R^2$ even with suboptimal prior assumptions, meaning that there is less advantage using improved heritability models.

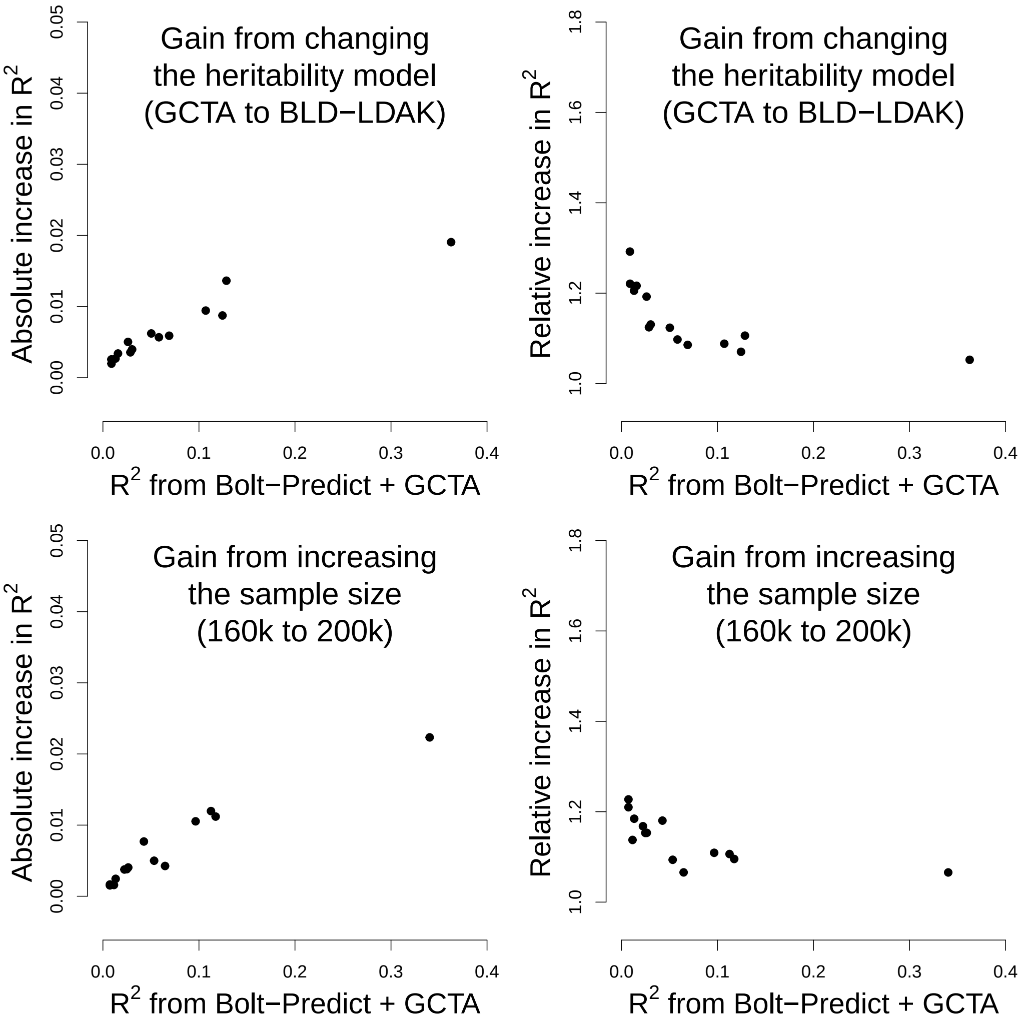


**Supplementary Figure 4: Impact of increasing the training sample size.** Our analyses of real data have shown that by improving the heritability model, we can substantially improve the accuracy of PRS for a wide variety of phenotypes. These include continuous, binary and ordinal phenotypes, that have low, medium and high SNP heritability, and that are both closely and distantly related to diseases. When we divided phenotypes based on $R^2$ (the squared correlation between observed and predicted phenotypes), we observed that the absolute advantage of our new tools was largest for traits with higher $R^2$, while the relative advantage of our new tools was largest for traits with lower $R^2$. This figure shows that these two trends are similar to those we observe when we improve prediction accuracy by increasing the sample size.

We restrict to the first 14 UK Biobank phenotypes (those for which we have individual-level data), and construct PRS using LDAK-Bolt-Predict. We see that the gains in prediction accuracy when we switch from the GCTA Model to the BLD-LDAK Model (top two plots) are similar to the gains when we increase the number of training individuals from 160 k to 200 k (bottom two plots). This is the case both when we measure the absolute increase in $R^2$ (left plots) and the relative increase in $R^2$ (right plots).

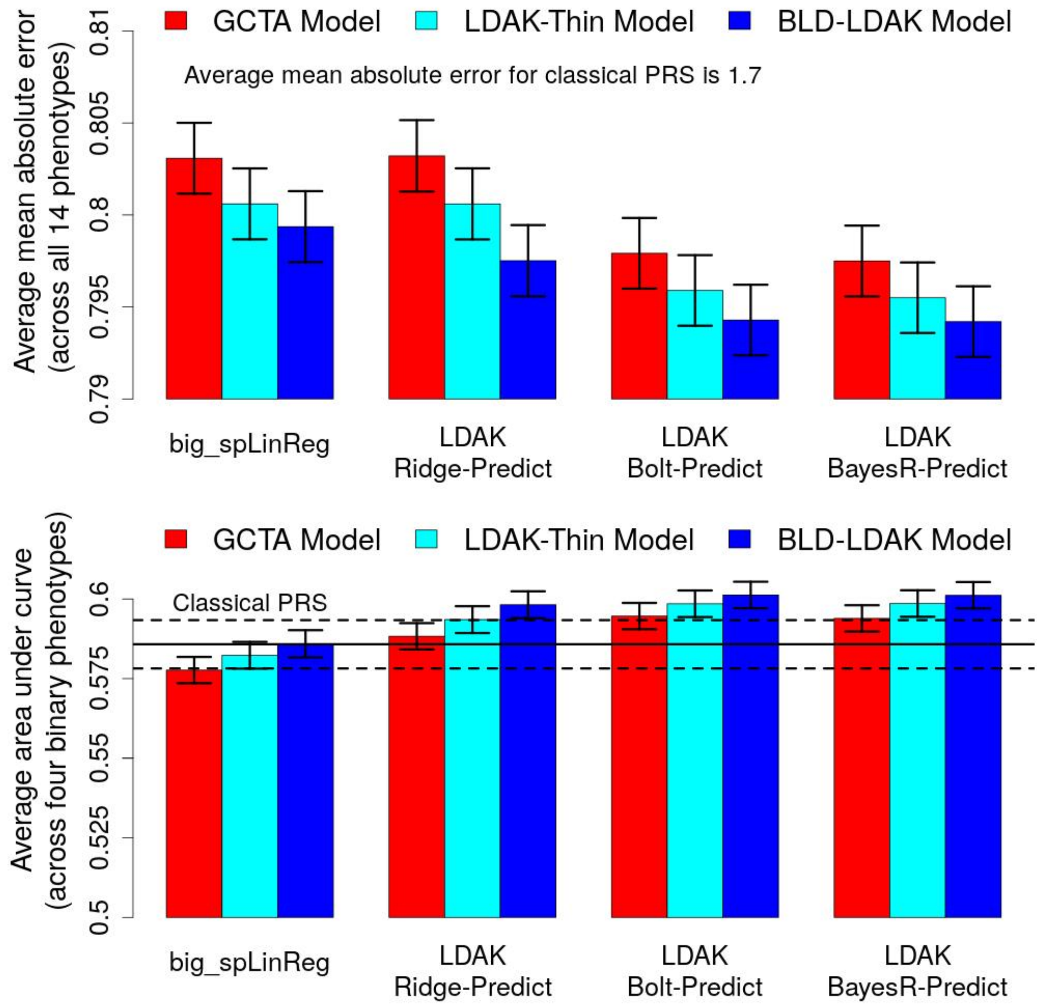


**Supplementary Figure 5: Alternative measures of prediction accuracy.** We use our four individual-level data tools to construct PRS for the first 14 UK Biobank phenotypes (using all 200 000 training samples). We measure the accuracy of each PRS across 20 000 test samples; bars report the average accuracy across phenotypes (vertical segments mark 95% confidence intervals). Colors indicate the assumed heritability model, while blocks indicate the prediction tool. The top row reports the mean absolute error between observed and predicted phenotypes, averaged across all 14 phenotypes. The average mean absolute error for classical PRS is 1.71 (s.d. 0.003). The bottom row reports the area under the receiver operating curve, averaged across the four binary phenotypes (area under curve can only be computed for binary phenotypes). The horizontal lines mark average area under curve for classical PRS and a 95% confidence interval.

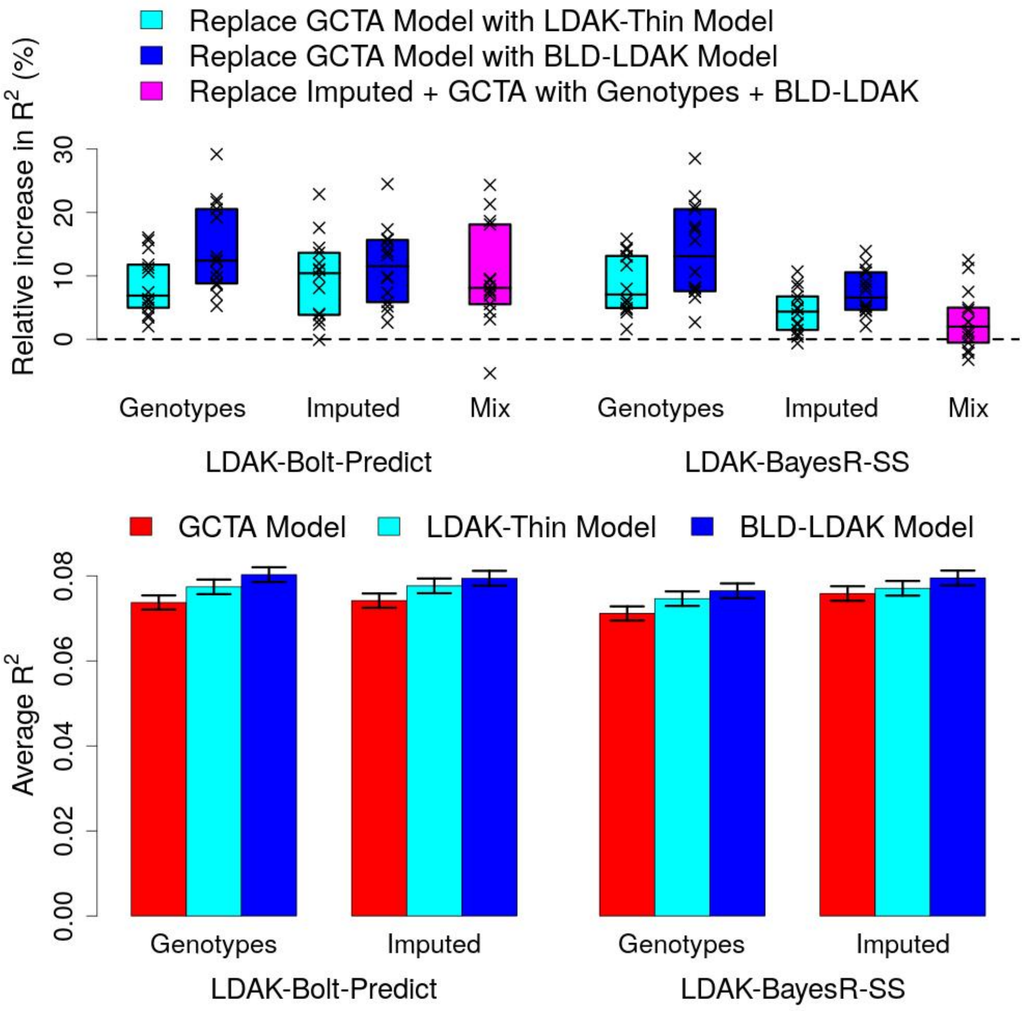


**Supplementary Figure 6: Including imputed SNP genotypes.** We use LDAK-Bolt-Predict and LDAK-BayesR-SS to construct PRS for the first 14 UK Biobank phenotypes (using all 200 000 training samples). First we restrict to 629,000 directly-genotyped SNPs (the same as for our main analyses), then we increase the number of SNPs to 7.5M by including imputed genotypes. We measure the accuracy of each PRS via $R^2$, the squared correlation between observed and predicted phenotypes across 20 000 test samples. When using LDAK-Bolt-Predict and including imputed genotypes, it was not computationally feasible to analyze all SNPs together, so we instead analyzed each chromosome separately. Note that we merged effect size estimates across chromosomes before performing cross-validation, so that we continued to select prior parameters based on genome-wide data (rather than separately for each chromosome).

The top row shows the improvement in prediction accuracy for individual phenotypes. For the light and dark blue boxes, points report the percentage increase in $R^2$ when each tool is switched from assuming the GCTA Model to either the LDAK-Thin or BLD-LDAK Model (boxes mark the median and inter-quartile range across the 14 phenotypes). We see that when using imputed data, $R^2$ increases when we improve the heritability model, similar to when using directly-genotyped data. For the purple boxes, points report the percentage increase in $R^2$ when each tool is switched from assuming the GCTA Model and using imputed data, to assuming the BLD-LDAK Model and using genotyped data (boxes mark the median and inter-quartile range across the 14 phenotypes). We see that the improvement in accuracy by switching from the GCTA Model to the BLD-LDAK Model is generally larger than the improvement in accuracy by switching from directly-genotyped to imputed SNPs. In the bottom row, bars report $R^2$ averaged across the 14 phenotypes (vertical segments mark 95% confidence intervals). Colors indicate the assumed heritability model.

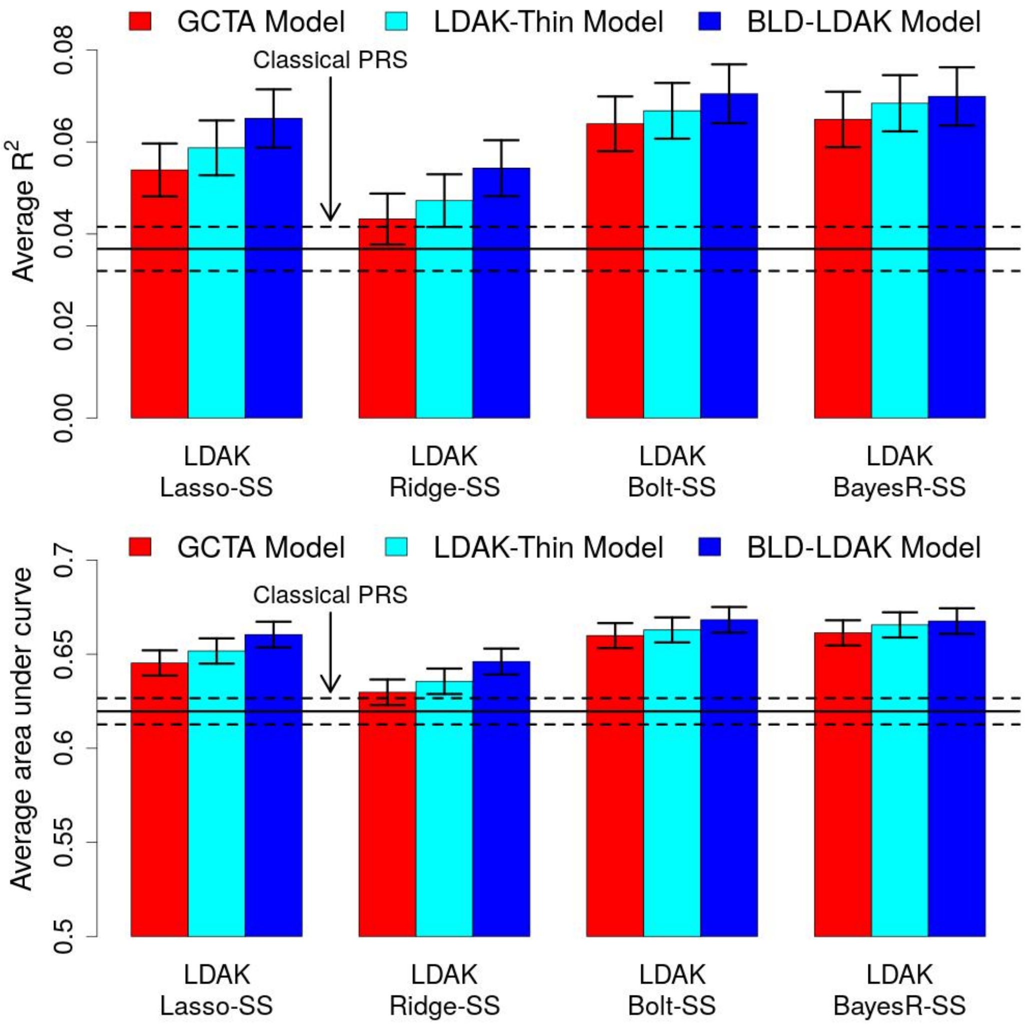


**Supplementary Figure 7: Eight additional diseases.** We use our four summary statistic tools to construct PRS for asthma, atrial fibrillation, breast cancer, inflammatory bowel disease, prostate cancer, rheumatoid arthritis, schizophrenia and type 2 diabetes, using summary statistics from published studies (that did not use UK Biobank data). We then tested the accuracy of the PRS using UK Biobank individuals. In the top row, bars report $R^2$, the squared correlation between observed and predicted phenotypes, averaged across the eight diseases (vertical segments mark 95% confidence intervals). Colors indicate the assumed heritability model. In the bottom row, bars report area under the receiver operating curve, averaged across the eight diseases (vertical segments mark 95% confidence intervals). Colors indicate the assumed heritability model.

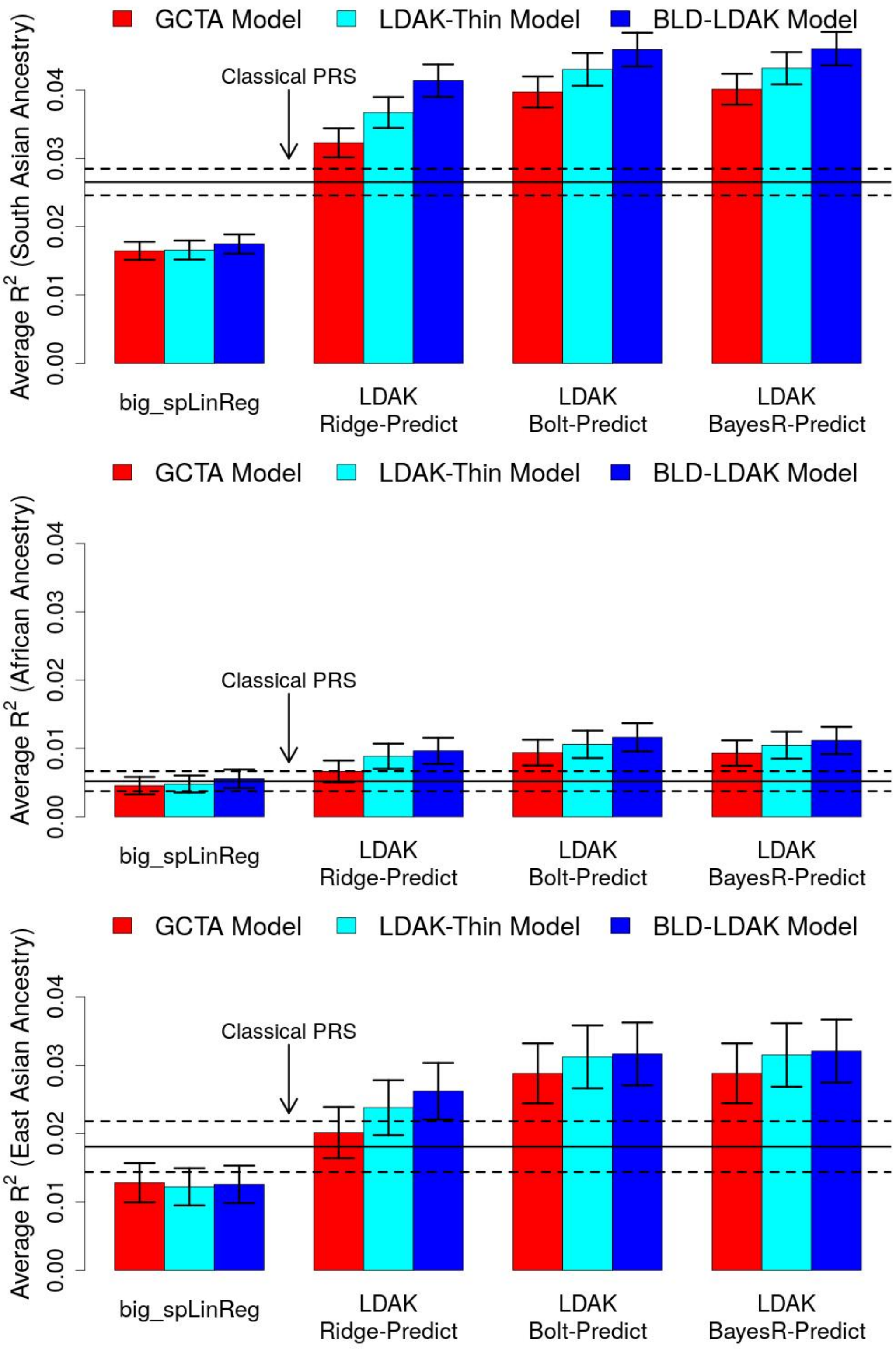


**Supplementary Figure 8: Cross-ancestry prediction.** As part of our main analyses, we used our four individual-level data tools to construct prediction models for the first 14 UK Biobank phenotypes. These models were constructed using data from 200 000 white British individuals. Here we test how well they predict phenotypes for individuals inferred to have South Asian (top), African (middle) and East Asian (bottom) ancestry. Bars report $R^2$, the squared correlation between observed and predicted phenotypes, averaged across the 14 phenotypes (vertical segments mark 95% confidence intervals). Colors indicate the assumed heritability model, while blocks indicate the prediction tool. The horizontal lines mark average $R^2$ for classical PRS and a 95% confidence interval.

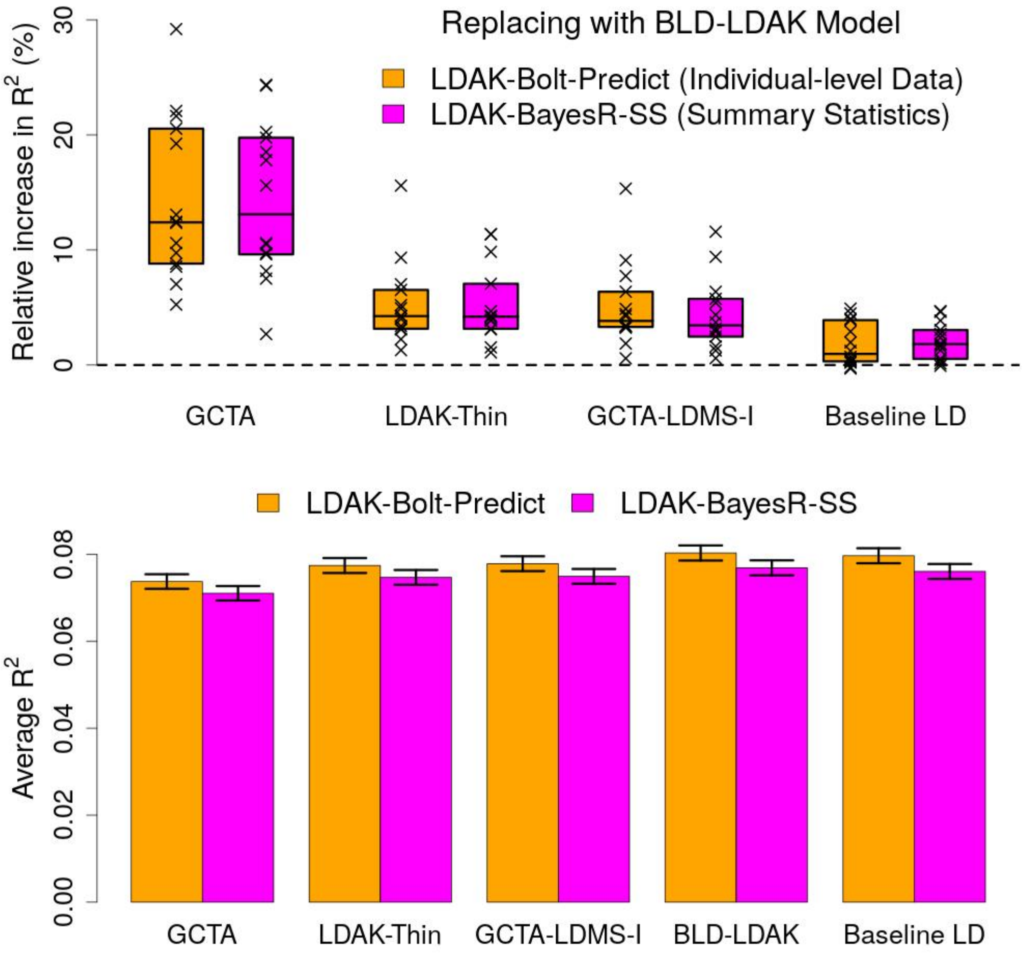


**Supplementary Figure 9: Alternative heritability models.** We use LDAK-Bolt-Predict and LDAK-BayesR-SS to construct PRS for the first 14 UK Biobank phenotypes (using all 200 000 training samples). We measure the accuracy of each PRS via $R^2$, the squared correlation between observed and predicted phenotypes across 20 000 test samples. In addition to the GCTA, LDAK-Thin and BLD-LDAK Models, we also consider the GCTA-LDMS-I[10] and Baseline LD Model,[11] the models recommended by the authors of GCTA[12] and LD Score Regression,[13] respectively. In the top row, points report the percentage increase in $R^2$ for individual phenotypes when we switch from assuming the GCTA, LDAK-Thin, GCTA-LDMS-I or Baseline LD Model to the BLD-LDAK Model (boxes mark the median and inter-quartile range across the 14 phenotypes). In the bottom row, bars report $R^2$ averaged across the 14 phenotypes for each heritability model (vertical segments mark 95% confidence intervals).

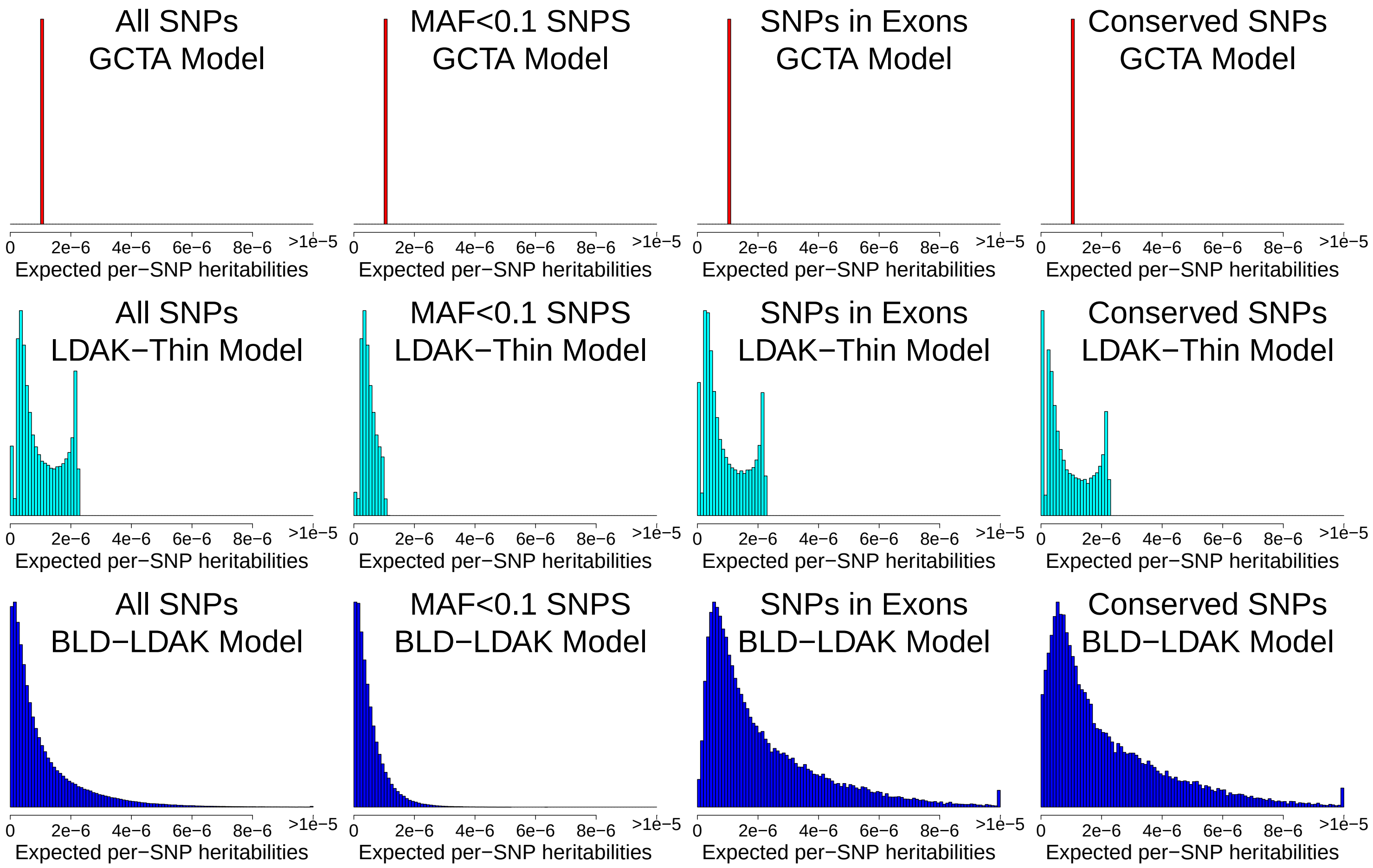


**Supplementary Figure 10: Comparison of heritability models.** Prior to running out new prediction tools, we used SumHer to obtain estimates of $\mathbb{E}[h_j^2]$, the expected heritability contributed by each SNP, given the heritability model.[3] Our main analyses considered three heritability models: the GCTA, LDAK-Thin and BLD-LDAK Models. This figure seeks to explain why improving the heritability model (i.e., switching from the GCTA to the LDAK-Thin Model, or from the LDAK-Thin to the BLD-LDAK Model) resulted in more accurate prediction models. The top, middle and bottom rows report estimates of $\mathbb{E}[h_j^2]$ from our analysis of height when assuming the GCTA, LDAK-Thin and BLD-LDAK Models, respectively; Columns 1, 2, 3 & 4 consider estimates across the whole genome, SNPs with MAF<0.1, SNPs in exons and SNPs in conserved regions, respectively.

The GCTA Model assumes that $\mathbb{E}[h_j^2]$ is constant, and therefore the estimates of $\mathbb{E}[h_j^2]$ are the same for all SNPs. The LDAK-Thin Model assumes that SNPs with lower MAF have smaller $\mathbb{E}[h_j^2]$. This relationship is evident in the second column, which reports estimates of $\mathbb{E}[h_j^2]$ for SNPs with MAF<0.1, and is consistent with selection causing SNPs that have a larger influence on the phenotype to have smaller MAF.[14,15] The BLD-LDAK Model generalizes the LDAK-Thin Model by allowing $\mathbb{E}[h_j^2]$ to vary both according to MAF and 65 SNP annotations; six of these annotations are related to linkage disequilibrium, while the remaining 59 are functional classifications (see Supplementary Table 7 for full details). Two of the most impactful functional classifications are whether a SNP is exonic or within a conserved region; the third and fourth columns show that for exonic and conserved SNPs, respectively, estimates of $\mathbb{E}[h_j^2]$ tend to be higher than average.[15]

Our analyses of real phenotypes repeatedly found that PRS constructed assuming the LDAK-Thin Model outperformed those constructed assuming the GCTA Model, while PRS constructed assuming the BLD-LDAK Model performed better still. These results confirm that SNPs with higher MAF tend to have larger effect sizes than SNPs with lower MAF (and hence prediction tools benefit from giving more weight to SNPs with higher MAF). Moreover, they confirm that there are functional categories of SNPs, such as exons or conserved regions, that are enriched for heritability (and therefore prediction tools benefit from giving more weight to SNPs in these categories).

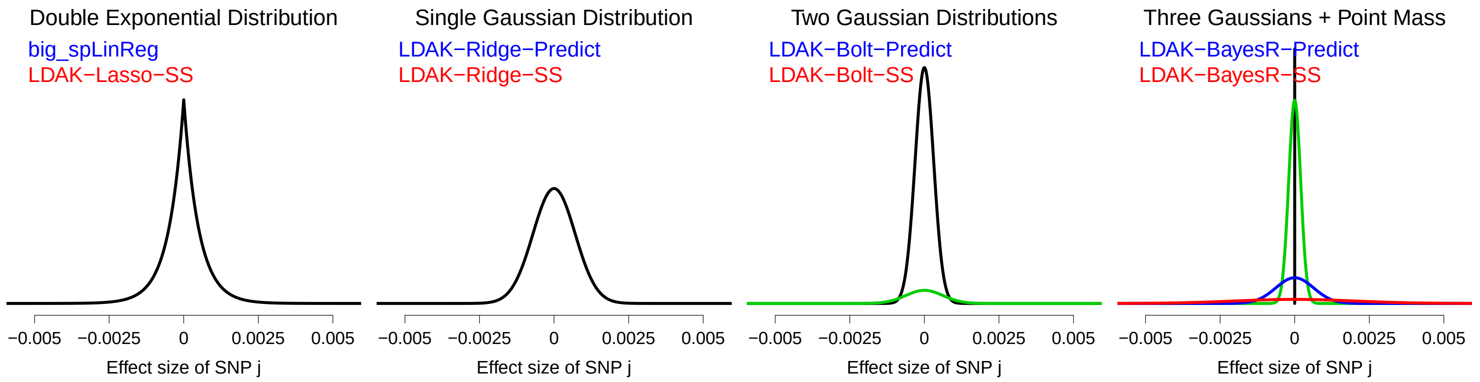


**Supplementary Figure 11: Comparison of prior distribution forms.** When analyzing individual-level data, we recommend using LDAK-Bolt-Predict, while when analyzing summary statistics, we recommend using LDAK-BayesR-SS (in both cases, assuming the BLD-LDAK Model). This is because these prediction tools performed best across a wide variety of phenotypes. While there were individual phenotypes for which an alternative tool performed better, the difference was always slight and never significant ($P > 0.5$ from a one-sided Wald Test). Here we seek to provide a biological justification for the different prediction tools. Please note that this section is for interest only, and we do not advise it is used to decide which tool to use. Instead, if you have a phenotype for which you are concerned our recommendations are inadequate, we advise you use cross-validation (e.g., first construct PRS using a variety of tools based on 90% of individuals, then measure the accuracy of these PRS using the remaining 10% of individuals, and finally select the tool corresponding to the most accurate PRS).

LDAK-Ridge-Predict and LDAK-Ridge-SS use a single Gaussian distribution. This is an “infinitesimal model," in which every SNP is assumed to influence the phenotype (i.e., an omnigenic model of inheritance). The use of a single Gaussian distribution has been criticized on account of its “thin tails" (i.e., it considers very unlikely the possibility of SNPs with large effects).[16] In particular, the distribution tends to perform poorly for phenotypes with one or more loci of large effect (e.g., loci that explain more than 1% of phenotypic variance). big_spLinReg and LDAK-Lasso-SS use a double exponential distribution. Compared to the Gaussian distribution, this has thicker tails, and therefore can perform better for phenotypes where substantial heritability is concentrated in a few loci. Moreover, tools that use the double exponential distribution will generally produce sparse solutions (most effect size estimates are zero), which can be desirable for phenotypes considered to have low polygeneity (note that while big_spLinReg produces sparse solutions, LDAK-Lasso-SS does not, because its effect size estimates are posterior means instead of posterior modes).

In recent years, it has become popular to use mixture priors, that allow for subsets of SNPs to have distinct effect size distributions.[1,4,17] LDAK-Bolt-Predict and LDAK-Bolt-SS use two Gaussian distributions, that differ in their variances. The Gaussian distribution with the larger variance is designed to capture the contributions of the (relatively few) SNPs with larger effect sizes, while the other Gaussian distribution is designed to capture the contributions of the SNPs with smaller effect sizes. Lastly, LDAK-BayesR-Predict and LDAK-BayesR-SS use three Gaussian distributions, with small, medium and large variances, as well as a point mass at zero. The point mass allows for some SNPs to have zero effect (i.e., to not contribute towards the phenotype), while the Gaussian distributions with small, medium and large variances are designed to capture the contributions of SNPs with small, medium and large effect sizes, respectively.

In our analyses, we found that LDAK-Bolt-Predict, LDAK-BayesR-Predict and LDAK-BayesR-SS performed best. Each of these tools uses a mixture prior, and thus their superior performances reflect that it is difficult to describe the distribution of effect sizes across the genome using a single prior distribution. It may appear a contradiction that LDAK-Bolt-Predict and LDAK-BayesR-Predict have similar performance, considering the former assumes that all SNPs influence the phenotype, whereas the latter assumes that a substantial proportion of SNPs have zero effect. Instead, this suggests that it often suffices (at least when constructing prediction models) to assume the SNPs with smallest influence on the phenotype have effect size zero (or conversely to assume the SNPs that do not influence the phenotype have very small effect sizes).

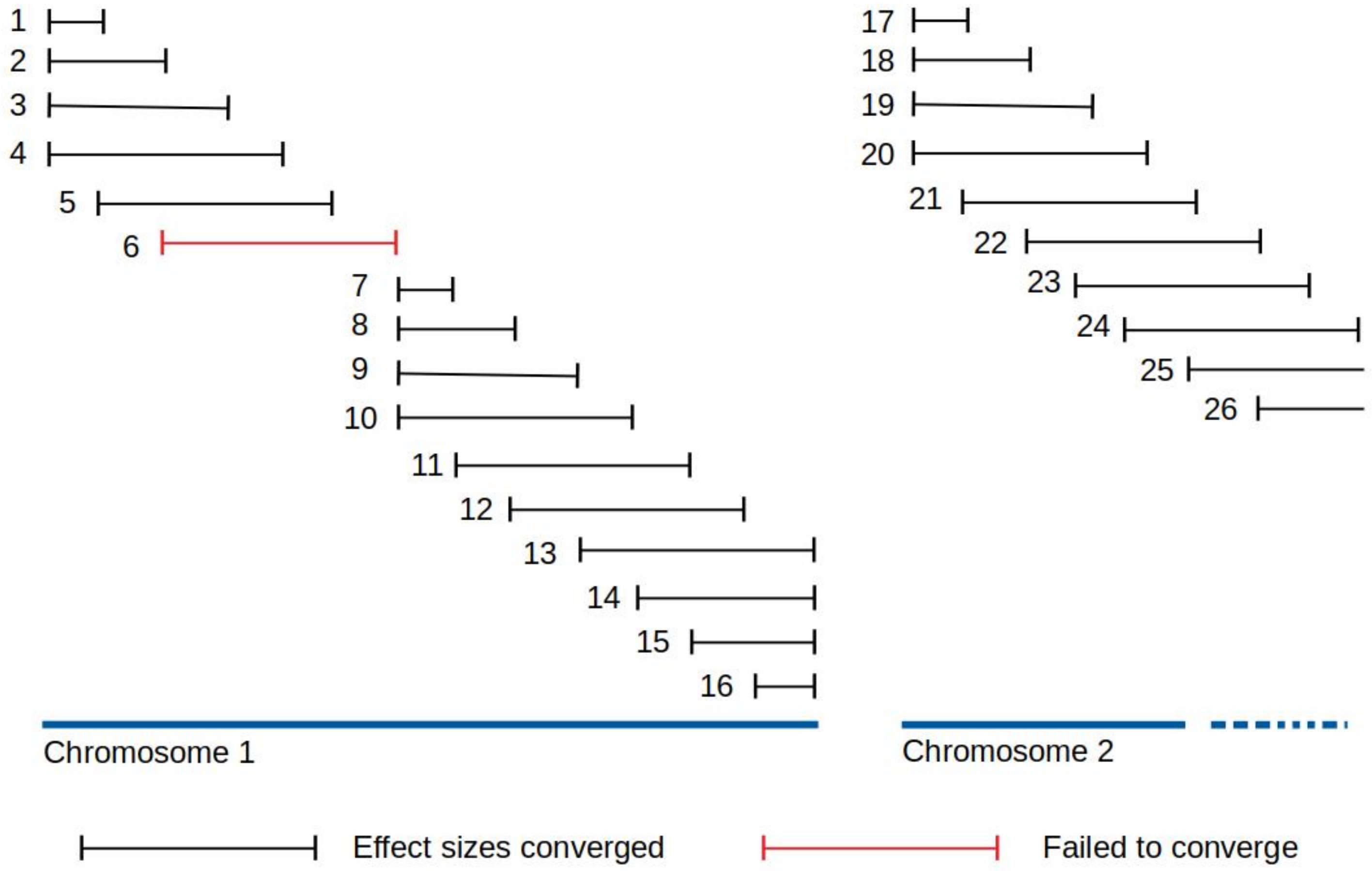


**Supplementary Figure 12: A sliding window approach for estimating effect sizes.** In Methods, we explain how our summary statistic prediction tools estimate effect sizes iteratively using variational Bayes. For this, we found it was not feasible to iterate over all SNPs in the genome. Therefore, our tools instead use sliding windows, illustrated above (numbers indicate the order in which windows are processed). By default, we iteratively estimate effect sizes for all SNPs in a 1 cM window, stopping when the estimated proportion of variance explained by these SNPs changes by less than $10^{-5}$. We then move 1/8 cM along the genome and repeat for the next 1 cM window. If a window fails to converge within 50 iterations, we reset the effect sizes of its SNPs to their values prior to considering that window, then start the next window at the end of the current window.

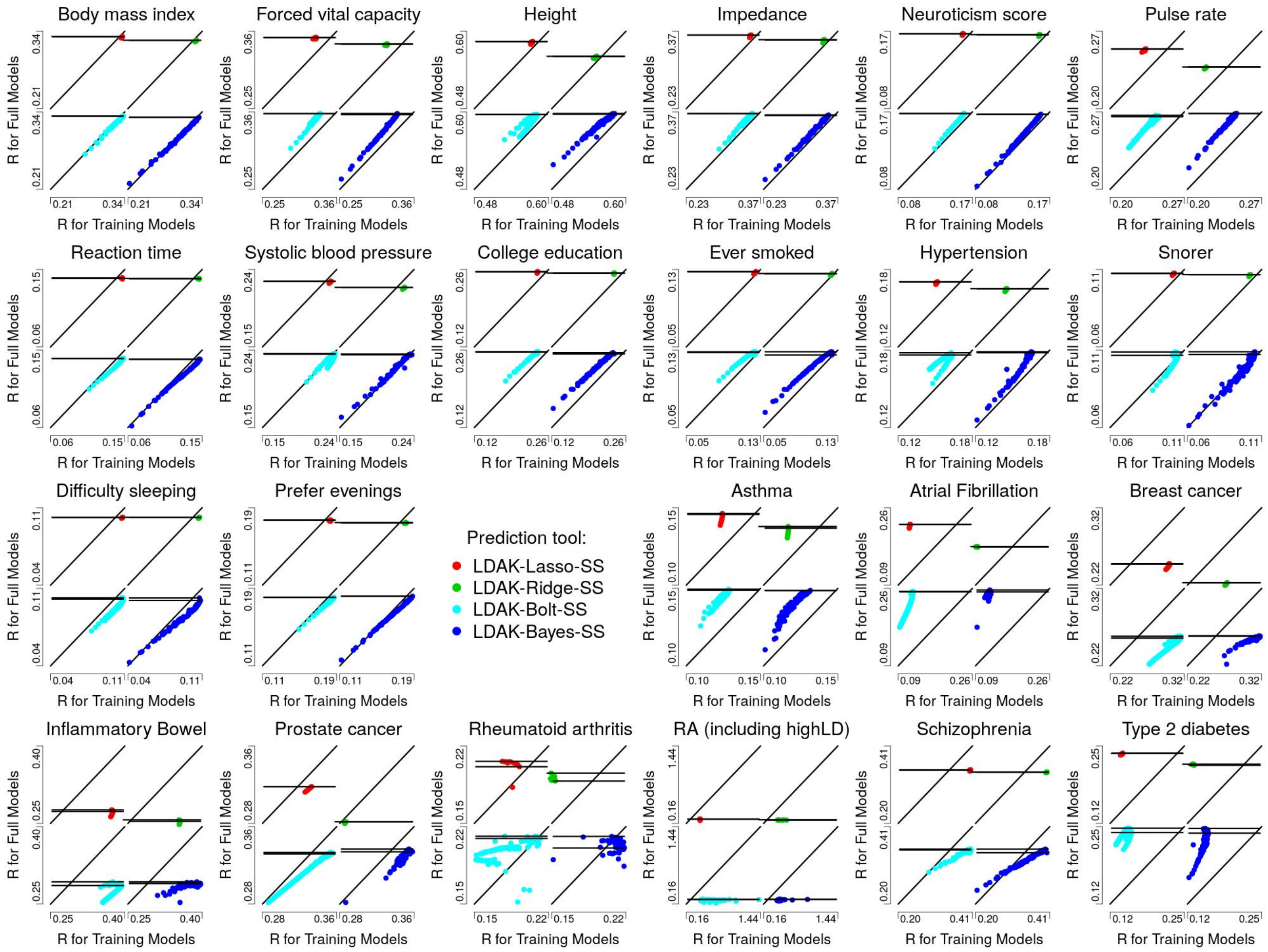


**Supplementary Figure 13: Validity of pseudo summary statistics.** Points compare estimates of $R$ for training models (computed using pseudo summary statistics) and full models (computed using independent data). For each phenotype, we consider separately the pairs of models constructed by LDAK-Lasso-SS (top left plots), LDAK-Ridge-SS (top right plots), LDAK-Bolt-SS (bottom left plots) and LDAK-BayesR-SS (bottom right plots). In each plot, the diagonal line marks $y = x$, while the two horizontal lines mark $R$, respectively, for the best full model and for the full model corresponding to the best training model. Ideally, these lines would coincide, meaning that the prior distribution parameters chosen via cross-validation always result in the most accurate final model. However, in practice, this is unrealistic, because we are using independent datasets to measure the accuracy of training and full models, and there is noise in the estimates of $R$.

The first 14 panels show that for the first 14 UK Biobank phenotypes, there is close concordance between estimates of $R$ for training and full models, and that the most accurate training model generally corresponds to one of the best full models. In general, the two horizontal lines either perfectly coincide or overlap, indicating that we can effectively perform cross-validation using pseudo summary statistics. The last nine panels consider the 8 additional diseases, for which we constructed PRS using summary statistics from studies independent of UK Biobank, then tested them using UK Biobank data. Note that we expect larger differences in estimates of $R$ between training and full models, reflecting that there are differences in disease definitions, case/control ratios and quality control between the published studies and our UK Biobank data. Nonetheless, in most cases there is close concordance between the two horizontal lines.

We have observed that the accuracy of estimates of $R$ can reduce when there are loci of strong effect within regions of long-range linkage disequilibrium (LD). Therefore, as a safety precaution, we recommend always excluding long-range LD regions (approximately 3% of the genome) when estimating $R$ for training models. In the above figure, we generally exclude long-range LD regions when estimating $R$ for training models (but not when estimating $R$ for full models). However, for rheumatoid arthritis, we also report estimates of $R$ for training models when long-range LD regions are retained (Panels 3 & 4 on the bottom row). For this disease, a single SNP within the major histocompatibility complex explains 2% of phenotypic variation. We see that estimates of $R$ are unreliable when long-range LD regions are retained, but that they become reasonable when long-range LD regions are excluded.

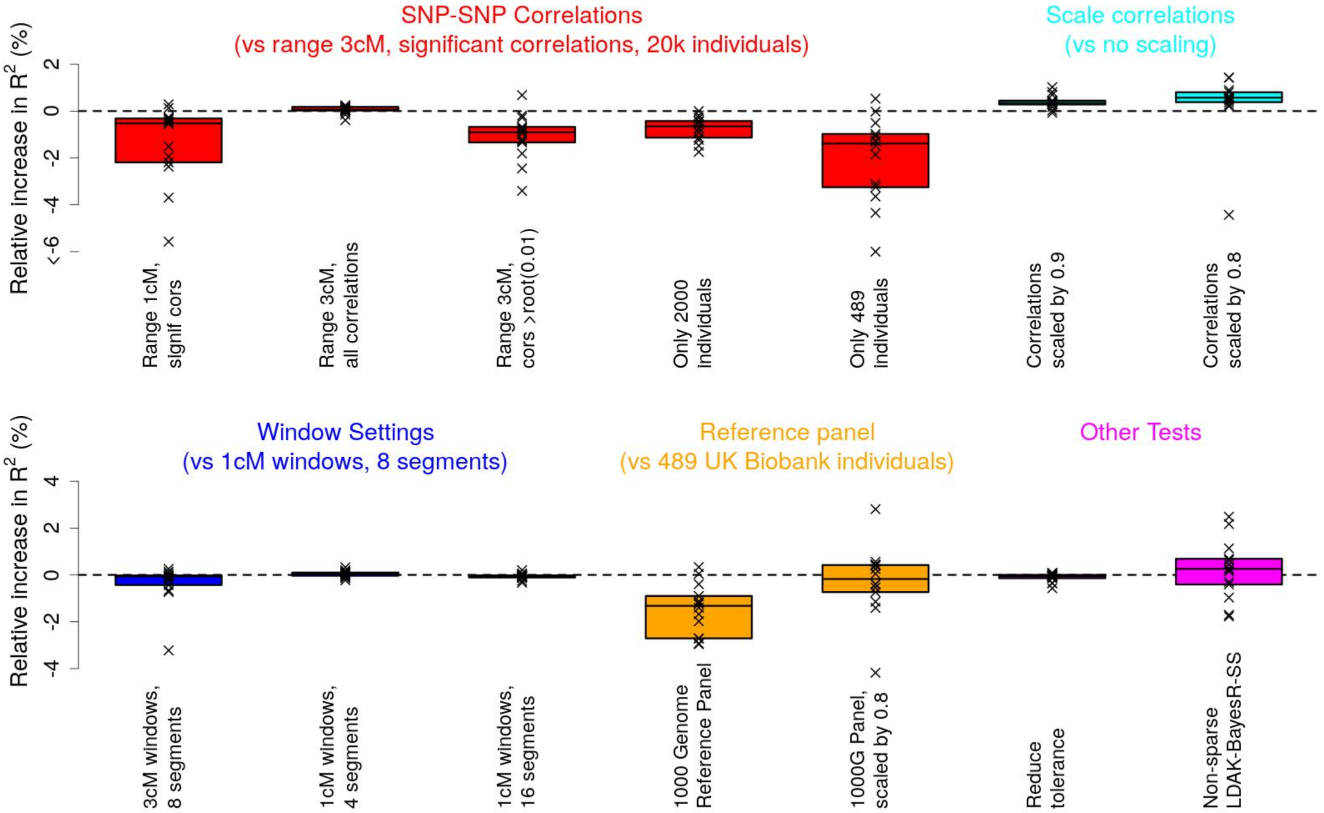


**Supplementary Figure 14: Sensitivity of MegaPRS to setting choices.** LDAK-BayesR-SS is contained within MegaPRS (see Methods). Here we investigate the impact of changing settings of MegaPRS from their defaults and of using different reference panels. We use LDAK-BayesR-SS to construct PRS for the first 14 UK Biobank phenotypes (using all 200 000 training samples). Points report the percentage increase in $R^2$, the squared correlation between observed and predicted phenotypes across 20 000 test samples, when we change settings (boxes mark the median and inter-quartile range across the 14 phenotypes).

For the five red boxes, we change settings when calculating SNP-SNP correlations: by default, MegaPRS records correlations within 3 cM that are significant (P<0.01 from a two-sided likelihood ratio test, which when the reference panel contains 20 000 individuals, corresponds to correlations with magnitude greater than $\sqrt{0.0003}$). Here we instead record correlations within 1 cM, or record both significant and non-significant correlations, or record those whose magnitude is greater than $\sqrt{0.01}$, or reduce the reference panel to 2000 individuals, or reduce the reference panel to 489 individuals. For the two light-blue boxes, we scale the estimates of SNP-SNP correlations by 0.9 or 0.8 (instead of not scaling). For the three dark-blue boxes, we change settings when estimating effect sizes: by default, MegaPRS estimates effect sizes for a 1 cM window, then moves 1/8th of a window along the genome and repeats; here we instead use a 3 cM window, or move 1/4th of a window along the genome, or move 1/16th of a window along the genome. For the first orange box, we replace the UK Biobank reference panel with 489 Europeans individuals from the 1000 Genome Project[18] (here we compare results with those obtained using a reference panel of only 489 UK Biobank individuals); for the second orange box, we do the same except scale estimates of SNP-SNP correlations by 0.8. For the first purple box, we change the convergence tolerance to $10^{-6}$ (instead of $10^{-5}$). For the second purple box, we use a non-sparse version of LDAK-BayesR-SS (in its effect size prior distribution, we replace the point mass at zero with a Gaussian distribution with variance $\sigma^2$/1000).

Overall, we find that the performance of LDAK-BayesR-SS is fairly robust to changing settings. The largest impact is if we replace the UK Biobank reference panel with a 1000 Genomes Project panel. In this case, $R^2$ reduces on average by about 2% due to reducing the number of individuals from 20 000 to 489 (fifth box on the top row) and on average by about a further 1% due to replacing UK Biobank genotypes with 1000 Genome Project genotypes (fourth box of the bottom row). We note that there is a small advantage if we shrink estimates of SNP-SNP correlations (last two boxes on the top row), and that by shrinking, we can offset some of the reduction due to substituting in the 1000 Genome Project reference panel (fifth box on the bottom row).

| | | | | | | Accuracy of LDAK-Bolt-Predict PRS (assuming the BLD-LDAK Model) | | | | | |
|---|---|---|---|---|---|---|---|---|---|---|---|
| | | $h^2_{\text{SNP}}$ (SD) | | | Inflation | 100 k samples | | 200 k samples | | 400 k (predicted) | |
| **Phenotype** | $h^2$ **(SD)** | **GCTA** | **LDAK-Thin** | **BLD-LDAK** | **(SD)** | $R^2$ **(SD)** | $R^2/h^2_{\text{SNP}}$ **(SD)** | $R^2$ **(SD)** | $R^2/h^2_{\text{SNP}}$ **(SD)** | $R^2$ | $R^2/h^2_{\text{SNP}}$ |
| Body mass index | 0.29 (0.01) | 0.34 (0.01) | 0.31 (0.01) | 0.30 (0.01) | 0.000 (0.003) | 0.08 (0.00) | 0.27 (0.01) | 0.12 (0.00) | 0.39 (0.01) | 0.17 | 0.58 |
| Forced vital capacity | 0.35 (0.02) | 0.35 (0.01) | 0.31 (0.01) | 0.30 (0.01) | 0.000 (0.003) | 0.10 (0.00) | 0.32 (0.01) | 0.13 (0.00) | 0.44 (0.02) | 0.19 | 0.63 |
| Height | 0.68 (0.02) | 0.68 (0.03) | 0.63 (0.03) | 0.61 (0.02) | 0.001 (0.004) | 0.32 (0.01) | 0.52 (0.01) | 0.38 (0.01) | 0.62 (0.01) | 0.47 | 0.78 |
| Impedance | 0.32 (0.01) | 0.37 (0.01) | 0.33 (0.01) | 0.32 (0.01) | 0.000 (0.003) | 0.10 (0.00) | 0.31 (0.01) | 0.14 (0.00) | 0.44 (0.01) | 0.20 | 0.63 |
| Neuroticism score | 0.13 (0.01) | 0.16 (0.01) | 0.14 (0.01) | 0.13 (0.00) | -0.000 (0.003) | 0.02 (0.00) | 0.14 (0.01) | 0.03 (0.00) | 0.23 (0.02) | 0.05 | 0.39 |
| Pulse rate | 0.15 (0.01) | 0.21 (0.01) | 0.18 (0.01) | 0.18 (0.01) | 0.000 (0.003) | 0.06 (0.00) | 0.32 (0.02) | 0.07 (0.00) | 0.42 (0.02) | 0.10 | 0.59 |
| Reaction time | 0.11 (0.01) | 0.12 (0.00) | 0.10 (0.00) | 0.11 (0.00) | -0.000 (0.003) | 0.01 (0.00) | 0.11 (0.01) | 0.02 (0.00) | 0.18 (0.02) | 0.03 | 0.31 |
| Systolic blood pressure | 0.18 (0.01) | 0.21 (0.01) | 0.19 (0.01) | 0.18 (0.01) | -0.001 (0.003) | 0.04 (0.00) | 0.21 (0.01) | 0.06 (0.00) | 0.31 (0.02) | 0.09 | 0.48 |
| College education | 0.31 (0.01) | 0.26 (0.01) | 0.23 (0.01) | 0.24 (0.01) | -0.000 (0.003) | 0.04 (0.00) | 0.18 (0.01) | 0.06 (0.00) | 0.27 (0.01) | 0.10 | 0.42 |
| Ever smoked | 0.13 (0.01) | 0.10 (0.00) | 0.09 (0.00) | 0.09 (0.00) | -0.001 (0.003) | 0.01 (0.00) | 0.11 (0.01) | 0.02 (0.00) | 0.17 (0.02) | 0.03 | 0.28 |
| Hypertension | 0.13 (0.01) | 0.14 (0.01) | 0.13 (0.00) | 0.12 (0.00) | 0.000 (0.003) | 0.02 (0.00) | 0.17 (0.02) | 0.03 (0.00) | 0.27 (0.02) | 0.05 | 0.44 |
| Snorer | 0.08 (0.01) | 0.08 (0.00) | 0.07 (0.00) | 0.07 (0.00) | -0.000 (0.003) | 0.01 (0.00) | 0.07 (0.01) | 0.01 (0.00) | 0.16 (0.02) | 0.02 | 0.31 |
| Difficulty sleeping | 0.07 (0.01) | 0.09 (0.00) | 0.08 (0.00) | 0.08 (0.00) | -0.000 (0.003) | 0.01 (0.00) | 0.07 (0.01) | 0.01 (0.00) | 0.13 (0.02) | 0.02 | 0.23 |
| Prefer evenings | 0.14 (0.01) | 0.16 (0.01) | 0.14 (0.00) | 0.15 (0.00) | -0.000 (0.003) | 0.02 (0.00) | 0.16 (0.01) | 0.03 (0.00) | 0.24 (0.02) | 0.05 | 0.38 |
| **Average** | **0.22 (0.00)** | **0.23 (0.00)** | **0.21 (0.00)** | **0.21 (0.00)** | **-0.000 (0.001)** | **0.06 (0.00)** | **0.21 (0.00)** | **0.08 (0.00)** | **0.31 (0.00)** | **0.11** | **0.46** |

**Supplementary Table 1: Heritabilities and best prediction models for the first 14 UK Biobank phenotypes.** For each phenotype, we report estimates of heritability ($h^2$), SNP heritability ($h^2_{\text{SNP}}$), inflation due to cryptic relatedness (see below) and the accuracies of the best prediction models (those computed from individual-level data using LDAK-Bolt-Predict assuming the BLD-LDAK Model). To estimate $h^2$, we first filtered the full UK Biobank dataset based on ancestry (we only kept individuals who were both recorded and inferred through principal component analysis to be white British), then identified 49 080 closely-related individuals (each individual has at least one first or second degree relative in the dataset); depending on phenotype, there were between 39 732 and 49 055 individuals. Finally, we performed Haseman Elston Regression,[19] using an unweighted kinship matrix computed from pruned SNPs. To estimate $h^2_{\text{SNP}}$ we used SumHer[3] assuming either the GCTA, LDAK-Thin or BLD-LDAK Model. When constructing prediction models, we varied $n$, the number of training samples from 100 000 to 200 000, with interval 10 000. Here we report $R^2$ and $R^2/h^2_{\text{SNP}}$ for $n$=100 000 and $n$=200 000, then predictions of $R^2$ and $R^2/h^2_{\text{SNP}}$ for $n$=400 000 (we obtained these predictions by regressing $R^2/h^2_{\text{SNP}}$ on $1 - \exp(a + bn)$).

We estimated inflation due to cryptic relatedness (population structure and familial relatedness) using our previously-described protocol [15,20,21] (described at www.ldak.org/quality-control). In brief, we used Haseman-Elston Regression to estimate SNP heritability from quarters of the genome (separately), then from the whole genome. If there is no inflation due to cryptic relatedness, then per-quarter estimates should sum to the whole-genome estimate. By contrast, if there is inflation, we would expect all five estimates to be inflated equally. Therefore, the total inflation can be estimated by subtracting the whole-genome estimate from the sum of the per-quarter estimates (and dividing by three).

| Phenotype | GCTA Model (K=1) $logl_{SS}$ | GCTA Model (K=1) AIC | LDAK-Thin Model (K=1) $logl_{SS}$ | LDAK-Thin Model (K=1) AIC | GCTA-LDMS-I Model (K=20) $logl_{SS}$ | GCTA-LDMS-I Model (K=20) AIC | BLD-LDAK Model (K=66) $logl_{SS}$ | BLD-LDAK Model (K=66) AIC | Baseline LD Model (K=75) $logl_{SS}$ | Baseline LD Model (K=75) AIC |
|---|---|---|---|---|---|---|---|---|---|---|
| Body mass index | 0 | 0 | 324.77 | -649.54 | 498.46 | -958.92 | 951.84 | -1773.68 | 812.13 | -1476.26 |
| Forced vital capacity | 0 | 0 | 233.89 | -467.78 | 511.08 | -984.16 | 1266.50 | -2403.00 | 1149.47 | -2150.94 |
| Height | 0 | 0 | 423.20 | -846.40 | 1269.00 | -2500.00 | 3670.30 | -7210.60 | 3301.80 | -6455.60 |
| Impedance | 0 | 0 | 341.63 | -683.26 | 611.28 | -1184.56 | 1263.05 | -2396.10 | 1132.82 | -2117.64 |
| Neuroticism score | 0 | 0 | 112.94 | -225.88 | 126.97 | -215.94 | 313.68 | -497.36 | 271.21 | -394.42 |
| Pulse rate | 0 | 0 | 178.01 | -356.02 | 263.30 | -488.60 | 756.91 | -1383.82 | 650.50 | -1153.00 |
| Reaction time | 0 | 0 | 86.68 | -173.36 | 71.28 | -104.56 | 213.18 | -296.36 | 189.25 | -230.50 |
| Systolic blood pressure | 0 | 0 | 185.48 | -370.96 | 238.99 | -439.98 | 593.50 | -1057.00 | 534.88 | -921.76 |
| College education | 0 | 0 | 187.51 | -375.02 | 191.80 | -345.60 | 498.15 | -866.30 | 428.70 | -709.40 |
| Ever smoked | 0 | 0 | 65.51 | -131.02 | 64.98 | -91.97 | 178.41 | -226.83 | 166.54 | -185.09 |
| Hypertension | 0 | 0 | 116.75 | -233.50 | 156.67 | -275.34 | 407.60 | -685.20 | 370.94 | -593.88 |
| Snorer | 0 | 0 | 42.06 | -84.12 | 45.51 | -53.02 | 139.11 | -148.21 | 119.27 | -90.53 |
| Difficulty sleeping | 0 | 0 | 67.41 | -134.81 | 64.62 | -91.24 | 175.14 | -220.28 | 157.21 | -166.42 |
| Prefer evenings | 0 | 0 | 120.77 | -241.54 | 140.84 | -243.68 | 317.50 | -505.00 | 279.76 | -411.52 |
| **Average** | **0** | **0** | **177.61** | **-355.23** | **303.91** | **-569.83** | **767.49** | **-1404.98** | **683.18** | **-1218.35** |

**Supplementary Table 2: Fit of heritability models for the first 14 UK Biobank phenotypes.** For each the first 14 UK Biobank phenotypes, we report $logl$, the approximate log likelihood computed by SumHer,[3,21] and the Akaike Information Criterion[22] (AIC), equal to $2K - 2logl$, where $K$ is the number of parameters in the heritability model (values are relative to those from the GCTA Model). We compare five heritability models (see Supplementary Table 6 for formal definitions): the GCTA Model (the most commonly-used model), the LDAK-Thin and BLD-LDAK Models (our two recommended models), and the GCTA-LDMS-I and Baseline LD Models (the models recommended by the authors of the softwares GCTA[23] and LD Score Regression,[13] respectively.) We see that the BLD-LDAK Model always results in best model fit (for each phenotype, the highest $logl$ and lowest AIC are marked in red). Note that this is a reduced version of the analysis in our previous publication,[21] where we used $logl$ to compare 12 heritability models.

| Phenotype | $R^2$ assuming GCTA Model Lasso | Ridge | Bolt | BayesR | $R^2$ assuming LDAK-Thin Model Lasso | Ridge | Bolt | BayesR | $R^2$ assuming BLD-LDAK Model Lasso | Ridge | Bolt | BayesR | Classical PRS $R^2$ |
|---|---|---|---|---|---|---|---|---|---|---|---|---|---|
| Body mass index | 0.091 | 0.098 | 0.107 | 0.107 | 0.099 | 0.107 | 0.115 | 0.115 | 0.100 | 0.111 | 0.117 | 0.117 | 0.074 |
| Forced vital capacity | 0.111 | 0.104 | 0.125 | 0.125 | 0.116 | 0.109 | 0.129 | 0.130 | 0.120 | 0.121 | 0.133 | 0.134 | 0.088 |
| Height | 0.367 | 0.297 | 0.363 | 0.370 | 0.366 | 0.313 | 0.370 | 0.381 | 0.371 | 0.342 | 0.382 | 0.384 | 0.237 |
| Impedance | 0.113 | 0.114 | 0.129 | 0.129 | 0.125 | 0.124 | 0.137 | 0.137 | 0.128 | 0.134 | 0.142 | 0.141 | 0.081 |
| Neuroticism score | 0.017 | 0.025 | 0.026 | 0.026 | 0.021 | 0.029 | 0.030 | 0.030 | 0.023 | 0.031 | 0.031 | 0.031 | 0.020 |
| Pulse rate | 0.062 | 0.046 | 0.069 | 0.069 | 0.067 | 0.052 | 0.071 | 0.072 | 0.068 | 0.059 | 0.075 | 0.075 | 0.047 |
| Reaction time | 0.008 | 0.015 | 0.016 | 0.016 | 0.011 | 0.017 | 0.017 | 0.018 | 0.012 | 0.019 | 0.019 | 0.019 | 0.015 |
| Systolic blood pressure | 0.043 | 0.040 | 0.050 | 0.050 | 0.045 | 0.043 | 0.053 | 0.053 | 0.047 | 0.049 | 0.057 | 0.056 | 0.029 |
| College education | 0.045 | 0.054 | 0.058 | 0.058 | 0.048 | 0.059 | 0.062 | 0.062 | 0.050 | 0.062 | 0.064 | 0.064 | 0.056 |
| Ever smoked | 0.006 | 0.012 | 0.013 | 0.013 | 0.009 | 0.014 | 0.015 | 0.015 | 0.009 | 0.015 | 0.016 | 0.016 | 0.011 |
| Hypertension | 0.023 | 0.022 | 0.029 | 0.029 | 0.024 | 0.024 | 0.030 | 0.031 | 0.027 | 0.028 | 0.032 | 0.033 | 0.018 |
| Snorer | 0.004 | 0.008 | 0.009 | 0.008 | 0.005 | 0.009 | 0.010 | 0.010 | 0.007 | 0.011 | 0.011 | 0.011 | 0.007 |
| Difficulty sleeping | 0.004 | 0.009 | 0.009 | 0.009 | 0.005 | 0.010 | 0.010 | 0.010 | 0.004 | 0.011 | 0.011 | 0.011 | 0.008 |
| Prefer evenings | 0.021 | 0.027 | 0.030 | 0.030 | 0.026 | 0.031 | 0.034 | 0.034 | 0.027 | 0.032 | 0.034 | 0.035 | 0.024 |
| **Average** | **0.065** | **0.062** | **0.074** | **0.074** | **0.069** | **0.067** | **0.077** | **0.078** | **0.071** | **0.073** | **0.080** | **0.081** | **0.051** |

**Supplementary Table 3: Accuracy of prediction models constructed using individual-level data.** For each of the 14 UK Biobank phenotypes, we construct models using LDAK-Lasso-Predict, LDAK-Ridge-Predict, LDAK-Bolt-Predict, LDAK-BayesR-Predict, assuming the GCTA, LDAK-Thin or BLD-LDAK Model, each time using individual-level data from all 200 000 training samples. We additionally construct classical PRS, again using data from all 200 000 training samples. Values report the accuracy of models, measured via $R^2$, the squared correlation between observed and predicted phenotypes across 20 000 test samples. The standard deviation of $R^2$ (estimated via jackknifing) is always less than 0.006. We see that the most accurate prediction models (marked in red) are produced by either LDAK-Bolt-Predict or LDAK-BayesR-Predict (given the similar performance of these two tools, we recommend LDAK-Bolt-Predict due to its faster runtime).

| LDAK-Lasso-SS | Sample | Number | UKBB | Estimated | $R^2$ (SD) | | | AUC (SD) | | |
|---|---|---|---|---|---|---|---|---|---|---|
| **Phenotype** | **Size** | **of SNPs** | **Cases** | $h^2_{SNP}$ **(SD)** | **GCTA** | **LDAK-Thin** | **BLD-LDAK** | **GCTA** | **LDAK-Thin** | **BLD-LDAK** |
| Asthma | 127669 | 190929 | 22316 | 0.08 (0.00) | 0.016 (0.001) | 0.016 (0.001) | 0.021 (0.001) | 0.582 (0.002) | 0.584 (0.002) | 0.595 (0.002) |
| Atrial fibrillation | 133073 | 545299 | 1701 | 0.07 (0.00) | 0.035 (0.005) | 0.042 (0.005) | 0.050 (0.006) | 0.621 (0.008) | 0.632 (0.008) | 0.640 (0.009) |
| Breast cancer | 214675 | 557359 | 5807 | 0.22 (0.00) | 0.052 (0.003) | 0.053 (0.003) | 0.058 (0.003) | 0.647 (0.004) | 0.649 (0.005) | 0.657 (0.004) |
| Inflammatory bowel disease | 34652 | 558975 | 1664 | 0.55 (0.02) | 0.043 (0.005) | 0.051 (0.005) | 0.071 (0.006) | 0.634 (0.008) | 0.646 (0.008) | 0.675 (0.007) |
| Prostate cancer | 140254 | 559023 | 2098 | 0.26 (0.01) | 0.097 (0.006) | 0.101 (0.006) | 0.102 (0.006) | 0.705 (0.007) | 0.708 (0.006) | 0.711 (0.006) |
| Rheumatoid arthritis | 58284 | 482013 | 2079 | 0.06 (0.01) | 0.038 (0.005) | 0.038 (0.005) | 0.042 (0.005) | 0.619 (0.008) | 0.619 (0.008) | 0.625 (0.008) |
| Schizophrenia | 76605 | 558526 | 195 | 0.59 (0.01) | 0.107 (0.020) | 0.115 (0.021) | 0.120 (0.022) | 0.719 (0.020) | 0.726 (0.020) | 0.729 (0.021) |
| Type 2 diabetes | 153002 | 558903 | 1246 | 0.08 (0.00) | 0.044 (0.006) | 0.053 (0.006) | 0.056 (0.006) | 0.637 (0.009) | 0.650 (0.009) | 0.653 (0.009) |
| **Average** | 117277 | 501378 | 4638 | 0.24 (0.00) | 0.054 (0.003) | 0.059 (0.003) | 0.065 (0.003) | 0.645 (0.003) | 0.652 (0.003) | 0.660 (0.003) |

| LDAK-Ridge-SS | Sample | Number | UKBB | Estimated | $R^2$ (SD) | | | AUC (SD) | | |
|---|---|---|---|---|---|---|---|---|---|---|
| **Phenotype** | **Size** | **of SNPs** | **Cases** | $h^2_{SNP}$ **(SD)** | **GCTA** | **LDAK-Thin** | **BLD-LDAK** | **GCTA** | **LDAK-Thin** | **BLD-LDAK** |
| Asthma | 127669 | 190929 | 22316 | 0.08 (0.00) | 0.014 (0.001) | 0.014 (0.001) | 0.019 (0.001) | 0.577 (0.002) | 0.578 (0.002) | 0.589 (0.002) |
| Atrial fibrillation | 133073 | 545299 | 1701 | 0.07 (0.00) | 0.019 (0.003) | 0.022 (0.004) | 0.030 (0.004) | 0.592 (0.008) | 0.600 (0.008) | 0.614 (0.009) |
| Breast cancer | 214675 | 557359 | 5807 | 0.22 (0.00) | 0.037 (0.003) | 0.040 (0.003) | 0.047 (0.003) | 0.624 (0.004) | 0.629 (0.004) | 0.640 (0.004) |
| Inflammatory bowel disease | 34652 | 558975 | 1664 | 0.55 (0.02) | 0.035 (0.004) | 0.039 (0.005) | 0.061 (0.005) | 0.620 (0.008) | 0.627 (0.008) | 0.660 (0.008) |
| Prostate cancer | 140254 | 559023 | 2098 | 0.26 (0.01) | 0.066 (0.005) | 0.071 (0.005) | 0.079 (0.006) | 0.668 (0.007) | 0.675 (0.007) | 0.684 (0.007) |
| Rheumatoid arthritis | 58284 | 482013 | 2079 | 0.06 (0.01) | 0.036 (0.004) | 0.037 (0.004) | 0.037 (0.004) | 0.617 (0.007) | 0.618 (0.007) | 0.616 (0.008) |
| Schizophrenia | 76605 | 558526 | 195 | 0.59 (0.01) | 0.103 (0.020) | 0.111 (0.021) | 0.116 (0.022) | 0.715 (0.021) | 0.722 (0.020) | 0.726 (0.021) |
| Type 2 diabetes | 153002 | 558903 | 1246 | 0.08 (0.00) | 0.036 (0.005) | 0.043 (0.006) | 0.047 (0.006) | 0.624 (0.009) | 0.636 (0.009) | 0.639 (0.009) |
| **Average** | 117277 | 501378 | 4638 | 0.24 (0.00) | 0.043 (0.003) | 0.047 (0.003) | 0.054 (0.003) | 0.630 (0.003) | 0.636 (0.003) | 0.646 (0.004) |

| LDAK-Bolt-SS | Sample | Number | UKBB | Estimated | $R^2$ (SD) | | | AUC (SD) | | |
|---|---|---|---|---|---|---|---|---|---|---|
| **Phenotype** | **Size** | **of SNPs** | **Cases** | $h^2_{SNP}$ **(SD)** | **GCTA** | **LDAK-Thin** | **BLD-LDAK** | **GCTA** | **LDAK-Thin** | **BLD-LDAK** |
| Asthma | 127669 | 190929 | 22316 | 0.08 (0.00) | 0.014 (0.001) | 0.014 (0.001) | 0.019 (0.001) | 0.577 (0.002) | 0.578 (0.002) | 0.589 (0.002) |
| Atrial fibrillation | 133073 | 545299 | 1701 | 0.07 (0.00) | 0.019 (0.003) | 0.022 (0.004) | 0.030 (0.004) | 0.592 (0.008) | 0.600 (0.008) | 0.614 (0.009) |
| Breast cancer | 214675 | 557359 | 5807 | 0.22 (0.00) | 0.037 (0.003) | 0.040 (0.003) | 0.047 (0.003) | 0.624 (0.004) | 0.629 (0.004) | 0.640 (0.004) |
| Inflammatory bowel disease | 34652 | 558975 | 1664 | 0.55 (0.02) | 0.035 (0.004) | 0.039 (0.005) | 0.061 (0.005) | 0.620 (0.008) | 0.627 (0.008) | 0.660 (0.008) |
| Prostate cancer | 140254 | 559023 | 2098 | 0.26 (0.01) | 0.066 (0.005) | 0.071 (0.005) | 0.079 (0.006) | 0.668 (0.007) | 0.675 (0.007) | 0.684 (0.007) |
| Rheumatoid arthritis | 58284 | 482013 | 2079 | 0.06 (0.01) | 0.036 (0.004) | 0.037 (0.004) | 0.037 (0.004) | 0.617 (0.007) | 0.618 (0.007) | 0.616 (0.008) |
| Schizophrenia | 76605 | 558526 | 195 | 0.59 (0.01) | 0.103 (0.020) | 0.111 (0.021) | 0.116 (0.022) | 0.715 (0.021) | 0.722 (0.020) | 0.726 (0.021) |
| Type 2 diabetes | 153002 | 558903 | 1246 | 0.08 (0.00) | 0.036 (0.005) | 0.043 (0.006) | 0.047 (0.006) | 0.624 (0.009) | 0.636 (0.009) | 0.639 (0.009) |
| **Average** | 117277 | 501378 | 4638 | 0.24 (0.00) | 0.043 (0.003) | 0.047 (0.003) | 0.054 (0.003) | 0.630 (0.003) | 0.636 (0.003) | 0.646 (0.004) |

| LDAK-BayesR-SS | Sample | Number | UKBB | Estimated | $R^2$ (SD) | | | AUC (SD) | | |
|---|---|---|---|---|---|---|---|---|---|---|
| **Phenotype** | **Size** | **of SNPs** | **Cases** | $h^2_{SNP}$ **(SD)** | **GCTA** | **LDAK-Thin** | **BLD-LDAK** | **GCTA** | **LDAK-Thin** | **BLD-LDAK** |
| Asthma | 127669 | 190929 | 22316 | 0.08 (0.00) | 0.019 (0.001) | 0.019 (0.001) | 0.022 (0.001) | 0.590 (0.002) | 0.592 (0.002) | 0.597 (0.002) |
| Atrial fibrillation | 133073 | 545299 | 1701 | 0.07 (0.00) | 0.059 (0.006) | 0.064 (0.006) | 0.064 (0.006) | 0.654 (0.008) | 0.660 (0.008) | 0.659 (0.008) |
| Breast cancer | 214675 | 557359 | 5807 | 0.22 (0.00) | 0.061 (0.003) | 0.063 (0.003) | 0.064 (0.003) | 0.660 (0.004) | 0.662 (0.004) | 0.664 (0.004) |
| Inflammatory bowel disease | 34652 | 558975 | 1664 | 0.55 (0.02) | 0.070 (0.006) | 0.075 (0.006) | 0.081 (0.006) | 0.673 (0.007) | 0.681 (0.007) | 0.688 (0.007) |
| Prostate cancer | 140254 | 559023 | 2098 | 0.26 (0.01) | 0.113 (0.006) | 0.113 (0.006) | 0.114 (0.006) | 0.724 (0.006) | 0.723 (0.006) | 0.724 (0.006) |
| Rheumatoid arthritis | 58284 | 482013 | 2079 | 0.06 (0.01) | 0.040 (0.005) | 0.041 (0.005) | 0.042 (0.005) | 0.625 (0.008) | 0.626 (0.008) | 0.626 (0.008) |
| Schizophrenia | 76605 | 558526 | 195 | 0.59 (0.01) | 0.112 (0.020) | 0.117 (0.021) | 0.116 (0.022) | 0.725 (0.020) | 0.727 (0.020) | 0.726 (0.021) |
| Type 2 diabetes | 153002 | 558903 | 1246 | 0.08 (0.00) | 0.045 (0.006) | 0.055 (0.006) | 0.056 (0.006) | 0.639 (0.009) | 0.653 (0.009) | 0.657 (0.009) |
| **Average** | 117277 | 501378 | 4638 | 0.24 (0.00) | 0.065 (0.003) | 0.068 (0.003) | 0.070 (0.003) | 0.661 (0.003) | 0.666 (0.003) | 0.668 (0.003) |

**Supplementary Table 4: Prediction of eight diseases using results from external association studies.** We use summary statistics for asthma from the study of Demenais *et al.*,[24] atrial fibrillation from Christophersen *et al.*,[25] breast cancer from Zhang *et al.*,[26] inflammatory bowel disease from Liu *et al.*,[27] prostate cancer from Schumacher *et al.*,[28] rheumatoid arthritis from Okada *et al.*,[29] schizophrenia from the Psychiatric Genomics Consortium,[30] and type 2 diabetes from Scott *et al.*[31] For each disease, we report the size of the association study, the number of SNPs for which we have summary statistics (after excluding those with ambiguous alleles or not present in our UK Biobank dataset), the number of cases in the UK Biobank, and estimates of SNP heritability $h^2_{SNP}$, obtained using SumHer[3] assuming the BLD-LDAK Model. We construct prediction models using LDAK-Lasso-SS (top table), LDAK-Ridge-SS (second table), LDAK-Bolt-SS (third table) and LDAK-BayesR-SS (bottom table), assuming the GCTA, LDAK-Thin and BLD-LDAK Models, then measure the accuracy of these models via $R^2$, the squared correlation between observed and predicted phenotypes, and AUC, the area under the receiver operating curve (we calculate $R^2$ and AUC using all the UK Biobank cases, plus three times as many controls). For each disease and each prediction tool, the heritability model leading to highest accuracy is marked in red. We see that for all diseases, both $R^2$ and AUC improve when we replace the GCTA Model with either the LDAK-Thin or BLD-LDAK Model.

| **South Asian ancestry** | **$R^2$ assuming GCTA Model** | | | | **$R^2$ assuming LDAK-Thin Model** | | | | **$R^2$ assuming BLD-LDAK Model** | | | | **Classical PRS** |
|---|---|---|---|---|---|---|---|---|---|---|---|---|---|
| **Phenotype** | **Lasso** | **Ridge** | **Bolt** | **BayesR** | **Lasso** | **Ridge** | **Bolt** | **BayesR** | **Lasso** | **Ridge** | **Bolt** | **BayesR** | **$R^2$** |
| Body mass index | 0.000 | 0.058 | 0.065 | 0.065 | 0.000 | 0.065 | 0.072 | 0.072 | 0.000 | 0.074 | 0.078 | 0.077 | 0.041 |
| Forced vital capacity | 0.023 | 0.038 | 0.047 | 0.047 | 0.023 | 0.047 | 0.053 | 0.052 | 0.026 | 0.053 | 0.058 | 0.058 | 0.034 |
| Height | 0.206 | 0.154 | 0.201 | 0.205 | 0.208 | 0.171 | 0.211 | 0.215 | 0.217 | 0.195 | 0.223 | 0.226 | 0.127 |
| Impedance | 0.000 | 0.080 | 0.096 | 0.096 | 0.000 | 0.092 | 0.105 | 0.105 | 0.000 | 0.105 | 0.112 | 0.110 | 0.059 |
| Neuroticism score | 0.000 | 0.015 | 0.013 | 0.014 | 0.000 | 0.016 | 0.016 | 0.015 | 0.000 | 0.018 | 0.018 | 0.018 | 0.012 |
| Pulse rate | 0.000 | 0.033 | 0.048 | 0.048 | 0.000 | 0.037 | 0.050 | 0.051 | 0.000 | 0.037 | 0.050 | 0.052 | 0.038 |
| Reaction time | 0.000 | 0.006 | 0.006 | 0.006 | 0.000 | 0.007 | 0.007 | 0.007 | 0.000 | 0.009 | 0.009 | 0.008 | 0.005 |
| Systolic blood pressure | 0.000 | 0.020 | 0.027 | 0.027 | 0.000 | 0.022 | 0.028 | 0.027 | 0.000 | 0.029 | 0.032 | 0.032 | 0.013 |
| College education | 0.000 | 0.017 | 0.017 | 0.018 | 0.000 | 0.021 | 0.022 | 0.022 | 0.000 | 0.022 | 0.023 | 0.023 | 0.018 |
| Ever smoked | 0.000 | 0.006 | 0.006 | 0.006 | 0.000 | 0.008 | 0.008 | 0.008 | 0.000 | 0.009 | 0.008 | 0.008 | 0.006 |
| Hypertension | 0.000 | 0.019 | 0.024 | 0.024 | 0.000 | 0.020 | 0.023 | 0.023 | 0.000 | 0.024 | 0.026 | 0.027 | 0.014 |
| Snorer | 0.000 | 0.001 | 0.001 | 0.001 | 0.000 | 0.001 | 0.001 | 0.001 | 0.000 | 0.000 | 0.000 | 0.000 | 0.000 |
| Difficulty sleeping | 0.000 | 0.004 | 0.004 | 0.004 | 0.000 | 0.005 | 0.005 | 0.005 | 0.000 | 0.004 | 0.004 | 0.004 | 0.004 |
| Prefer evenings | 0.000 | 0.001 | 0.002 | 0.002 | 0.000 | 0.002 | 0.002 | 0.002 | 0.000 | 0.002 | 0.002 | 0.002 | 0.001 |
| **Average** | **0.016** | **0.032** | **0.040** | **0.040** | **0.017** | **0.037** | **0.043** | **0.043** | **0.017** | **0.041** | **0.046** | **0.046** | **0.027** |

| **African ancestry** | **$R^2$ assuming GCTA Model** | | | | **$R^2$ assuming LDAK-Thin Model** | | | | **$R^2$ assuming BLD-LDAK Model** | | | | **Classical PRS** |
|---|---|---|---|---|---|---|---|---|---|---|---|---|---|
| **Phenotype** | **Lasso** | **Ridge** | **Bolt** | **BayesR** | **Lasso** | **Ridge** | **Bolt** | **BayesR** | **Lasso** | **Ridge** | **Bolt** | **BayesR** | **$R^2$** |
| Body mass index | 0.000 | 0.014 | 0.018 | 0.019 | 0.000 | 0.024 | 0.027 | 0.026 | 0.000 | 0.021 | 0.023 | 0.021 | 0.008 |
| Forced vital capacity | 0.008 | 0.011 | 0.014 | 0.014 | 0.008 | 0.010 | 0.016 | 0.017 | 0.010 | 0.010 | 0.015 | 0.015 | 0.008 |
| Height | 0.053 | 0.028 | 0.050 | 0.049 | 0.057 | 0.039 | 0.053 | 0.053 | 0.064 | 0.048 | 0.065 | 0.058 | 0.021 |
| Impedance | 0.000 | 0.014 | 0.021 | 0.022 | 0.000 | 0.022 | 0.026 | 0.027 | 0.000 | 0.024 | 0.028 | 0.030 | 0.012 |
| Neuroticism score | 0.001 | 0.000 | 0.000 | 0.000 | 0.001 | 0.001 | 0.001 | 0.001 | 0.001 | 0.001 | 0.000 | 0.000 | 0.000 |
| Pulse rate | 0.000 | 0.011 | 0.008 | 0.008 | 0.000 | 0.013 | 0.007 | 0.006 | 0.000 | 0.013 | 0.011 | 0.012 | 0.010 |
| Reaction time | 0.000 | 0.001 | 0.000 | 0.000 | 0.001 | 0.001 | 0.001 | 0.000 | 0.001 | 0.001 | 0.000 | 0.000 | 0.001 |
| Systolic blood pressure | 0.000 | 0.006 | 0.007 | 0.007 | 0.001 | 0.007 | 0.006 | 0.005 | 0.001 | 0.011 | 0.009 | 0.008 | 0.003 |
| College education | 0.000 | 0.002 | 0.002 | 0.002 | 0.000 | 0.001 | 0.001 | 0.001 | 0.000 | 0.001 | 0.001 | 0.001 | 0.005 |
| Ever smoked | 0.000 | 0.000 | 0.000 | 0.000 | 0.000 | 0.000 | 0.000 | 0.000 | 0.000 | 0.000 | 0.000 | 0.000 | 0.000 |
| Hypertension | 0.000 | 0.004 | 0.007 | 0.008 | 0.000 | 0.005 | 0.008 | 0.008 | 0.000 | 0.005 | 0.008 | 0.007 | 0.002 |
| Snorer | 0.000 | 0.000 | 0.000 | 0.000 | 0.000 | 0.000 | 0.000 | 0.000 | 0.000 | 0.000 | 0.000 | 0.000 | 0.000 |
| Difficulty sleeping | 0.000 | 0.000 | 0.000 | 0.000 | 0.000 | 0.000 | 0.000 | 0.000 | 0.000 | 0.000 | 0.000 | 0.000 | 0.001 |
| Prefer evenings | 0.001 | 0.001 | 0.002 | 0.002 | 0.000 | 0.001 | 0.003 | 0.003 | 0.000 | 0.001 | 0.002 | 0.002 | 0.002 |
| **Average** | **0.005** | **0.007** | **0.009** | **0.009** | **0.005** | **0.009** | **0.011** | **0.010** | **0.006** | **0.010** | **0.012** | **0.011** | **0.005** |

| **East Asian ancestry** | **$R^2$ assuming GCTA Model** | | | | **$R^2$ assuming LDAK-Thin Model** | | | | **$R^2$ assuming BLD-LDAK Model** | | | | **Classical PRS** |
|---|---|---|---|---|---|---|---|---|---|---|---|---|---|
| **Phenotype** | **Lasso** | **Ridge** | **Bolt** | **BayesR** | **Lasso** | **Ridge** | **Bolt** | **BayesR** | **Lasso** | **Ridge** | **Bolt** | **BayesR** | **$R^2$** |
| Body mass index | 0.001 | 0.044 | 0.049 | 0.049 | 0.000 | 0.049 | 0.051 | 0.051 | 0.003 | 0.047 | 0.049 | 0.050 | 0.036 |
| Forced vital capacity | 0.018 | 0.034 | 0.049 | 0.049 | 0.017 | 0.044 | 0.053 | 0.054 | 0.022 | 0.056 | 0.061 | 0.059 | 0.028 |
| Height | 0.150 | 0.109 | 0.154 | 0.153 | 0.144 | 0.119 | 0.156 | 0.159 | 0.143 | 0.133 | 0.157 | 0.159 | 0.086 |
| Impedance | 0.003 | 0.036 | 0.044 | 0.048 | 0.001 | 0.043 | 0.053 | 0.055 | 0.003 | 0.045 | 0.049 | 0.054 | 0.037 |
| Neuroticism score | 0.000 | 0.003 | 0.003 | 0.003 | 0.000 | 0.003 | 0.007 | 0.007 | 0.000 | 0.006 | 0.008 | 0.008 | 0.002 |
| Pulse rate | 0.000 | 0.031 | 0.053 | 0.052 | 0.000 | 0.036 | 0.058 | 0.058 | 0.000 | 0.035 | 0.057 | 0.060 | 0.026 |
| Reaction time | 0.000 | 0.006 | 0.005 | 0.005 | 0.000 | 0.005 | 0.005 | 0.005 | 0.000 | 0.008 | 0.007 | 0.006 | 0.005 |
| Systolic blood pressure | 0.000 | 0.006 | 0.021 | 0.017 | 0.000 | 0.011 | 0.026 | 0.023 | 0.000 | 0.012 | 0.022 | 0.021 | 0.008 |
| College education | 0.001 | 0.003 | 0.005 | 0.005 | 0.001 | 0.004 | 0.005 | 0.005 | 0.002 | 0.004 | 0.005 | 0.005 | 0.005 |
| Ever smoked | 0.001 | 0.001 | 0.001 | 0.001 | 0.001 | 0.002 | 0.002 | 0.002 | 0.001 | 0.003 | 0.003 | 0.003 | 0.001 |
| Hypertension | 0.000 | 0.008 | 0.017 | 0.020 | 0.001 | 0.009 | 0.016 | 0.018 | 0.000 | 0.012 | 0.019 | 0.018 | 0.014 |
| Snorer | 0.000 | 0.001 | 0.002 | 0.001 | 0.000 | 0.002 | 0.002 | 0.002 | 0.000 | 0.002 | 0.003 | 0.002 | 0.001 |
| Difficulty sleeping | 0.000 | 0.000 | 0.000 | 0.000 | 0.001 | 0.000 | 0.000 | 0.000 | 0.000 | 0.001 | 0.001 | 0.001 | 0.000 |
| Prefer evenings | 0.004 | 0.001 | 0.001 | 0.001 | 0.003 | 0.004 | 0.003 | 0.003 | 0.002 | 0.003 | 0.002 | 0.003 | 0.002 |
| **Average** | **0.013** | **0.020** | **0.029** | **0.029** | **0.012** | **0.024** | **0.031** | **0.032** | **0.013** | **0.026** | **0.032** | **0.032** | **0.018** |

**Supplementary Table 5: Cross-ancestry prediction.** This table is the same as Supplementary Table 3, except now $R^2$ is measured using individuals of South Asian (top), African (middle) or East Asian (bottom) ancestry. The number of individuals depends on the population and phenotype: South Asian, average 6 710 (range 4 370 - 7 057); African, average 2 588 (range 1 716 - 2 717); East Asian, average 1 273 (range 855 - 1 331). The best-performing tool for each phenotype is marked in red.

| **Model** | $K$ | **Defintion** | **Notes (further details provided in caption)** |
|---|---|---|---|
| GCTA | 1 | $\mathbb{E}[h_j^2] = \tau_1$ | |
| LDAK-Thin | 1 | $\mathbb{E}[h_j^2] = I_j[f_j(1-f_j)]^{0.75}\,\tau_1$ | $I_j$ indicates whether SNP $j$ remains after removing duplicates, $f_j$ is its MAF |
| GCTA-LDMS-I | 20 | $\mathbb{E}[h_j^2] = \sum_{k=1}^{20} I_{jk}\,\tau_k$ | $I_{jk}$ indicates whether SNP j belongs to LD-MAF Bin $k$ |
| Baseline LD | 75 | $\mathbb{E}[h_j^2] = \tau_1 + \sum_{k=2}^{75} c_{j(k-1)}\,\tau_k$ | Contains 74 annotations, listed in Supplementary Table 7 |
| BLD-LDAK | 66 | $\mathbb{E}[h_j^2] = [f_j(1-f_j)]^{0.75}(\tau_1 + \sum_{k=2}^{7} c_{j(k-1)}\,\tau_k$ | Contains 64 annotations from Baseline LD Model; |
| | | $+ \sum_{k=8}^{65} c_{j(k+9)}\,\tau_k + w_j\,\tau_{66})$ | $w_j$ is the LDAK weighting of SNP $j$, while $f_j$ is its MAF |

**Supplementary Table 6: Definitions of heritability models.** $K$ is the number of parameters. The GCTA Model[23] assumes $\mathbb{E}[h_j^2]$ is constant across the genome. The LDAK-Thin Model[21] first removes duplicate SNPs ($I_j$ indicates whether SNP $j$ remains after thinning so that no pair remains within 100 kb with $r_{jl}^2 > 0.98$), then for SNPs that remain, it assumes $\mathbb{E}[h_j^2]$ is proportional to $[f_j(1-f_j)]^{0.75}$, where $f_j$ is the MAF of SNP $j$. The GCTA-LDMS-I Model[10] partitions SNPs based on LD and MAF ($I_{j,k}$ indicates whether SNP $j$ belongs to Bin $k$), then allows $\mathbb{E}[h_j^2]$ to vary across bins. We divided SNPs four-ways based on LD score quartiles, then five-ways using the MAF boundaries 0.1, 0.2, 0.3 & 0.4, resulting in 20 bins.[21] The Baseline LD Model[11] uses 74 SNP annotations ($c_{j1}, c_{j2}, \ldots, c_{j74}$), which are described in Supplementary Table 7. The BLD-LDAK Model[21] uses 64 of these annotations, plus $w_j$, the LDAK weighting[32] of SNP $j$ (SNPs in regions of high linkage disequilibrium tend to get lower $w_j$, and vice versa), in addition to assuming that $\mathbb{E}[h_j^2]$ varies with $[f_j(1-f_j)]^{0.75}$.

| | | Annotation | % SNPs | | Annotation | % SNPs |
|---|---|---|---|---|---|---|
| LD-Rel. | 1 | MAF_Adj_Predicted_Allele_Age | NA | 2 | MAF_Adj_LLD_AFR | NA |
| | 3 | Recomb_Rate_10kb | NA | 4 | Nucleotide_Diversity_10kb | NA |
| | 5 | Backgrd_Selection_Stat | NA | 6 | CpG_Content_50kb | NA |
| MAF Indicators | 7 | MAFbin1 ($0.05 < \text{MAF} \leq 0.07$) | 6.1 | 8 | MAFbin2 ($0.07 < \text{MAF} \leq 0.10$) | 6.0 |
| | 9 | MAFbin3 ($0.10 < \text{MAF} \leq 0.13$) | 5.9 | 10 | MAFbin4 ($0.13 < \text{MAF} \leq 0.17$) | 6.0 |
| | 11 | MAFbin5 ($0.17 < \text{MAF} \leq 0.21$) | 5.9 | 12 | MAFbin6 ($0.21 < \text{MAF} \leq 0.26$) | 5.9 |
| | 13 | MAFbin7 ($0.26 < \text{MAF} \leq 0.32$) | 5.9 | 14 | MAFbin8 ($0.32 < \text{MAF} \leq 0.38$) | 6.0 |
| | 15 | MAFbin9 ($0.38 < \text{MAF} \leq 0.44$) | 6.0 | 16 | MAFbin10 ($0.44 < \text{MAF} \leq 0.50$) | 5.9 |
| Function Indicators | 17 | Coding_UCSC | 1.6 | 18 | Conserved_LindbladToh | 2.9 |
| | 19 | CTCF_Hoffman | 2.4 | 20 | DGF_ENCODE | 13 |
| | 21 | DHS_Trynka | 16 | 22 | Enhancer_Andersson | 0.4 |
| | 23 | Enhancer_Hoffman | 4.3 | 24 | FetalDHS_Trynka | 8.6 |
| | 25 | H3K27ac_Hnisz | 39 | 26 | H3K27ac_PGC2 | 27 |
| | 27 | H3K4me1_Trynka | 42 | 28 | H3K4me3_Trynka | 13 |
| | 29 | H3K9ac_Trynka | 12 | 30 | Intron_UCSC | 39 |
| | 31 | PromoterFlanking_Hoffman | 0.9 | 32 | Promoter_UCSC | 4.8 |
| | 33 | Repressed_Hoffman | 45 | 34 | SuperEnhancer_Hnisz | 16 |
| | 35 | TFBS_ENCODE | 13 | 36 | Transcr_Hoffman | 35 |
| | 37 | TSS_Hoffman | 1.9 | 38 | UTR_3_UCSC | 1.2 |
| | 39 | UTR_5_UCSC | 0.6 | 40 | WeakEnhancer_Hoffman | 2.1 |
| Other Annotations | 41 | Coding_UCSC.extend.500 | 6.7 | 42 | Conserved_LindbladToh.extend.500 | 33 |
| | 43 | CTCF_Hoffman.extend.500 | 7.1 | 44 | DGF_ENCODE.extend.500 | 54 |
| | 45 | DHS_Trynka.extend.500 | 49 | 46 | Enhancer_Andersson.extend.500 | 1.9 |
| | 47 | Enhancer_Hoffman.extend.500 | 9.1 | 48 | FetalDHS_Trynka.extend.500 | 28 |
| | 49 | H3K27ac_Hnisz.extend.500 | 42 | 50 | H3K27ac_PGC2.extend.500 | 33 |
| | 51 | H3K4me1_Trynka.extend.500 | 60 | 52 | H3K4me3_Trynka.extend.500 | 25 |
| | 53 | H3K9ac_Trynka.extend.500 | 23 | 54 | Intron_UCSC.extend.500 | 40 |
| | 55 | PromoterFlanking_Hoffman.extend.500 | 3.4 | 56 | Promoter_UCSC.extend.500 | 5.9 |
| | 57 | Repressed_Hoffman.extend.500 | 70 | 58 | SuperEnhancer_Hnisz.extend.500 | 17 |
| | 59 | TFBS_ENCODE.extend.500 | 34 | 60 | Transcr_Hoffman.extend.500 | 76 |
| | 61 | TSS_Hoffman.extend.500 | 3.6 | 62 | UTR_3_UCSC.extend.500 | 2.8 |
| | 63 | UTR_5_UCSC.extend.500 | 2.8 | 64 | WeakEnhancer_Hoffman.extend.500 | 8.9 |
| | 65 | DHS_peaks_Trynka | 11 | 66 | H3K4me1_peaks_Trynka | 17 |
| | 67 | H3K4me3_peaks_Trynka | 4.3 | 68 | H3K9ac_peaks_Trynka | 4.0 |
| | 69 | Super_Enhancer_Vahedi | 2.2 | 70 | Super_Enhancer_Vahedi.extend.500 | 2.2 |
| | 71 | Typical_Enhancer_Vahedi | 2.2 | 72 | Typical_Enhancer_Vahedi.extend.500 | 2.7 |
| | 73 | GERP.NS | NA | 74 | GERP.RSsup4 | 0.9 |

**Supplementary Table 7: Baseline LD Model SNP annotations.** This table lists the 74 annotations of the Baseline LD Model.[11] These can be divided into 6 LD-related annotations, 10 MAF indicators, 24 function indicators and 34 auxiliary annotations (predominantly indicators of buffer regions for the functional categories). The names are those provided in the annotation files on the LDSC website (www.github.com/bulik/ldsc). For binary annotations, we report the percentage of SNPs with value 1 (i.e., the proportion of SNPs within the corresponding category). For more details of each annotation, see the correspondence of Gazal et al.[33] and earlier publications.[11,34] The BLD-LDAK Model uses 64 of these annotations, excluding the MAF bins (Annotations 7-16).

| Tool | Prior distribution for $\beta_j$ | Parameter choices |
|---|---|---|
| LDAK-Lasso-SS<br>11 Models | $DE(\lambda/\mathbb{E}[h_j^2]^{\frac{1}{2}})$ | $2/\lambda^2 \in \{0.5, 0.6, \ldots, 1.4, 1.5\}$ |
| LDAK-Lasso-Sparse-SS<br>80 Models | $DE(\lambda/\mathbb{E}[h_j^2]^{\frac{1}{2}})$ | $\lambda \in n/1000 \times 100^{\frac{i}{19}} \times \overline{\mathbb{E}[h_j^2]}^{\frac{1}{2}}$, for $i \in \{0, 1, \ldots, 19\}$<br>$t \in \{0.9, 0.5, 0.2, 0.1\}$ ($t$ explained in caption) |
| LDAK-Ridge-SS<br>11 Models | $\mathbb{N}(0, v\mathbb{E}[h_j^2])$ | $v \in \{0.5, 0.6, \ldots, 1.4, 1.5\}$ |
| LDAK-Bolt-SS<br>132 Models | $p\,\mathbb{N}(0, (1\text{-}f_2)/p\,\mathbb{E}[h_j^2]) + (1\text{-}p)\,\mathbb{N}(0, f_2/(1\text{-}p)\,\mathbb{E}[h_j^2])$ | $p \in \{0.01, 0.02, 0.05, 0.1, 0.15, 0.2, 0.25, 0.3, 0.35, 0.4, 0.45, 0.5\}$<br>$f_2 \in \{0, 0.05, 0.1, 0.15, 0.2, 0.25, 0.3, 0.35, 0.4, 0.45, 0.5\}$ |
| LDAK-Bolt-Sparse-SS<br>22 Models | $p\,\mathbb{N}(0, 1/p\,\mathbb{E}[h_j^2]) + (1\text{-}p)\,\delta_{\{0\}}$ | $p \in \{0.01, 0.02, 0.05, 0.1, 0.15, 0.2, 0.25, \ldots, ..0.95, 1\}$ |
| LDAK-BayesR-SS<br>84 Models | $\pi_1\,\delta_{\{0\}} + \pi_2\,\mathbb{N}(0, s\mathbb{E}[h_j^2]/100) + \pi_3\,\mathbb{N}(0, s\mathbb{E}[h_j^2]/10)$<br>$+\pi_4\,\mathbb{N}(0, s\mathbb{E}[h_j^2])$ where $\pi_1 + \pi_2 + \pi_3 + \pi_4 = 1$<br>and $s = (\pi_2/100 + \pi_3/10 + \pi_4)^{-1}$ | EITHER $(\pi_1, \pi_2, \pi_3, \pi_4) = (1, 0, 0, 0)$<br>OR $(\pi_2, \pi_3, \pi_4) \in \{0, 0.005, 0.01, 0.02, 0.05, 0.1, 0.2\}^3$<br>with $\pi_2 + \pi_3 + \pi_4 > 0$ and $\pi_2 \geq \pi_3 \geq \pi_4$ |

**Supplementary Table 8: Prior distribution parameter choices.** Each of our summary statistic prediction tools constructs pairs of training and full prediction models. Here we detail the prior distribution parameters to which the pairs of models correspond. In addition to LDAK-Lasso-SS, LDAK-Ridge-SS, LDAK-Bolt-SS and LDAK-BayesR-SS (our four main summary statistic tools), we describe LDAK-Lasso-Sparse-SS and LDAK-Bolt-Sparse-SS, two alternative tools that we use in Extended Data Fig. 1. $DE(a)$ denotes a double exponential distribution with rate $a$, $\mathbb{N}(b, c)$ denotes a Gaussian distribution with mean $b$ and variance $c$, while $\delta_{\{0\}}$ denotes a point mass at zero. All tools assume the linear model $\mathbb{E}[Y] = X_1\beta_1 + X_2\beta_2 + \ldots + X_m\beta_m$, where $Y$ is the phenotype, while $X_j$ and $\beta_j$ denote the genotypes and effect size for SNP $j$, respectively. We assume that $X_j$ and $Y$ have been standardized to have mean zero and variance one, and therefore, the expected heritabilty contributed by SNP $j$ is $\mathbb{E}[\beta_j^2]$. The prior distributions used by LDAK-Bolt-SS, LDAK-Bolt-Sparse-SS and LDAK-BayesR-SS ensure that $\mathbb{E}[\beta_j^2]$ equals $\mathbb{E}[h_j^2]$, while the prior distributions used by LDAK-Lasso-SS, LDAK-Lasso-Sparse-SS and LDAK-Ridge-SS ensure that $\mathbb{E}[\beta_j^2]$ is proportional to $\mathbb{E}[h_j^2]$.

Although LDAK-Lasso-SS uses the same form for the effect size prior distribution as the existing tool lassosum[7], the two tools have quite different algorithms. Therefore, we also developed LDAK-Lasso-Sparse-SS, which can be considered a generalized version of lassosum.[7] Like lassosum, LDAK-Lasso-Sparse-SS uses coordinate descent (each effect size estimate is replaced by its conditional posterior mode). Also like lassosum, LDAK-Lasso-Sparse-SS introduces a shrinkage parameter $t$, which determines how much to scale SNP-SNP correlations. The authors of lassosum explain that scaling the correlations introduces an element of shrinkage, making their tool similar to the elastic net.[35]. We consider the same four values for $t$ as lassosum, while our choices for $\lambda$ ensure that when LDAK-Lasso-Sparse-SS is run assuming the GCTA Model (in which case $\mathbb{E}[h_j^2] = \overline{\mathbb{E}[h_j^2]}$), the corresponding values for the rate parameter of the double exponential distribution match those used by lassosum.

In general, it is substantially faster (and requires less memory) to construct prediction models using summary statistics than using individual-level data, and as a result, it is feasible to consider more parameter values. This is why LDAK-Ridge-SS considers 11 values for $v$, whereas LDAK-Ridge-Predict fixes $v = 1$, why LDAK-Bolt-SS considers 132 values for $(p, f_2)$, whereas LDAK-Bolt-Predict considers only 18, and why LDAK-BayesR-SS considers 84 values for $(\pi_1, \pi_2, \pi_3, \pi_4)$, whereas LDAK-BayesR-Predict considers only 35.
